# Capture of the SmSTRIPAK proxiome identifies the greenbeard proteins SmDOC1/2 as regulators of the MAK2 pathway to control sexual development in *Sordaria macrospora*

**DOI:** 10.1101/2025.10.28.685088

**Authors:** Lucas S. Hollstein, Kerstin Schmitt, Lucas Well, André Fleißner, Oliver Valerius, Stefanie Pöggeler

## Abstract

Hyphal fusion and sexual development in filamentous fungi rely on coordinated signaling of numerous conserved nodes such as the striatin interacting phosphatase and kinase (STRIPAK) complex or the pheromone response (PR) MAP kinase cascade (MIK2, MEK2, MAK2, HAM5). Here we used the homothallic ascomycete *Sordaria macrospora* (Sm) to screen for putative protein interactors of the SmSTRIPAK complex. Using the STRIPAK complex interactor 1 (SCI1) subunit of the complex as bait, we enriched and identified canonical SmSTRIPAK components and a determinant of communication (DOC) protein. The DOC proteins were previously described in the closely related and heterothallic species *Neurospora crassa*, functioning in allorecognition of germlings and hyphal fusions. We generated ΔSmdoc1, ΔSmdoc2 single deletion strains and the double deletion mutant ΔSmdoc1ΔSmdoc2 in *S. macrospora*. Deletion phenotypes were paradoxical: single knockouts (ΔSmdoc1 or ΔSmdoc2) were nearly sterile and sexual development was impaired, yet the double mutant (ΔSmdoc1ΔSmdoc2) exhibited wild-type fertility and development, demonstrating non-redundant and mutually antagonistic roles. Similarly, we demonstrated an impairment of the *N. crassa* Δ*doc-2* mutant in sexual development. Using gene tagging at the native locus, we performed TurboID-based proximity mapping with SmDOC1 and SmDOC2 as bait proteins. This proximity mapping demonstrated close ties of SmDOC1/2 to components of the PR MAP kinase pathway and revealed mutual SmDOC1 - SmDOC2 proximity. Yeast Two-Hybrid experiments with SmDOC1 confirmed the direct interaction with the MAP kinases MEK2 and MAK2. Fluorescence microscopy revealed that SmDOC1-TagRFP-T localized to ring-like structures around septal pores. Our results demonstrate that the DOC system is not restricted to heterothallic *N. crassa*, but also plays an essential role in the development of fruiting bodies in the homothallic fungus *S. macrospora*. These findings suggest the DOC1/2 proteins as a novel system that integrates STRIPAK and PR pathways, providing a possible mechanistic explanation for their non-additive deletion strain phenotypes.

## Introduction

Filamentous fungi have evolved complex and conserved regulatory circuits to orchestrate growth and differentiation processes, such as hyphal fusion, sexual reproduction and multicellular development. These circuits function as molecular information processing switches, by coordinating the appropriate cellular responses depending on a variety of environmental and developmental factors. Two examples of such regulatory networks are the striatin-interacting phosphatase and kinase (STRIPAK) complex and the pheromone response (PR) pathway. Both signaling pathways are conserved in unicellular and filamentous fungi and act as crucial elements to balance developmental programs in eukaryotes.

The PR signaling pathway consists of the three-tiered mitogen activated protein (MAP) kinase cascade MIK2 (MAPKKK), MEK2 (MAPKK), the terminal kinase MAK2 (MAPK) and the scaffolding protein HAM5 in filamentous ascomycetes such as the self-sterile (heterothallic) *Neurospora crassa* and the self-fertile (homothallic) *S. macrospora* (Dettmann et al., 2014; Jonkers et al., 2014; Leeder et al., 2013; Li et al., 2005; Pandey et al., 2004; Schmidt et al., 2020). The PR signaling complex regulates sexual development, hyphal fusion and melanin-dependent ascospore germination (Schmidt et al., 2020). Mounting evidence suggests bidirectional crosstalk between the STRIPAK complex and the PR pathway, with STRIPAK complex components acting as upstream regulators and downstream targets of MAP kinase cascades (Dettmann et al., 2013; Stein et al., 2020). In the heterothallic fungus *N*. *crassa*, the MAK-2 pathway is accompanied by the determinant of communication (doc) system, which was discovered by bulk segregant analysis followed by whole genome sequencing using a wild *N. crassa* population (Heller et al., 2016). The highly polymorphic “greenbeard genes” *doc-1*, *doc-2*, and *doc-3* were shown to regulate long-distance self/non-self recognition by defining five distinct communication groups (CGs) within *N. crassa* populations. The concept of “greenbeard genes” was advanced as a theoretical proposition in the form of a thought experiment, intended to provide an explanation for the proposition that altruism may be a self-serving strategy from the standpoint of genes (Gardner & West, 2010; Hamilton, 1964). Isolates with the same CG affiliation show higher communication frequencies, whereas isolates from different CGs show lower communication and cell fusion frequencies. The *doc* genes regulate the chemotropic interactions of isolates depending on the set of *doc* alleles harbored by the two individuals (allorecognition) and prevent somatic cell fusion of genetically dissimilar cells (Heller et al., 2016).

In contrast to the kinase driven mode of action in the PR pathway, the STRIPAK complex is mainly characterized by its phosphatase activity. The STRIPAK complex is a multisubunit phosphatase assembly that has been extensively characterized in fungi, where it regulates fruiting body formation, cell fusion, vegetative and sexual development, pathogenicity, secondary metabolism, and virulence of plant pathogens (Chen et al., 2023; Dettmann et al., 2013; Elramli et al., 2019; Huang et al., 2021; Kück & Pöggeler, 2024; Peterson et al., 2024; Zhang et al., 2018). In *S. macrospora*, the SmSTRIPAK complex has been investigated for over 20 years. It consists of the striatin homolog PRO11 (homolog of human striatin), the scaffolding subunit SmPP2AA (PP2AA), the catalytic subunit SmPP2Ac1 (PP2AC), the developmental protein PRO22 (STRIP1/2), the kinase activator SmMOB3 (phocein), the small coiled-coil protein STRIPAK complex interactor 1 (SCI1, homolog of human SIKE1) and sarcolemma membrane-associated protein (SLMAP) PRO45 (Beier et al., 2016; Bernhards & Pöggeler, 2011; Bloemendal et al., 2012; Nordzieke et al., 2015; Pöggeler & Kück, 2004; Reschka et al., 2018). Furthermore, the *S. macrospora* homologs of the human kinases MST1/2 (SmKIN3) and MST3/4 (SmKIN24) were shown to associate with the assembly (Frey et al., 2015b; Radchenko et al., 2018). The deletion of SmSTRIPAK components often results in impaired vegetative growth and an arrest of sexual development at early stages, leading to sterility (Kück & Stein, 2021). According to a cryo-EM structure of the human STRIPAK complex, the SmSTRIPAK complex stoichiometry consists of a PRO11 homotetramer, which functions as the main backbone of the multiprotein assembly (Jeong et al., 2021). It is accompanied by one copy of each of the other components: SmPP2AA, SmPP2Ac1, PRO22, SmMOB3, SCI1 and PRO45. The STRIPAK complex functions as a signaling hub by controlling dephosphorylation of numerous target proteins, thereby regulating diverse cellular processes. Correspondingly, phosphoproteomic analyses of SmSTRIPAK null mutants identified over 100 proteins with SmSTRIPAK-dependent phosphorylation sites (Märker et al., 2020; Stein et al., 2020).

To capture dynamic and transient protein-protein interactions of the SmSTRIPAK complex, that possibly evade conventional co-immunoprecipitation (co-IP) experiments, the Biotin identification (BioID) method was recently established for *S. macrospora* (Hollstein et al., 2022). The BioID approach relies on a promiscuous biotin ligase, in our case TurboID, which is fused to proteins of interest (Branon et al., 2018; Roux et al., 2012). This fusion construct is expressed in the organism and upon the availability of biotin, adjacent proteins get covalently labeled with biotin *in vivo*. This covalent labeling enables protein extraction and enrichment of biotinylated proteins under denaturing conditions (Fig 1). This supports efficient solubilization of membrane-bound, aggregated or hardly to resolve proteins and reduces artificial association of proteins with the bait during or after cell lysis. The biotinylated proteins are selectively enriched from denatured whole cell lysates by affinity purification and are subsequently identified by mass spectrometry (Roux et al., 2018; Sears et al., 2019). Proteins of the bait-ligase fusion harboring strain are quantified relative to a control strain (e.g. expressing an unfused biotin ligase).

**Fig 1.**
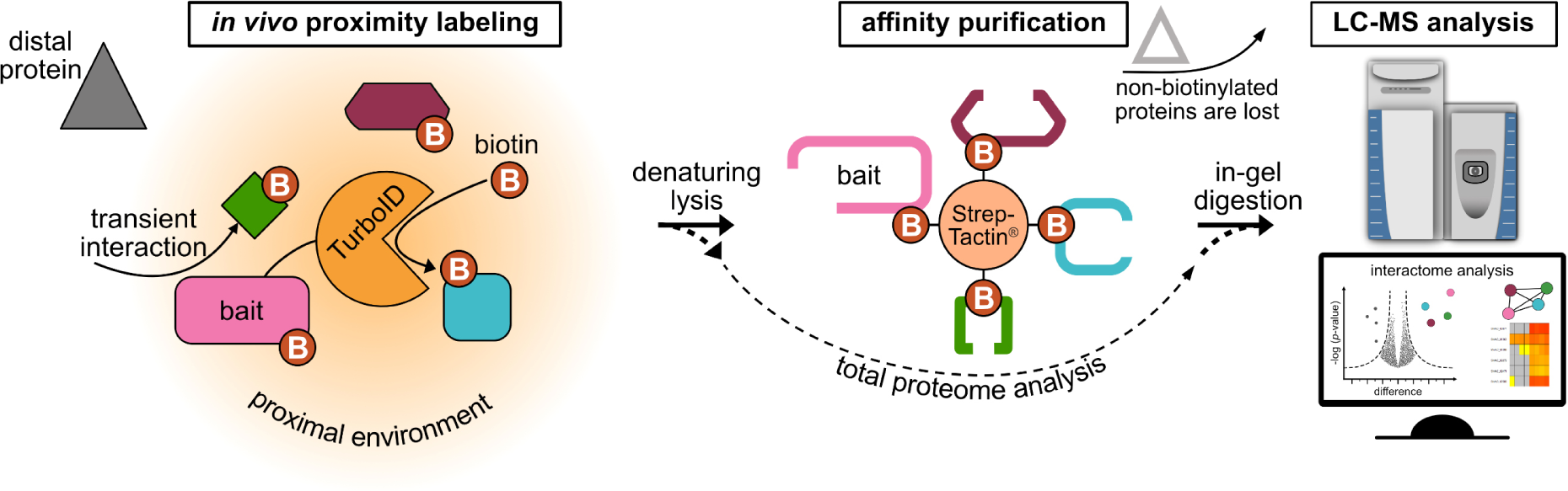
Workflow of the BioID method to capture *in vivo* co-localizing proteins. Overview of the BioID method. The promiscuous biotin ligase (here TurboID) is genetically fused to the bait protein. Proteins in the proximal environment are covalently labeled with biotin, whereas distal proteins are not labeled. Proteins are extracted under denaturing conditions and biotinylated proteins are enriched using the streptavidin mutant Strep-Tactin^®^. Non-biotinylated proteins are removed during washing, and only biotinylated proteins are eluted from the affinity purification resin. The proteins are then digested with trypsin and the resulting peptides are analyzed with LC-MS. Significantly enriched proteins are determined by relative quantification in comparison to the control (e.g. expressing an unfused biotin ligase). Input (total proteome) controls were taken prior to biotin affinity capture and analyzed by LC-MS to account for protein abundances within the different strains.

In our study, BioID mass spectrometry was applied to capture the protein networks of the SmSTRIPAK complex within the microenvironment of the subunit SCI1. Using multiple control setups, we significantly enriched and identified the *S. macrospora* homolog of *N. crassa*’s DOC-2 as a putative interactor of the SmSTRIPAK complex. We investigated and characterized the *Smdoc1* and *Smdoc2* genes of the homothallic *S. macrospora* and demonstrated that their function is not restricted to heterothallic species, but extends to homothallic *S. macrospora*. The single knockout strains ΔSmdoc1 and ΔSmdoc2 are severely impaired in sexual development, while the double knockout strain ΔSmdoc1ΔSmdoc2 display wild-type like phenotypes. For further analysis, *Smdoc1/2* fusions with the TurboID ligase and fluorescent tags were constructed and integrated at their respective *Smdoc* loci. Proximity labeling with mass spectrometry and Yeast Two-Hybrid (Y2H) interaction studies demonstrated close ties and direct interactions of the SmDOC system with the PR MAP kinase signaling module. Our findings propose a novel role of the *doc* genes in sexual reproduction and a so far uncharacterized negative regulatory network that might link the SmSTRIPAK signaling with the PR MAP kinase pathway via the SmDOC1/2 system.

## Results

### SCI1-BioID experiments identified SmDOC2 as a putative SmSTRIPAK interactor

To gain a deeper understanding of the SmSTRIPAK complex, we performed an unbiased screening for putative protein-protein interactors by applying the BioID method in combination with mass spectrometry. We used the *sci1* subunit of the SmSTRIPAK complex as bait and fused it to the *TurboID* biotin ligase for the proximity labeling (Hollstein et al., 2022). Significantly enriched proteins were determined by relative quantification with a control strain, which expresses an unfused TurboID ligase under control of the constitutive *clock-controlled gene 1* (*ccg1*) promoter from *N. crassa*. The *S. macrospora* strains were cultivated in liquid medium, cells were lysed in the presence of SDS as detergent, and biotinylated proteins were enriched from the protein crude extract using Strep-Tactin^®^ Sepharose^®^. The captured proteins were digested with trypsin, and the resulting peptides were analyzed by LC-MS as previously described in Hollstein et al. (2022). This SCI1-BioID experiment identified the already known SmSTRIPAK subunits PRO11, SmMOB3 and PRO22, thereby validating the proximity labeling approach (Fig 2AB and S6 Table). In addition to the known SmSTRIPAK components, the BioID experiment significantly enriched the Low temperature viability 1 (LTV1) protein and a protein annotated as SMAC_06902. Sequence analysis via BLASTP with SMAC_06902 as query identified the *N. crassa* greenbeard protein DOC-2 (NCU07192) with an amino acid identity of 86 % (100 % query cover) and an e-value of 0.0 as closest hit. Thus, we named the protein encoded by *SMAC_06902* SmDOC2. Using the amino acid sequence of the neighboring gene of *Smdoc2*, SMAC_06903 as query for BLASTP, we identified *N. crassa* DOC-1 (NCU07191) as closest hit with 91 % sequence identity (100 % query cover), hence we named the protein SmDOC1. Amino acid alignments of SmDOC1 and SmDOC2 with DOC-1 and DOC-2 from *N. crassa* are shown in S1AB Fig. The loci of the *doc* genes of *S. macrospora* and the common *N. crassa* lab strain FGSC 2489 are syntenic (S1C Fig).

**Fig 2.**
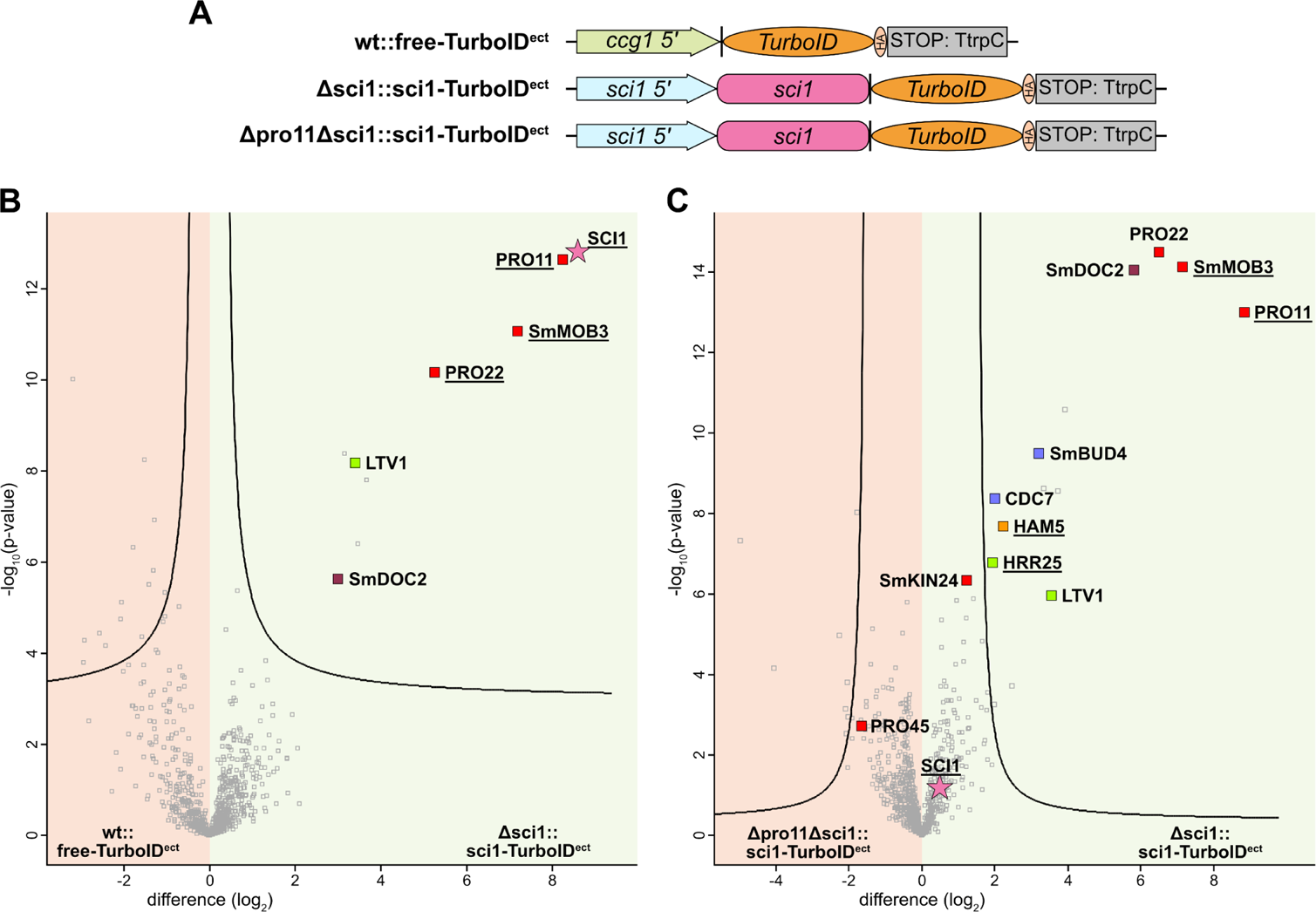
Volcano plot analysis of the SCI1-BioID experiments. **A)** Constructs for SCI1-BioID experiments using either the constitutive *clock-controlled gene 1* (*ccg1*) promoter from *N. crassa* or the native *sci1* promoter for expression. *TurboID* exhibits an N-terminal *linker* (2x GGGGS) and a C-terminal *3x HA* tag. **BC)** Volcano plot analysis of SCI1-BioID experiments using a free TurboID control (B) and a SmSTRIPAK-dependent control (C), expressing *sci1-TurboID* in the Δpro11 background. The graphs plot the difference in label free <u>q</u>uantification (LFQ) intensities (log_2_ transformed) of the control strain on the left (red) and Δsci1::sci1-TurboID^ect^ on the right (green). The -log_10_(p-value) is plotted on the y axis. Significantly enriched proteins are separated from non-significant proteins by the plotted curve. The bait protein SCI1 is marked with a pink star, other marked proteins include components of the SmSTRIPAK complex (red), components of the septation initiation network (SIN) in blue, components of the pheromone response pathway (orange), the casein kinase HRR25 and LTV1 (green) and SmDOC2 (magenta). Proteins that were identified with biotin site information are underlined. **B)** Eight biological replicates of each, wt::free-TurboID^ect^ and Δsci1::sci1-TurboID^ect^ were grown in liquid BMM medium for three days at 27 °C under constant light. Statistical parameters: FDR = 0.01; s0 = 0.1 **C)** Eight biological replicates of each, Δpro11Δsci1::sci1-TurboID^ect^ and Δsci1::sci1-TurboID^ect^ were grown in liquid BMM medium for four and three days, respectively, at 27 °C under constant light. Statistical parameters: FDR = 0.01; s0 = 2. For details such as MS/MS counts, sequence coverage and other metrics see S6 and S7 Tables.

For a more refined enrichment quantification of proteins in SCI1-BioID experiments, we constructed a novel control strain by crossing the SCI1-TurboID strain with the SmSTRIPAK mutant Δpro11. The outcome of the cross was analyzed by PCR and Southern blot experiments to properly verify the double deletion Δpro11Δsci1 (S2 Fig). This experimental setup aims to identify the striatin (PRO11)-dependent (and in a broader context the SmSTRIPAK-dependent) proxiome of SCI1-TurboID. PRO11 is the homolog of vertebrate striatin and functions as the main scaffold protein of the SmSTRIPAK complex. The Δpro11 deletion strain exhibits a sterile phenotype and only produces rudimentary ascogonia (Bloemendal et al., 2012). Consistent with the sterile phenotype of Δpro11, the newly generated Δpro11Δsci1::sci1-TurboID^ect^ strain produced no fruiting bodies, even after prolonged incubation (S3 Fig). The BioID experiment using the Δsci1::sci1-TurboID^ect^ strain in combination with the Δpro11Δsci1::sci1-TurboID^ect^ control strain showed pronounced enrichment of the SmSTRIPAK components PRO22, PRO11 and SmMOB3 (Fig 2C). The SmSTRIPAK associated kinase SmKIN24 was slightly enriched but did not pass the significance threshold. Moreover, PRO45 (SLMAP homolog), a direct interactor of SCI1, was found in the SCI1 proximity with two biotin sites independent of the Δpro11 deletion strain background. PRO45 was shown to directly interact with SCI1 in Y2H experiments (Reschka et al., 2018). Contrary to the experimental BioID setup using the free TurboID control, the bait protein SCI1-TurboID is not among the significantly enriched proteins, since it is expressed equally by both strains (Fig 2A). However, in both experimental setups SmDOC2 was significantly enriched. Upon deletion of the gene encoding the SmSTRIPAK scaffold PRO11, SmDOC2 was identified with an extremely high enrichment value of log_2_(difference) = 5.8, which corresponds to a 55-fold increase in intensity. Only the known SmSTRIPAK components PRO11, SmMOB3 and PRO22 report higher enrichment values or statistical significance than SmDOC2 in this experiment (Fig 2C). Among the other significantly enriched proteins are components of the septation initiation network (SIN), namely the proteins STE kinase CDC7 and the downstream landmark protein SmBUD4. The SIN has previously been described in connection with the STRIPAK complex (Heilig et al., 2013; Singh et al., 2011; Stein et al., 2021). The proximity labeling data also revealed significant enrichment of the scaffolding protein HAM5, which acts in the MAP kinase cascade of the pheromone response pathway (Jonkers et al., 2014). In this experiment HAM5 was identified with two peptides containing biotinylated lysine residues in seven out of the eight replicates of Δsci1::SCI-TurboID^ect^. Other notable hits among the significantly enriched proteins include the casein kinase HO and radiation repair 25 (HRR25) and the ribosome assembly factor LTV1.

Due to the vastly different developmental potential of the strains used in the SmSTRIPAK-dependent experimental setup, comparing a fertile with a sterile strain (Δsci1::sci1-TurboID^ect^ and Δpro11Δsci1::sci1-TurboID^ect^), we performed mass spectrometry analysis of the crude protein extracts prior to biotin affinity purification. This enabled us to identify false positive hits, that emerged from differences in global protein abundances, rather than changes in the proxiome of the SCI1-TurboID labeling complex. Consistent with the literature, this proteome dataset showed downregulation of proteins involved in melanin biosynthesis in the Δpro11 background, which has been observed in RNA seq experiments before (Engh et al., 2010). Most importantly, this analysis showed that the protein abundances of SmDOC1 and SmDOC2 are not affected by the deletion of *pro11* in the Δpro11Δsci1::SCI-TurboID^ect^ BioID control strain (S4 Fig, S4 and S8 Tables). Hence, the significant enrichment of SmDOC2 in the BioID experiment probing the SmSTRIPAK-dependent environment of SCI1 is based on the proximity of SCI1 and SmDOC2, rather than proteomic downregulation in the Δpro11 deletion strain background.

### Deletion of *Smdoc1* and *Smdoc2* impairs sexual development

Based on the experimental SCI1-BioID data, SmDOC2 represents the most compelling candidate for further investigation among the identified proteins for several reasons: SmDOC2 demonstrated significant enrichment against the free TurboID control. Additionally, SmDOC2 was enriched in a SmSTRIPAK-dependent manner, since its enrichment in the SCI1-BioID is drastically reduced in the Δpro11 deletion strain background. Only the SmSTRIPAK components PRO11, SmMOB3, and PRO22 showed higher enrichment values, placing SmDOC2 in an outstanding category of potential interactors. Unlike the other candidates described above, SmDOC proteins have been identified across multiple proteomic approaches using SmSTRIPAK components as bait (Nordzieke et al., 2015; Reschka, 2018). Another argument for prioritizing SmDOC2 lies in the functional overlap between DOC proteins and the SmSTRIPAK complex. Both regulatory systems control hyphal fusion events, a fundamental process in fungal development. The DOC proteins of *N. crassa* mediate pre-contact allorecognition between germlings and control this first checkpoint during somatic cell fusion (Heller et al., 2016). Similarly, the SmSTRIPAK complex signaling is essential for hyphal fusion and multicellular development in filamentous fungi (Kück & Pöggeler, 2024). This functional convergence might place the SmDOC proteins as a novel SmSTRIPAK-interacting regulatory node connecting these two critical signaling pathways. Therefore, we decided to functionally characterize *Smdoc1* and *Smdoc2* in *S. macrospora*.

For functional characterization of *Smdoc1/2* in *S. macrospora*, single deletion strains of ΔSmdoc1, ΔSmdoc2 and the double deletion strain ΔSmdoc1ΔSmdoc2 were generated (Fig 3A) and verified by PCR and Southern hybridization experiments (S5-S7 Figs). When grown on agar plates under standard conditions, the ΔSmdoc1 and ΔSmdoc2 single knockouts exhibit severe impairments in fruiting body formation and only a few fruiting bodies are produced (Fig 3B). Ascospores harvested from these fruiting bodies were viable. Additionally, the single knockouts ΔSmdoc1 and ΔSmdoc2 exhibit an unusually dense mycelium layer on top of the agar, which was cut-resistant when cutting agar pieces with a lancet. Vegetative growth analyses were performed in 30 cm long race tubes filled with synthetic SWG fructification medium. In this experiment, we noticed a delay in colony establishment of ΔSmdoc1 and ΔSmdoc2. The single knockouts took up to two days post-inoculation until the growth front of the mycelium was visible to the eye. Accordingly, this slower colony establishment of ΔSmdoc1 and ΔSmdoc2 resulted in shorter total growth when measurements were taken after three days. However, once the colony establishment stage was passed, the single knockouts reached wild-type like growth rates per day (∼ 25-30 mm per 24 h). The double knockout ΔSmdoc1ΔSmdoc2 did not show any delay in colony establishment or vegetative growth rate and was indistinguishable from the wild type in this experiment (S8 Fig). Surprisingly the ΔSmdoc1ΔSmdoc2 double knockout produced fruiting bodies in quantities similar to the wild type (Fig 3B). The double knockout does not display any abnormal growth phenotypes. The single deletion strains ΔSmdoc1 and ΔSmdoc2 were complemented by ectopic integration of constructs harboring the *Smdoc1* or *Smdoc2* ORF flanked by 1 kb upstream and downstream regions. The fruiting body formation of ΔSmdoc2 reverted to wild-type levels upon reintroduction of the *Smdoc2* gene. Similarly, the ΔSmdoc1::Smdoc1^ect^ strain regained the ability to produce fruiting bodies, however the density of fruiting bodies was increased when compared to the wild type (S9 Fig).

**Fig 3.**
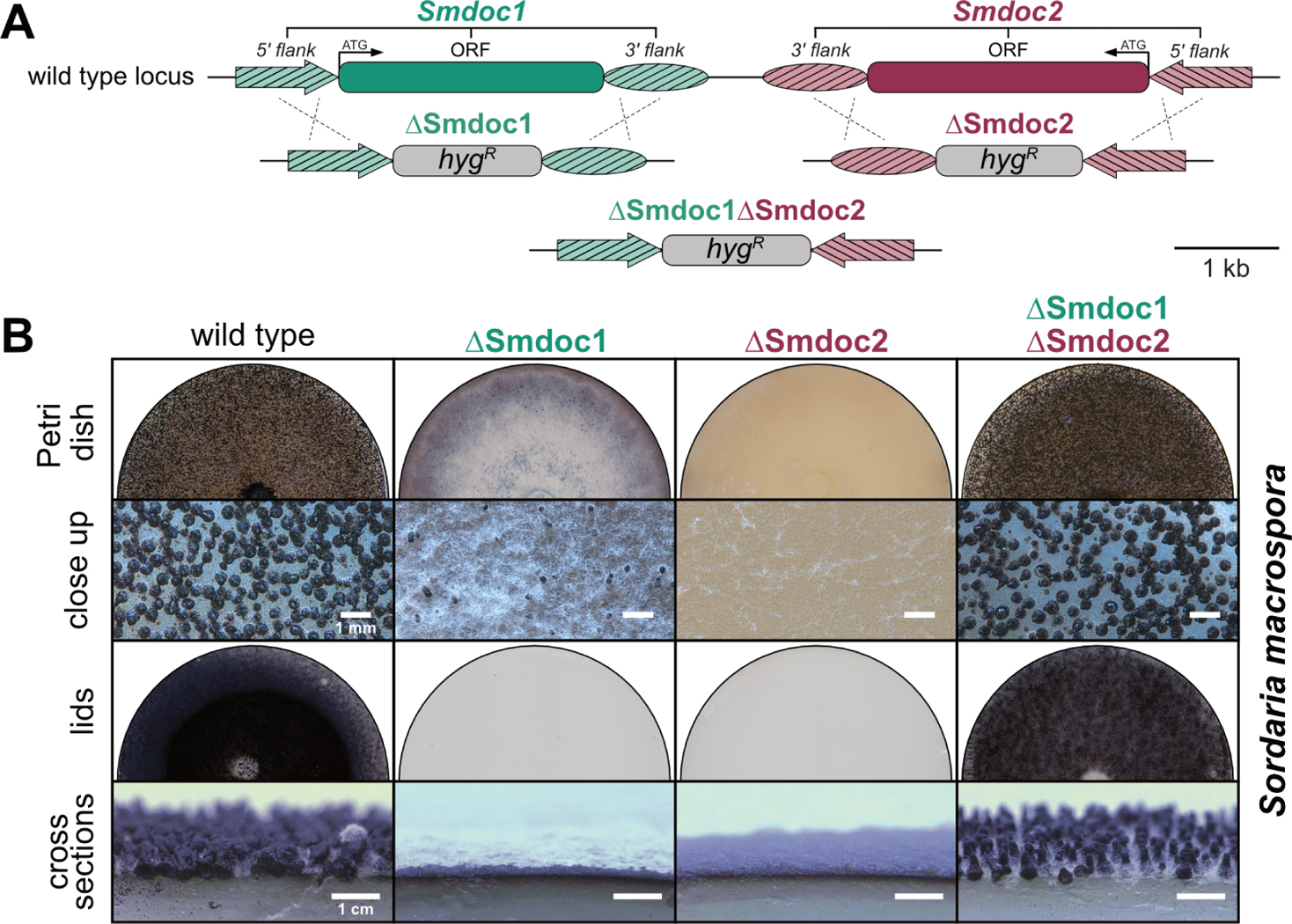
The *Smdoc* genes control sexual development in *S. macrospora*. **A)** Genetic map of the *Smdoc* locus in the homothallic ascomycete *S. macrospora*. The single knockouts ΔSmdoc1 and ΔSmdoc2 as well as the double knockout ΔSmdoc1ΔSmdoc2 were generated via homologous recombination. The knockout cassette (*hyg^R^*) confers resistance to the hygromycin antibiotic. 1 kb flanking regions were used to target the cassette to the respective locus. **B)** The single spore isolates were grown at 27 °C on solid *Sordaria* Westergaard’s (SWG) fructification medium. Pictures of the Petri dishes, the close ups, the lids and the cross sections were taken after 14 days of incubation. Once the black ascospores are fully matured inside the fruiting bodies, they are forcefully ejected towards the light source and stick to the lids of the Petri dishes, thereby staining them black. *hyg^R^*, hygromycin resistance cassette expressing the *hygromycin B phosphotransferase* gene from *E. coli* under control of the constitutive *trpC* promoter from *A. nidulans*

Due to the severe impairment in sexual development in the homothallic *S. macrospora*, we questioned whether sexual development of the heterothallic *N. crassa doc* knockout strains was impaired. Previous studies in *N. crassa* focused on aspects of pre-contact communication in germlings, but did not investigate the sexual development. The *N. crassa* Δ*doc-1* and Δ*doc-2* deletion strains were described to be “macroscopically indistinguishable” from the commonly used lab strain FGSC 2489 during vegetative growth (Heller et al., 2016). The Δ*doc-1* and Δ*doc-1*Δ*doc-2* deletion strains produced the female sexual structures, the protoperithecia, in a similar time frame as the *N. crassa* wild type (Fig 4). However, the Δ*doc-2* deletion strain produced protoperithecia only after prolonged incubation. Fertilization of Δ*doc-2* protoperithecia with conidia of the wild type, Δ*doc-1*, Δ*doc-2*, or Δ*doc-1*Δ*doc-2* as male crossing partners did not result in the formation of perithecia. Both mating types were tested, but Δ*doc-2* protoperithecia never progressed in sexual development after fertilization. Interestingly, the fertilization of wild-type protoperithecia with Δ*doc-2* microconidia resulted in normal sexual development with formation of perithecia and production of ascospore numbers similar to the wild type. Crossings of Δ*doc-1* x Δ*doc-1* and Δ*doc-1*Δ*doc-2* x Δ*doc-1*Δ*doc-2* showed wild-type like development of sexual structures and ascospore production.

**Fig 4.**
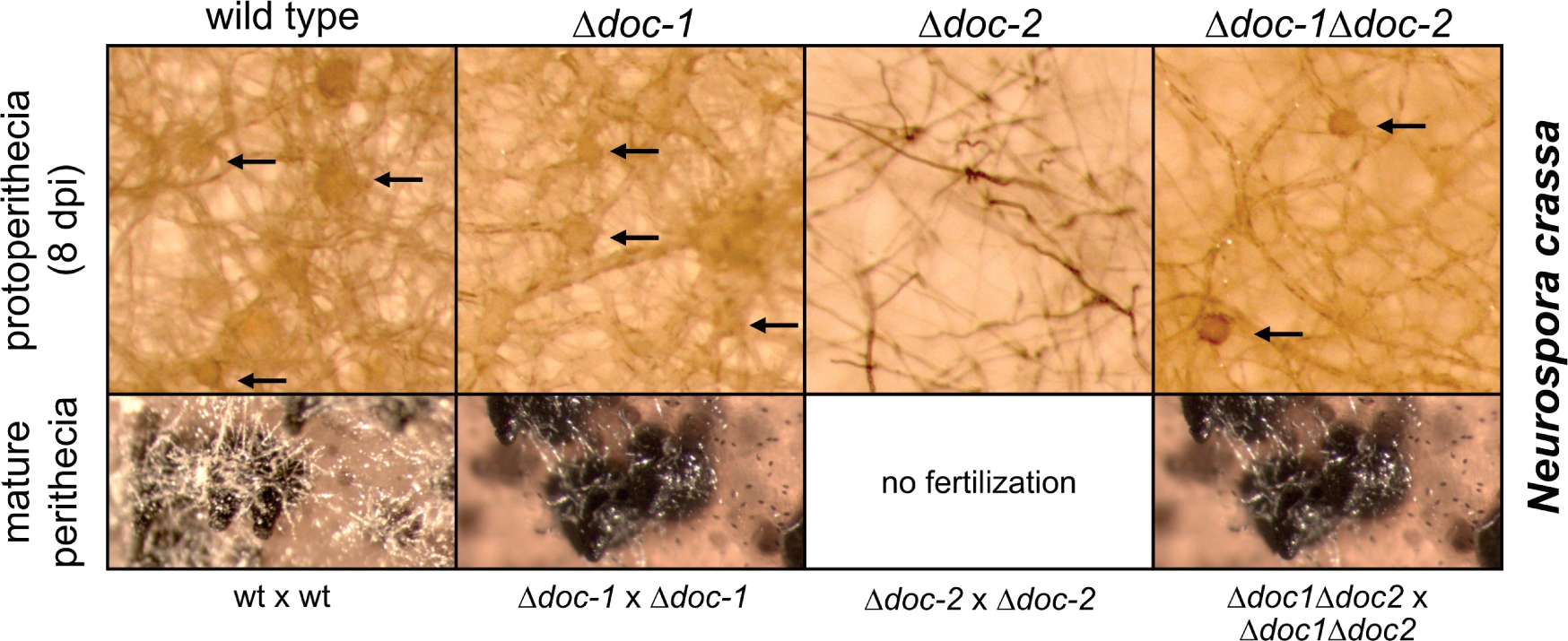
Sexual development of Δ*doc-2* in *N. crassa* is impaired. Sexual crosses of the wild type, Δ*doc-1*, Δ*doc-2* and Δ*doc-1*Δ*doc-2* of the heterothallic ascomycete *N. crassa*. Formation of the female sexual structures, the protoperithecia, is delayed in Δ*doc-2*. Fertilization of the protoperithecia with the male gametes, the microconidia resulted in mature perithecia containing ascospores, except for Δ*doc-2*. Here, fertilization of the Δ*doc-2* protoperithecia with microconidia from any of the strains did not result in perithecia formation. dpi, days post inoculation

### The SmDOC1/2-proxiomes include components of the MAK2 MAP kinase cascade

To systematically investigate the putative interaction networks of SmDOC1 and SmDOC2 *in vivo*, we constructed strains expressing the *Smdoc1-TurboID* or *Smdoc2-TurboID* fusions from their native loci, under control of their native 5’ regions (Figs 5A and 6A). This approach aims to preserve endogenous regulatory networks and maintain physiological expression levels, avoiding artifacts that could arise from ectopic integration or overexpression systems.

**Fig 5.**
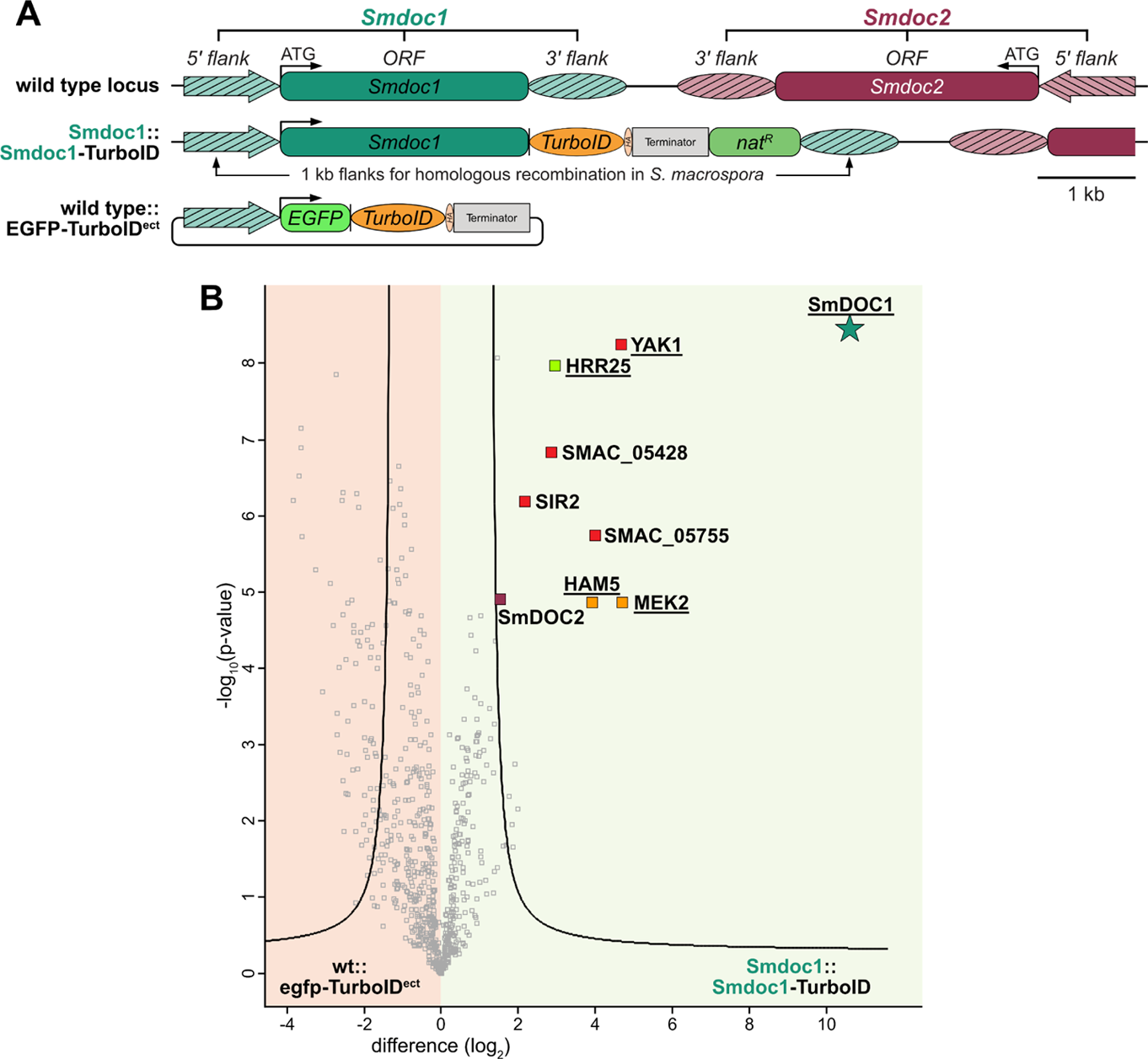
Volcano plot analysis of the SmDOC1-BioID experiment. **A)** Genomic organization of the *Smdoc1* locus in *S. macrospora.* The Smdoc1::Smdoc1-TurboID strain was constructed by integration of the *Smdoc1-TurboID* fusion at the native *Smdoc1* locus. The fusion gene is accompanied by a nourseothricin resistance cassette (*nat^R^*). Both genes are flanked by 1 kb 5’ and 3’ *Smdoc1* regions for homologous recombination. For relative quantification, we ectopically integrated an *egfp-TurboID* fusion gene under control of the 1 kb 5’ *Smdoc1* region into the *S. macrospora* wild type (wt::egfp-TurboID^ect^). Expression of the *TurboID* ligase is terminated by the terminator of the *anthranilate synthase trpC* gene of *A. nidulans*. TurboID is C-terminally tagged with a triple HA-Tag. **B)** Statistical volcano plot analysis of the BioID experiment using SmDOC1-TurboID as bait. The graph plots the difference in label free quantification (LFQ) intensities (log_2_ transformed) of the control strain on the left and the *Smdoc1-TurboID* fusion on the right. The -log_10_(p-value) is plotted on the y axis. Significantly enriched proteins are separated from non-significant proteins by the plotted curve. Proteins that were identified with biotin site information are underlined. The bait protein SmDOC1 is marked with a star colored in dark green. Among the significantly enriched proteins, components of the MAK2 pathway are marked in orange and the casein kinase HRR25 is marked in green. 5 biological replicates of wt::egfp-TurboID^ect^ and Smdoc1::Smdoc1-TurboID were grown in liquid BMM medium for three days at 27 °C under constant light. Statistical parameters of the volcano plots: FDR = 0.01; s0 = 2. For details such as MS/MS counts, sequence coverage and other metrics see S9 Table. *nat^R^,* nourseothricin resistance cassette expressing the *nourseothricin acetyltransferase* gene from *S. noursei* under control of the constitutive *trpC* promoter from *A. nidulans*.

**Fig 6.**
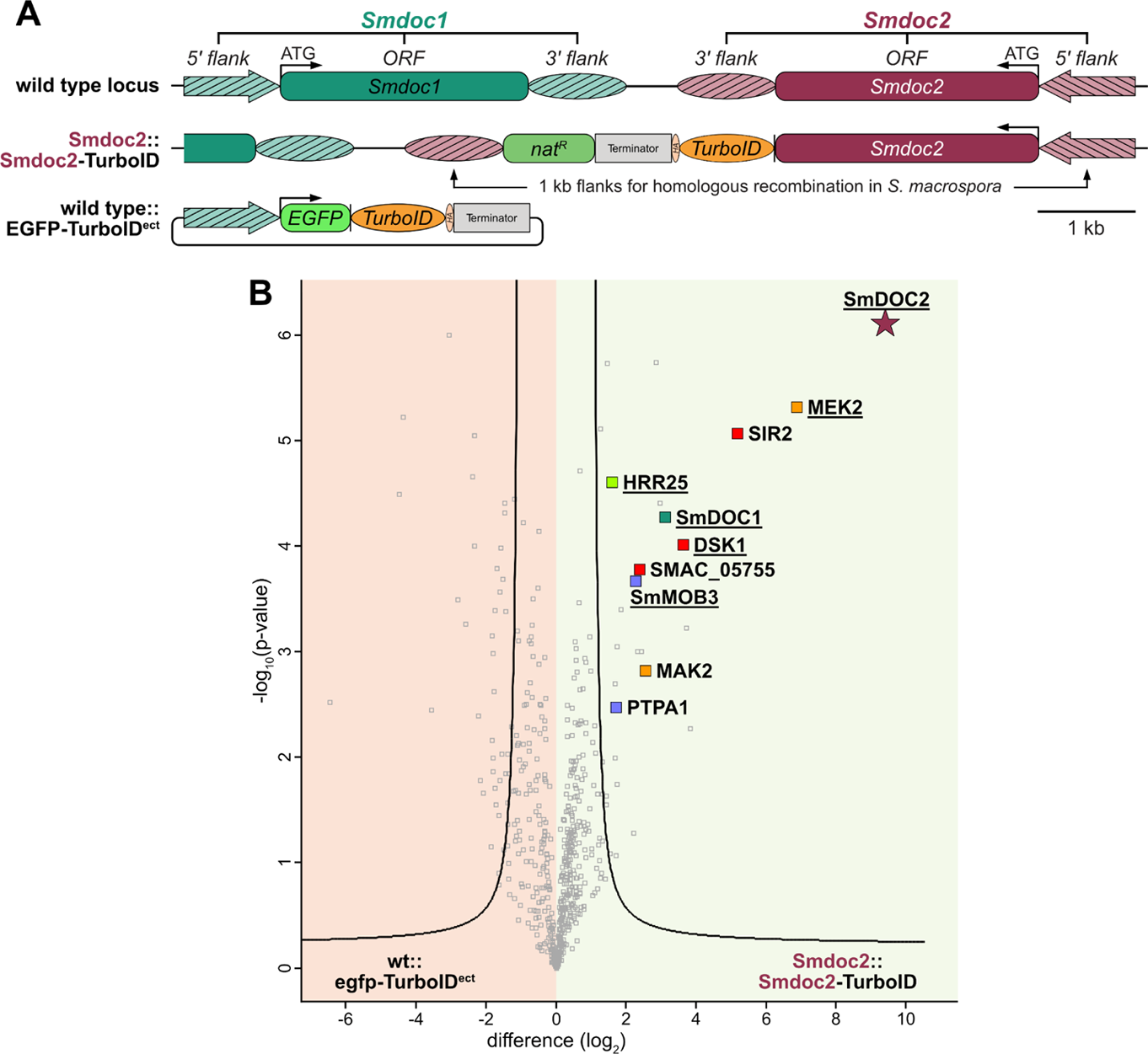
Volcano plot analysis of the SmDOC2-BioID experiment. **A)** Genomic organization of the *Smdoc2* locus in *S. macrospora.* The Smdoc2::Smdoc2-TurboID strain was constructed by integration of the *Smdoc2-TurboID* fusion at the native *Smdoc2* locus. The fusion gene is accompanied by a nourseothricin resistance cassette (*nat^R^*). Both genes are flanked by 1 kb 5’ and 3’ *Smdoc2* regions for homologous recombination. For relative quantification in BioID experiments, we ectopically integrated an *egfp-TurboID* fusion gene under control of the 1 kb 5’ *Smdoc1* region into the *S. macrospora* wild type (wt::egfp-TurboID^ect^). Expression of the *TurboID* ligase is terminated by the terminator of the *anthranilate synthase trpC* gene of *A. nidulans*. TurboID is C-terminally tagged with a triple HA-Tag. **B)** Statistical volcano plot analysis of the BioID experiments using SmDOC2-TurboID as bait. The graph plots the difference in label free quantification (LFQ) intensities (log_2_ transformed) of the control strain on the left and the *Smdoc2-TurboID* fusion on the right. The -log_10_(p-value) is plotted on the y axis. Significantly enriched proteins are separated from non-significant proteins by the plotted curve. Proteins that were identified with biotin site information are underlined. The bait protein SmDOC2 is marked with a star colored magenta. Among the significantly enriched proteins, components of the MAK2 pathway are marked in orange, the casein kinase HRR25 is marked in green and the SmSTRIPAK associated protein SmMOB3 and PTPA1 are marked in blue. 4 biological replicates of wt::egfp-TurboID^ect^ and Smdoc2::Smdoc2-TurboID were grown in liquid BMM medium for four days at 27 °C under constant light. Statistical parameters of the volcano plots: FDR = 0.01; s0 = 2. For details such as MS/MS counts, sequence coverage and other metrics see S10 Table. *nat^R^,* nourseothricin resistance cassette expressing the *nourseothricin acetyltransferase* gene from *S. noursei* under control of the constitutive *trpC* promoter from *A. nidulans*.

Both C-terminally tagged *Smdoc*-fusions were integrated via homologous recombination into the Δku80 strain using 1 kb flanking regions (Groth et al., 2021). The *Smdoc* baits were fused to *TurboID* via a GGGGSGGGS linker to allow flexibility. Transcription is terminated by the terminator of the *anthranilate synthase* gene of *A. nidulans* (Figs 5A and 6A). The thereby generated Smdoc1::Smdoc1-TurboID and Smdoc2::Smdoc2-TurboID strains were analyzed by PCR and Southern hybridization experiments to verify the correct integration of the bait-TurboID fusion construct and to verify the absence of the wild-type *Smdoc1* or *Smdoc2* gene (S10 and S11 Figs). Importantly, both strains retained fertility and did not show any phenotypes of the knockout strains, demonstrating functionality of the *Smdoc-TurboID* fusions. While the SmDOC1-TurboID (130 kDa) and SmDOC2-TurboID (133 kDa) fusion proteins exhibit large molecular weights, the unfused TurboID ligase in the control strain wt::free-TurboID^ect^ (39 kDa) appears small by comparison. However, large proteins are often exposed to translational challenges, including the higher risk of translational errors or co-translational misfolding due to the prolonged synthesis duration (Fernandes et al., 2017; Lyu et al., 2021). To balance the size discrepancy between the SmDOC-fusion proteins and the free TurboID control, we constructed an appropriate BioID control by genetically fusing *egfp* to *TurboID* via a GGGGSGGGGS linker. We chose *egfp*, because it is known to fold autonomously and has been extensively validated for its use in *S. macrospora* in complementation assays of gene knockouts, fluorescence microscopy and GFP-Trap experiments (Kelkar et al., 2012; Nordzieke et al., 2015; Pöggeler et al., 2003; Werner et al., 2019). The resulting *egfp-TurboID* fusion gene encoded a protein of 66 kDa and was put under the control of the 1 kb 5’ region of *Smdoc1* rather than an overexpression or constitutive promoter to more closely match the expression and biotinylation activity of the SmDOC-TurboID fusion proteins. The *egfp-TurboID* construct was ectopically integrated into the *S. macrospora* wild type and yielded fertile single spore isolates. The biotinylation activity of the SmDOC1-TurboID, SmDOC2-TurboID and EGFP-TurboID fusion proteins was assessed in Western blot experiments using a Streptavidin-HRP conjugate for signal detection (S12 and S13 Figs). Fluorescence microscopy of this wt::egfp-TurboID^ect^ strain showed uniform EGFP signal in the cytoplasm without hotspots, which could indicate protein aggregation or degradation (S14 Fig). The wt::egfp-TurboID^ect^ control strain was used for relative quantification in the SmDOC1-BioID and SmDOC2-BioID experiments.

The SmDOC1 BioID experimental setup consisted of five biological replicates of the Smdoc1::Smdoc1-TurboID strain and the wt::egfp-TurboID^ect^ control strain for relative quantification. In total 1563 proteins were identified in the Strep-Tactin^®^ Sepharose^®^ eluate samples. Filtering for detection of any given protein in at least four out of five replicates in either of the strains reduced the protein count to 720.

The bait protein itself, SmDOC1, was the highest enriched protein (Fig 5B) and a total of 14 phosphorylated residues and 13 biotinylated sites were identified for SmDOC1. Other significantly enriched proteins included (sorted from high to low enrichment values): the protein kinase YAK1 (SMAC_02146), the MAP kinase kinase MEK2 (SMAC_06526) and the MAK2 scaffold HAM5 (SMAC_02471), protein SMAC_05755 containing a domain of unknown function 7624 (DUF7624), the casein kinase HRR25 (SMAC_01363), the putative septal pore associated protein SMAC_05428, the deacetylase SIR2 (SMAC_12019) and SmDOC2 (SMAC_06902). Biotin site information was recovered for the bait SmDOC1 itself, YAK1, MEK2, HAM5 and HRR25. In this SmDOC1 BioID experiment, no SmSTRIPAK components were significantly enriched. For details see S9 Table.

For the SmDOC2-BioID experiment, four biological replicates of Smdoc2::Smdoc2-TurboID and the control strain wt::egfp-TurboID^ect^ were used (Fig 6A). The statistical analysis of this setup reported the bait, SmDOC2 itself as the most highly enriched protein, including the recovery of nine distinct phosphorylated residues and 12 biotinylated sites (Fig 6B). In total 1279 proteins were identified in the Strep-Tactin^®^ Sepharose^®^ eluate samples. Filtering for complete detection in all four replicates in either strain, reduced the protein count to 573. Notably, the candidates enriched in the SmDOC1-BioID and SmDOC2-BioID experiments showed considerable overlap. Consistent with the proteinaceous environment of SmDOC1, the SmDOC2 experiment significantly enriched components of the pheromone response pathway, namely MEK2 (SMAC_06526) and MAK2 (SMAC_03492). Additionally, the deacetylase SIR2 (SMAC_12019), SMAC_05755, and the casein kinase HRR25 (SMAC_01363) were identified with both SmDOC proteins as bait. In accordance with the enrichment of SmDOC2 in SmDOC1-BioID experiments, the SmDOC2-BioID enriched SmDOC1 reciprocally. Among the other significantly enriched proteins of the SmDOC2 BioID were (sorted from high to low enrichment values): the protein kinase DSK1 (SMAC_01589), the SmSTRIPAK subunit SmMOB3 (SMAC_00877) and the PP2A phosphatase activator PTPA1 (SMAC_03446). Biotin site information was recovered for peptides of SmDOC2, MEK2, SmDOC1, DSK1 and HRR25. For details see S10 Table.

Given the consistent enrichment of components of the PR signaling cascade (HAM5 and MEK2 in SmDOC1-BioID; MEK2 and MAK2 in SmDOC2-BioID), we constructed a pathway-specific BioID control strain to dissect the MAK2-dependent interactions of SmDOC2. The *S. macrospora* Δmak2 deletion is sterile, exhibits a hyphal fusion defect and is impaired in ascospore germination (Schmidt et al., 2020). The Δmak2 deletion strain was generated by replacement of the *mak2* ORF with a resistance cassette (*hyg^R^*), which was subsequently removed using the FLP/*FRT* recombination to create a markerless Δmak2 deletion strain (Kopke et al., 2010; Schmidt et al., 2020). The new BioID strain was constructed by crossing the Δmak2 deletion strain in the spore color mutant background fus1-1 with the Smdoc2::Smdoc2-TurboID (*nat^R^*) strain. Spores were picked from recombinant perithecia, and sterile isolates (*nat^R^*) were tested for the deletion of *mak2* by PCR (S15 Fig). The resulting Δmak2; Smdoc2::Smdoc2-TurboID strain was used to explore the MAK2-dependent proxiome of SmDOC2-TurboID. The biotinylation activity of the SmDOC2-TurboID fusion protein was not affected by the introduction of the Δmak2 deletion strain background (S16 Fig). Consistent with the *mak2* deletion, MAK2 peptides were absent from the BioID eluates as well as the input controls of Δmak2, Smdoc2::Smdoc2-TurboID. The significant enrichment of the proteins SIR2, SmDOC1, DSK1, MEK2, SmMOB3, PTPA1 and HRR25 was not affected in the Δmak2 deletion background when compared to the wt::egfp-TurboID^ect^ strain (S17 Fig). Biotin sites were detected for SmDOC2, MEK2, SmDOC1, DSK1 and HRR25 in the replicates of Δmak2; Smdoc2::Smdoc2-TurboID. The LFQ intensities for SIR2, SmDOC1 and DSK1 are slightly increased in the Δmak2 background when compared to the Smdoc2::Smdoc2-TurboID strain, whereas the intensities for MEK2 are reduced upon deletion of *mak2* (S11 Table). An additional candidate among the significantly enriched protein is SmBRO1 (SMAC_01835), a homolog of the *N. crassa* BRO1 (NCU08001). BRO1 was reported to be essential in *N. crassa* and investigation of *bro1* knockdowns revealed a defect in germling fusion (Hammadeh, 2022). However, it needs to be noted that the enrichment value of SmBRO1 in Δmak2; Smdoc2::Smdoc2-TurboID is rather low (log2 difference = 2.7), and the input controls show a slight upregulation of SmBRO1 in the Δmak2 background. Interestingly, biotin sites of SmBRO1 were identified in all four biological replicates in the Δmak2 background, but not in the Smdoc2::Smdoc2-TurboID strain or the free EGFP-TurboID control. Biotinylated BRO1 peptides are also absent from the input control samples.

### SmDOC1/2 BioID experiments enrich two MEK2 isoforms

Previous proteogenomic analysis of *S. macrospora* identified novel alternative splicing events, contributing to a refinement of the genome annotation (Blank-Landeshammer et al., 2019). Among them, a novel intron retention event was identified that affects the PR signaling component MEK2, resulting in an extended MEK2 isoform with an alternative and extended protein C-terminus (MEK2-t2). Proteomic analysis showed a downregulation of the MEK2-t2 isoform during sexual development starting at three days of incubation (Blank-Landeshammer et al., 2019). The SmDOC1- and SmDOC2-BioID experiments enriched unique peptides of both, the shorter MEK2-t1 and the novel MEK2-t2 isoform in the BioID eluates. Interestingly, no MEK2-t2 peptides were identified in the BioID eluates of control samples or the input control samples (S12 and S13 Table).

### SmDOC1 directly interacts with components of the PR signaling pathway

To verify the putative interactions of the SmDOC1/2 proteins with components of the PR signaling pathway proposed by the *in vivo* proximity labeling data, we performed Y2H analysis. For this purpose, *Smdoc1*, *Smdoc2*, *mek2-t1*, *mek2-t2*, *ham5* and *mak2* cDNAs were cloned into vectors pGBKT7 (bait) and pGADT7 (prey). These vectors harbor the Gal4 DNA-binding domain (BD) or the Gal4 activation domain (AD), respectively. When SmDOC1 was fused to the AD, it was able to interact with BD-MEK2-t1, BD-MEK2-t2 and BD-MAK2 (Fig 7). An interaction of AD-SmDOC1 with BD-HAM5 was not detected. The negative controls showed no evidence of autoactivation (S18AB Fig). Conversely there was no interaction of SmDOC1 as bait with any of the tested AD-prey fusions, although the positive control with AD-ranBPM x BD-SmDOC1 (Tucker et al., 2009) yielded positive results (S18C Fig). Matings of the positive control AD-ranBPM with BD-SmDOC2 resulted in reduced cell growth when compared to the mating of AD-ranBPM with BD-SmDOC1 (S18D Fig).

**Fig 7.**
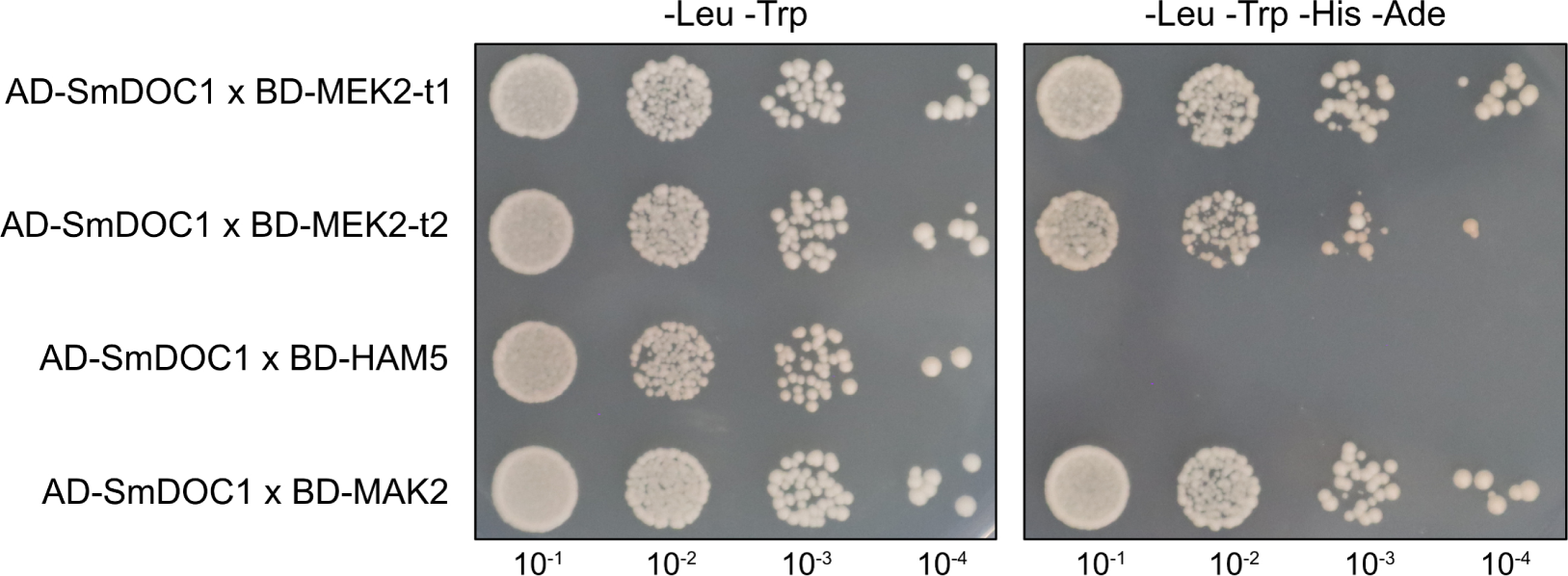
Yeast Two-Hybrid assay for the SmDOC1 interaction with PR components. Constructs of *Smdoc1*, *mek2-t1*, *mek2-t2*, *ham5* and *mak2* were either fused to the Gal4 activation domain (AD) or the Gal4 DNA-binding domain (BD). Serial dilutions of the yeast cells (10^-1^ = 0.5 x 10^5^ cells) were spotted onto synthetic defined (SD) medium lacking leucine and tryptophan (-Leu-Trp) or lacking leucine, tryptophan, histidine and adenine (-Leu-Trp-His-Ade). Cells harboring both plasmids (AD and BD) grow on SD-Leu-Trp. Positive interactions are indicated by hybridized yeast cell survival on selective SD-Leu-Trp-His-Ade medium. The plates were incubated at 30 °C. AD, activation domain; BD, binding domain

### SmDOC1 localizes to a ring-like structure around the septal pore

Fluorescence microscopy was performed to determine the subcellular localization of SmDOC1 and SmDOC2. Therefore, we performed *in locus* tagging of *Smdoc1* with the red fluorescent protein *TagRFP-T* at its C terminus (Shaner et al., 2008). The *Smdoc1-TagRFP-T* construct was integrated at the native *Smdoc1* locus using homologous recombination in the Δku80 strain based on homologous 1 kb 5’ and 3’ flanking regions. Nourseothricin-resistant single spore isolates were analyzed by PCR and Southern blot experiments were used to verify the absence of untagged wild-type *Smdoc1* (S10 Fig). The resulting Smdoc1::Smdoc1-TagRFP-T strain showed wild-type like sexual development and production of fertile ascospores. Microscopic analyses showed weak, but distinct and consistent fluorescence signal at the septa of mature hyphae (Fig 8A). The magnification of a single septum shows a ring-like fluorescence signal at the septal pore (Fig 8B). This signal was never observed in the *S. macrospora* wild type (S19 Fig). Ectopic integration of the *Smdoc1-TagRFP-T* fusion under control of the constitutive *ccg1* promoter from *N. crassa* rather than the native *Smdoc1* 5’ region did not result in increased fluorescence signal intensity (S19 Fig).

**Fig 8.**
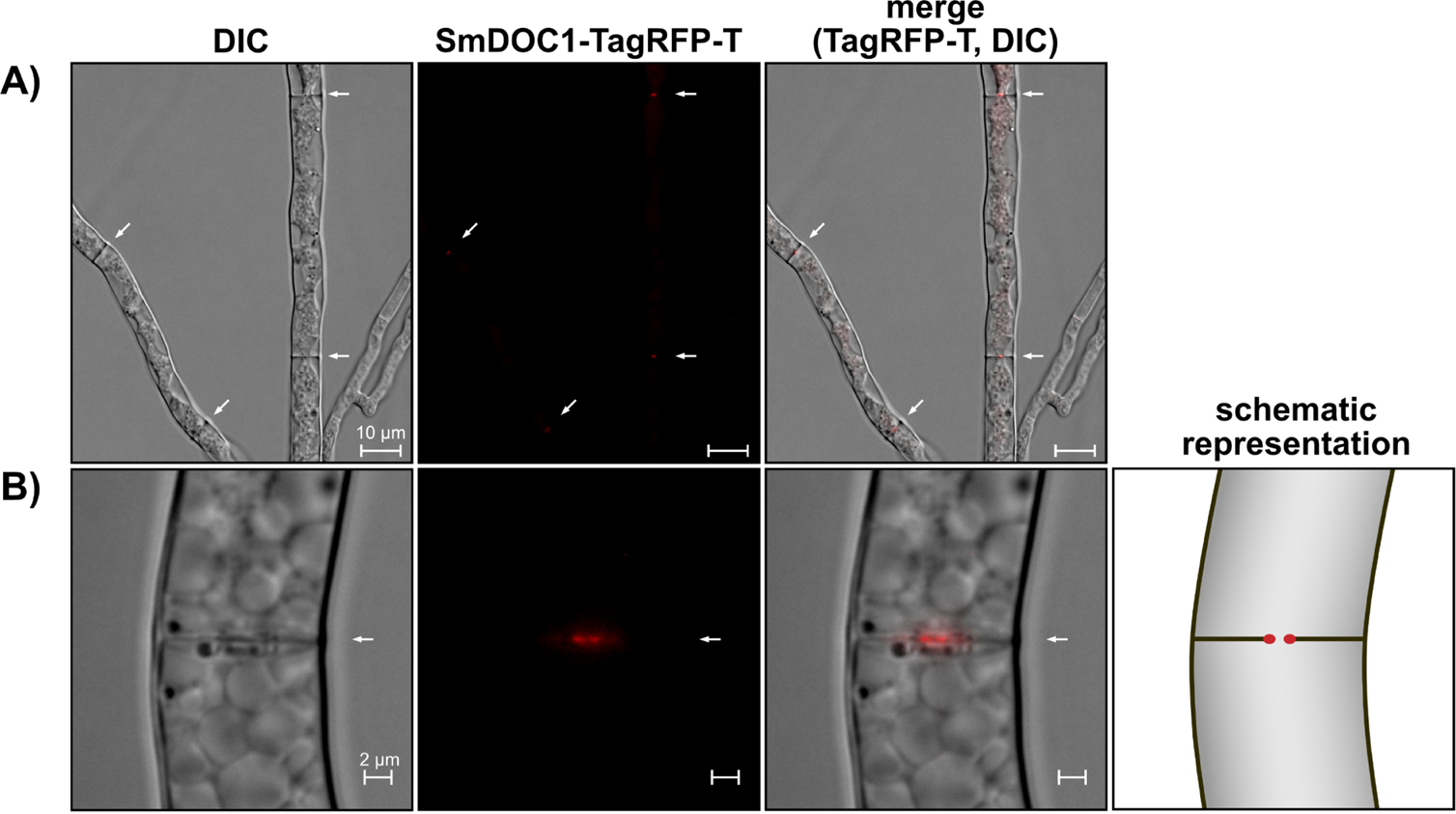
SmDOC1-TagRFP-T localizes to a ring-like structure at the septal pore. *S. macrospora* expressing the *Smdoc1-TagRFP-T* fusion from the native *Smdoc1* locus using the native 5’ region. Arrows indicate septa within the focal plane. SmDOC1-TagRFP-T localizes to septa of mature hyphae in a ring-like structure around the septal pore. The samples were grown on microscopic slides covered in BMM. DIC, differential interference contrast, scale bars are indicated.

To determine the subcellular localization of SmDOC2, we generated strains expressing *egfp* and *TagRFP-T* tagged variants of *Smdoc2* by *in locus* integration and subsequent verification by PCR and Southern blot experiments (S20 Fig). The resulting strains were fertile and did not show the ΔSmdoc2 phenotype, demonstrating the functionality of the tagged SmDOC2 fusion proteins. In both strains, only weak signals were observed in the membrane or cell wall region of the hyphae (S21 Fig). However, similar signals were detected in the wild type and therefore might be attributed to autofluorescence. Expression of *Smdoc2-egfp* or *Smdoc2-TagRFP-T* under control of the constitutive *ccg1* promoter did not result in improved signal yield (S21 Fig).

## Discussion

### The proxiome of SCI1 embraces the role of the SmSTRIPAK complex as a signaling hub

This study mapped the *in vivo* protein neighborhood of the SCI1 subunit of the SmSTRIPAK complex in the filamentous ascomycete *S. macrospora* using proximity labeling with biotin. For this, two independent control setups were applied to assure reliable relative quantification of the mass spectrometry data and avoid false positive hits. The proxiome dataset demonstrates the capture of already known as well as previously suspected links between the SmSTRIPAK complex and other conserved fungal signaling pathways such as the SIN, MAPK signaling and ribosome biogenesis.

One example of the interaction of the SmSTRIPAK with other conserved signaling pathways is the significant enrichment of the SIN component CDC7 and the downstream protein SmBUD4. This aligns with the well documented connection between the STRIPAK complex and the SIN, first established in *Schizosaccharomyces pombe*, where the STRIPAK homolog (termed SIN-inhibitory PP2A (SIP) complex) negatively regulates the SIN by dephosphorylating the scaffolding protein Cdc11p (Singh et al., 2011). Additionally, our findings align with phosphoproteomic evidence from SmSTRIPAK mutant strains, which revealed SmSTRIPAK-dependent phosphorylation of the *S. macrospora* SIN components CDC7 (two sites) and SmBUD4 (five sites). The model suggests that deletion of SmSTRIPAK subunits results in higher phosphorylation of CDC7, thereby inhibiting SIN assembly (Stein et al., 2021). However, the interactions of the SmSTRIPAK with the SIN are not restricted to CDC7 and SmBUD4, but also include the SIN kinase SmKIN3, which exhibits SmSTRIPAK dependent phosphorylation and was shown to physically interact with PRO11 in co-IP experiments (Frey et al., 2015b). More evidence for the capture of biologically relevant protein-protein interactions in the SCI1 proxiome dataset is posed by the co-enrichment of the functionally coupled pair HRR25-LTV1. This evolutionary conserved circuit, present in both yeast and humans, is involved in the biogenesis of the small 40S ribosomal subunit through HRR25-mediated phosphorylation of the assembly factor LTV1. The phosphorylation event triggers the release of LTV1 from pre-40S ribosomal subunits and allows subunit maturation (Ghalei et al., 2015; Schäfer et al., 2006). Deletion of LTV1 in yeast results in increased sensitivity to cold temperatures (Loar et al., 2004; Seiser et al., 2006). Notably, HRR25 was previously detected with moderate spectral counts in SCI1-eGFP pulldown experiments (Reschka, 2018). Phosphoproteomic analyses of the SmSTRIPAK mutant strains ΔSmpp2ac1, Δpro11, Δpro22 and the double mutant Δpro11Δpro22 reported increased phosphorylation of S134 of LTV1 (Märker et al., 2020; Stein et al., 2020). Recent transcriptome profiling of STRIPAK mutants in the basidiomycete *Cryptococcus neoformans* suggests a connection of ribosome biogenesis with the STRIPAK. Deletion of the catalytic PP2A subunit in the Δpph22 mutant resulted in strong downregulation of gene clusters associated with ribosome assembly and RNA processing (Peterson et al., 2025).

Furthermore, the enrichment of HAM5 in the SCI1 proxiome dataset demonstrates the link between the SmSTRIPAK complex and the PR pathway (Jonkers et al., 2014; Schmidt et al., 2020). In *N. crassa*, HAM-5 functions as the MAP kinase scaffold protein of the PR signaling pathway, showing close associations to each of the kinases from the three tiered cascade, demonstrated by extensive co-localization studies and positive interactions in Y2H experiments (Dettmann et al., 2014; Jonkers et al., 2014; Schmidt et al., 2020). Beyond this, an interaction between the STRIPAK complex and other components of the PR signaling pathway is supported by GFP-trap experiments using the three kinases of the MAK-2 cascade in *N. crassa* as bait. This GFP-trap identified the STRIPAK subunits HAM-3 (PRO11), MOB-3 (SmMOB3), PP2A-A (SmPP2AA) and PPG-1 (SmPP2Ac1). However, the STRIPAK subunits were only identified with low sequence coverage or not in all replicates (Dettmann et al., 2014). While this suggests a transient interaction or possibly a false positive hit, further experiments demonstrated that the STRIPAK complex and PR signaling are tightly interconnected through reciprocal regulation of phosphorylation and direct protein-protein interactions. Biochemical *in vitro* experiments showed phosphorylation of the STRIPAK complex subunit MOB-3 at its N-terminus by MAK-2. Additionally an *in vivo* interaction of the STRIPAK with the PR signaling pathway was demonstrated by capture of MAK-2 in co-IP experiments under mild washing conditions using the PRO11 homolog HAM-3 as bait (Dettmann et al., 2013). Phosphoproteomic studies in *S. macrospora* identified two phosphorylation sites of HAM5 that were differentially regulated in SmSTRIPAK deletion mutants Δpro11, Δpp2Ac1Δpro22 and Δpro11Δpro22, thus suggesting HAM5 as a substrate of the SmSTRIPAK phosphatase activity (Stein et al., 2020).

Collectively, this SCI1-proxiome dataset provides additional *in vivo* evidence for previously reported association of the SmSTRIPAK complex with the SIN and the PR pathways. Additionally, it proposes a role of the SmSTRIPAK complex in ribosome biogenesis by interaction with the HRR25-LTV1 circuit. These findings emphasize the STRIPAK’s role as a signaling hub in fungi and demonstrate the power of the BioID methodology to identify biologically relevant protein environments *in vivo*.

### DOC proteins regulate sexual development in homothallic and heterothallic ascomycetes

Besides its significant enrichment as the most promising candidate in the SCI1 proxiome dataset, SmDOC2 was also recovered in previous tandem affinity purification (TAP) experiments using PRO45 as bait, where it was identified with 66 spectral counts. Thereby ranking SmDOC2 the 17^th^ out of the total 580 proteins when sorted by cumulative spectral counts of the three replicates. In this TAP-MS experiment, other SmSTRIPAK components were identified as follows: PRO11 (164 counts), SCI1 (81 counts) and SmMOB3 (22 counts) (Nordzieke et al., 2015). Additionally, SmDOC1 was identified in two of three SCI1-eGFP pulldown replicates (Reschka, 2018). The identification of SmDOC1/2 proteins as potential SmSTRIPAK-interacting proteins proposes an intriguing link between two fungal regulatory systems, and we decided to generate *Smdoc1/2* knockout strains to further investigate the interaction.

When assessing self-communication during germling fusion, Heller and colleagues reported a non-additive pattern of the *doc* knockout phenotypes in the heterothallic ascomycete *N. crassa* (Heller et al., 2016). The single knockouts Δ*doc-1* and Δ*doc-2* showed reduced self-communication, while the double knockout Δ*doc-1*Δ*doc-2* showed wild-type levels of communication, implying that the presence of only one DOC protein disrupts balanced allorecognition. Our results extend this model and demonstrate that the DOC system is not only restricted to heterothallic species, but also regulates interactions in the homothallic *S. macrospora*. Using the ΔSmdoc1, ΔSmdoc2 and ΔSmdoc1ΔSmdoc2 deletions strains of *S. macrospora,* we showed that *Smdoc1* and *Smdoc2* play critical roles in sexual development. While single deletions of *Smdoc1* or *Smdoc2* were severely impaired in fruiting body formation, the double deletion mutant ΔSmdoc1ΔSmdoc2 was not impaired and displayed wild-type fertility (Fig 3). Since earlier studies of the DOC system in *N. crassa* focused on germling communication, we analyzed the *N. crassa doc* knockouts and identified an impairment in sexual reproduction. The *N. crassa* Δ*doc-2* deletion strain is delayed in protoperithecia formation, and the protoperithecia, which were formed after prolonged incubation, could not be fertilized with conidia of the wild type or the other *doc* knockouts. This suggests that *doc-2* plays a critical, non-redundant role in female, but not male fertility in heterothallic fungi. Despite their differences in mating systems, both *N. crassa* (heterothallic) and *S. macrospora* (homothallic), exhibit overlaps in the characteristic non-additive phenotypes of single *doc* knockout strains. This implies a conserved mechanism, requiring balanced DOC interactions for proper sexual development.

### SmDOC1 is associated with the spatial environment of the septal pore

To determine the subcellular localization, we generated *Smdoc1/2* fusions tagged with fluorescent proteins and subjected to fluorescence microscopy. However, the localization of SmDOC2 remains elusive since *egfp*- and *TagRFP-T* tagged variants of *Smdoc2* could not be localized in fluorescence microscopy and weak signals could be attributed to wild-type autofluorescence. Additionally, the proximity labeling data does not reveal any obvious spatial environment, since DSK1, SIR2, SmMOB3, MEK2 and MAK2 localize to the nucleus, the nuclear envelope, the cytoplasm or septa in *S. macrospora* or other fungal species (Cai et al., 2022; Frey et al., 2015a; Schmidt et al., 2020; Takeuchi & Yanagida, 1993; Tang et al., 2012). Previous studies localized *N. crassa* DOC-2 to the hyphal membrane and septa (Heller et al., 2016).

Fluorescence microscopy of SmDOC1-TagRFP-T showed a localization to a ring-like structure around the septal pore *in vivo* (Fig 8). This localization is further supported by the significant enrichment of proteins associated with the septal pore in the SmDOC1 proxiome. One example is the putative septal pore associated protein SMAC_05428. The *N. crassa* homolog of SMAC_05428, NCU01984 was predicted to be a septal pore associated protein based on mass spectrometry analysis of Woronin body associated protein and bioinformatics approaches that compared the composition and characteristics of known septal proteins (Lai et al., 2012). Moreover, the protein kinase YAK1 was significantly enriched in the SmDOC1 proxiome. Characterization of YAK1 orthologs filamentous fungi including *Candida albicans*, *Fusarium graminearum*, *Botrytis cinerea*, *Magnaporthe oryzae* and *A. fumigatus* have asserted its role in hyphal growth, conidiation and stress response to oxidative stressors (Goyard et al., 2008; Han et al., 2015; Wang et al., 2011; Yang et al., 2018). The *A. fumigatus* YakA ortholog localized to the center of septa and is involved in plugging of fungal septa upon exposure to stress. Additionally, the ΔyakA deletion strain exhibited a penetration defect into solid substrates and was unable to grow under iron-limiting conditions. Analysis of ΔYakA mutant phosphoproteome showed phosphorylation changes for proteins involved in septal formation, including the SO scaffolding protein of the MAK1 signalling pathway (Fleissner & Glass, 2007; van Rhijn et al., 2024). The *S. macrospora* YAK1 was previously identified in pulldown experiments with SCI1-eGFP (Reschka, 2018). Consistent with the localization of SmDOC1-TagRFP-T and BioID enrichment of septal pore associated proteins such as SMAC_05428 and YAK1, the PR components MIK2, MEK2, MAK2 and HAM5 were shown to localize to septal pores in *S. macrospora* (Schmidt et al., 2020). This further underlines the role of SmDOC1 as a protein that localizes to the spatial environment of the septal pore.

### SmDOC1/2 closely interact with PR signaling components

Proximity labeling experiments using SmDOC1 or SmDOC2 as bait showed consistent significant enrichment of components of the MAK2 MAPK cascade (Fig 9). The PR component HAM5 was captured only by SmDOC1, MAK2 only by SmDOC2, whereas MEK2 was significantly enriched using both SmDOC proteins as bait. Moreover, our proximity labeling data showed significant reciprocal enrichment of SmDOC2 when using SmDOC1 as bait, and vice versa, indicating a direct or closely associated interaction between the two SmDOC proteins. Since biotinylation is a highly specific post-translational modification, mainly restricted to histones or carboxylases (Cronan, 1990; Kothapalli et al., 2005; Zempleni et al., 2009), the capture of biotinylated peptides of SmDOC1, HAM5 and MEK2 poses additional evidence indicating artifact-free data and true positive hits.

**Fig 9.**
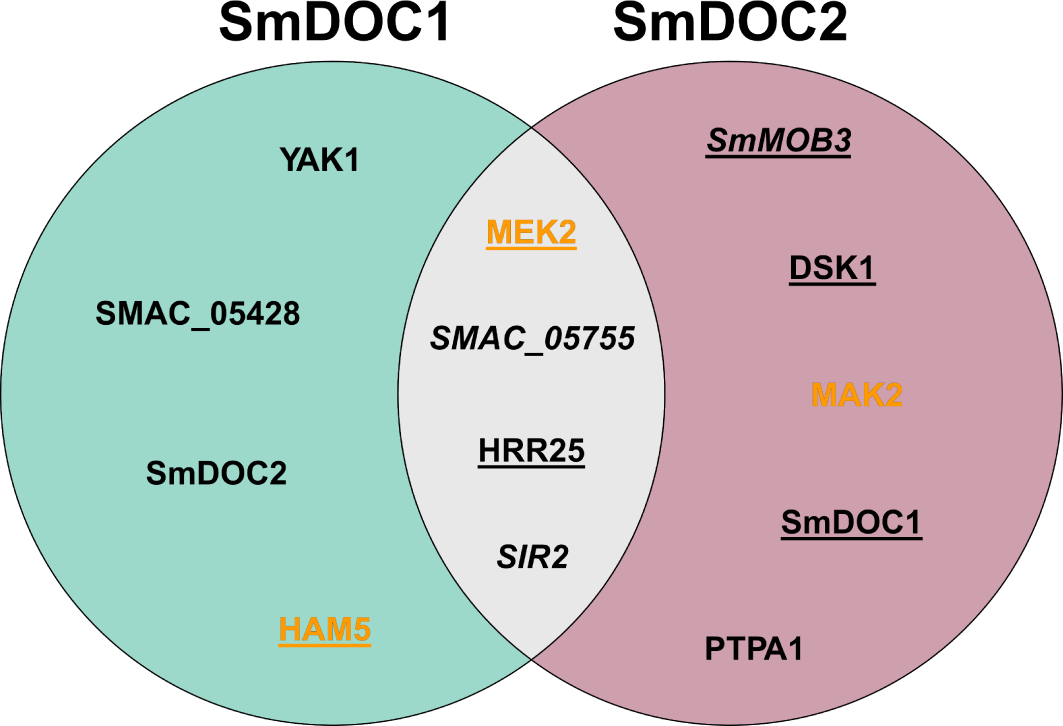
Venn diagram of the SmDOC1/2 proxiomes. Venn diagram showing the significantly enriched proteins from the BioID using SmDOC1 (left, Fig 5) and SmDOC2 (right, Fig 6) as bait. Proteins that were identified with either bait protein are drawn to the overlapping region in the middle (grey). Underlined proteins were identified with biotin site information. Components of the PR signaling pathway are highlighted in orange and proteins that are associated with PR signaling in literature, are written in italics. For details see S9 and S10 Tables.

The close ties of SmDOC1/2 to the PR signaling pathway are also reflected by the Y2H experiments (Fig 7). Here, SmDOC1 interacted with MEK2-t1, MEK2-t2 and MAK2, but not HAM5. However, the absence of HAM5 interaction might be caused by the incompatibility of fusion of the BD to HAM5. Previous Y2H experiments with *S. macrospora* PR signaling pathway components were only able to demonstrate the well characterized interaction of HAM5 with MEK2 and MAK2 when HAM5 was fused to the AD (Schmidt et al., 2020). Therefore, the BD-HAM5 fusion used here may be non-functional, so the lack of signal cannot be taken as evidence against a possible SmDOC1-HAM5 interaction.

Y2H experiments with SmDOC2 were not successful, since already the positive control AD-ranBPM x BD-SmDOC2 showed drastically reduced viability of the cells, when compared to SmDOC1 (S21D Fig). This indicates potential toxicity effects of SmDOC2 in the Y2H assay.

Beyond the direct interactions in Y2H, the SmDOC1/2 BioID experiments significantly enriched proteins that have been reported to be associated with PR pathway components. Among them, the uncharacterized protein SMAC_05755, the *N. crassa* homolog of NCU02606, was significantly enriched in both SmDOC proxiome datasets. The protein NCU02606 was previously identified in *N. crassa* GFP-trap experiments using the PR kinases MAK-2, MEK-2 and NRC-1 as bait, while being absent from the control samples only containing unfused GFP. However, the ΔNCU02606 deletion strain did not exhibit any impairments in germling communication assays of *N. crassa* (Dettmann et al., 2014). Another example for the close association of SmDOC1/2 with the PR pathway is demonstrated by the enrichment of SIR2, which has functional ties to MEK2 in human cell systems. Downregulation of SIRT2 (homolog of *S. macrospora* SIR2) in human cells caused increased acetylation of the MAPKK MEK1 (homolog of *S. macrospora* MEK2) and resulted in hyperactivation of extracellular signal regulated kinases (ERKs) 1/2 (homolog of *S. macrospora* MAK2). Mutant MEK1 variants mimicking constant acetylation showed similar effects as constitutively active phospho-mimic mutants characterized by inappropriate cell growth and proliferation. Immunoprecipitation experiments demonstrated the direct protein-protein interaction of MEK1 and ERK1/2 with SIRT2 (Cha et al., 2021; Yeung et al., 2014). Interestingly, a feedback loop was discovered in which ERK1/2 activation increased protein levels of SIRT2, its protein stability and its deacetylase activity (Choi et al., 2013). While it is not clear whether the PR pathways associated proteins SMAC_05755 or SIR2 are functionally connected to SmDOC1/2, their co-capture in the proxiome dataset demonstrates that SmDOC1/2 occupy the same molecular neighborhood *in vivo*.

### SmDOC1/2 might aid in disassembly of the PR signaling complex

Taking all results into consideration, we postulate the following fundamental mechanistic principles for the SmDOC system. 1) Due to the paradoxical and non-additive phenotypes of *doc* single deletion strains in *S. macrospora* and *N. crassa*, SmDOC1/2 must exhibit an inhibitory effect on sexual development, rather than an activating stimulus. Hence, the double deletion strain ΔSmdoc1ΔSmdoc2, which does not encode any *Smdoc* genes, is not impaired in sexual development. 2) SmDOC1 and SmDOC2 are not redundant in their function. 3) A mutual antagonism mechanism regulates the activity of SmDOC1 and SmDOC2. If only one of the components is absent, the mutual inhibitory effect on the remaining SmDOC protein (e.g. SmDOC1 in ΔSmdoc2 or SmDOC2 in ΔSmdoc1) is lifted, which in turn leads to the suppression of sexual development by the remaining SmDOC protein. Alternatively, the proteins could function in competing pathways where their balanced activity is crucial for normal development, and the complete absence of both pathways allows for the engagement of alternative developmental mechanisms. Based on the reciprocal capture of SmDOC1 in the proximity labeling data of SmDOC2-TurboID, and vice versa, we suggest that the SmDOC proteins might interact with each other, possibly by forming a heterodimer. 4) The proximity enrichment of MEK2, MAK2, HAM5 and proteins associated with the PR complex, direct interaction of SmDOC1 with MEK2 and MAK2 in Y2H assays and overlapping subcellular localization in fluorescence microscopy, all converge on a model in which SmDOC1/2 operate in close proximity and directly interact with the PR signaling pathway. We speculate that SmDOC1/2 might assist in disassembly of the PR signaling complex to regulate sexual development in *S. macrospora* (Fig 10). The inactive system is characterized by reciprocal inhibition of SmDOC1 and SmDOC2. Upon chemotropic interactions, the PR MAP kinase cascade of MIK2, MEK2, MAK2 and the scaffold HAM5 assembles, and phosphorylation of the kinase cascade takes place. Once the terminal kinase, MAK2 is phosphorylated, it translocates to the nucleus, altering gene expression by the activation of transcription factors (Fischer et al., 2018; Leeder et al., 2013). During this step, active MAK2 phosphorylates HAM5, initiating a negative feedback mechanism, which leads to disassembly of the MAPK complex (Jonkers et al., 2014). We speculate that at this stage SmDOC1/2 are activated, e.g. by phosphorylation through MAK2 or another kinase, which lifts their reciprocal repression cycle.

**Fig 10.**
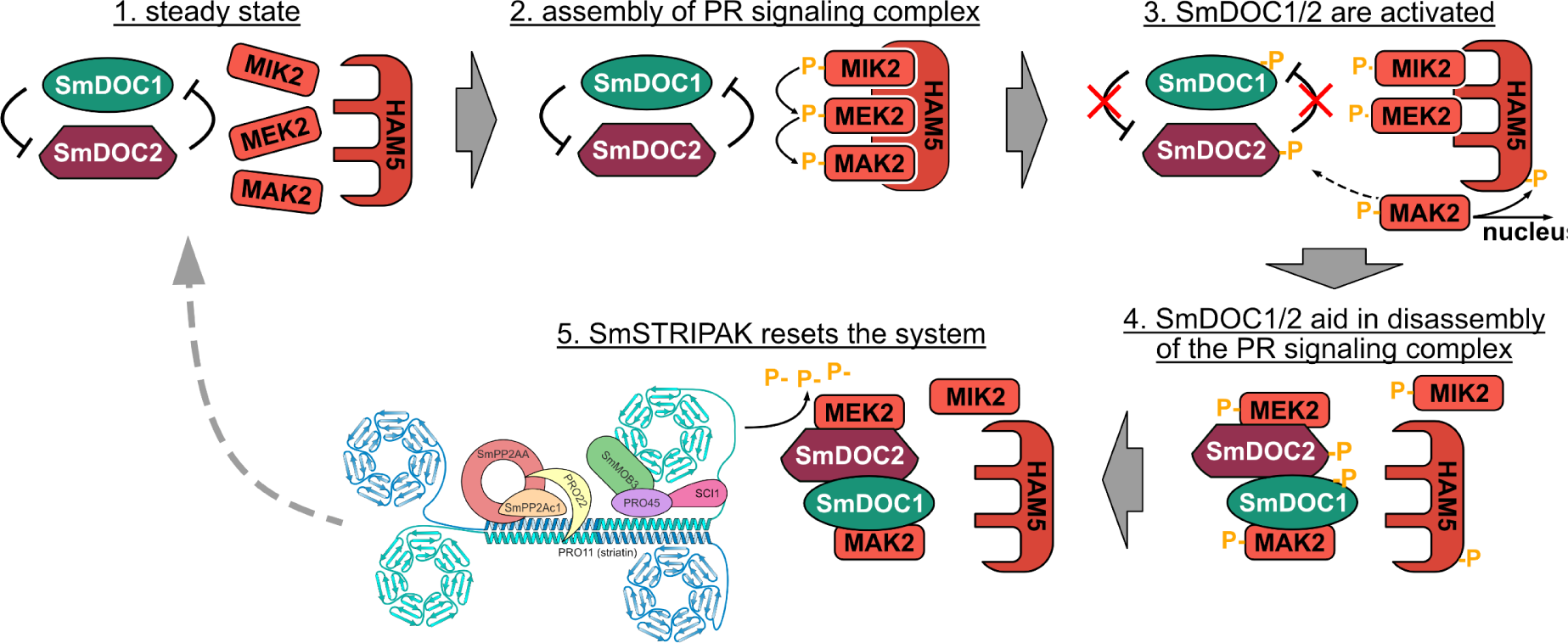
Mechanistic model of SmDOC1/2 in a homothallic interaction. This model describes the mechanistic SmDOC1/2 system in *S. macrospora*. We speculate that SmDOC1/2 might regulate PR complex assembly through a cyclic mechanism. 1) In the inactive state SmDOC1 and SmDOC2 suppress each other’s activity by direct or indirect reciprocal inhibition. 2) Upon chemotropic signaling, the PR MAP kinase cascade (MIK2/MEK2/MAK2/HAM5) assembles, and phosphorylation is passed downstream along the cascade. 3) We propose that MAK2 or associated kinases phosphorylate and activate SmDOC1/2, releasing their reciprocal repression. 4) Activated SmDOC1/2 bind directly to the PR components, MEK2, MAK2 and HAM5, facilitating complex disassembly. 5) The SmSTRIPAK complex resets the cycle through phosphatase activity, dephosphorylating SmDOC1/2 and PR components, returning the system to its initial state.

Once SmDOC1/2 are activated, they might bind to MEK2, MAK2 and HAM5, as indicated by the proximity labeling data and the Y2H experiments, thereby aiding in the disassembly of the PR signaling complex. In the last step, the SmSTRIPAK could reset the cycle by its phosphatase activity, removing the phosphorylation of SmDOC1/2 and possibly dephosphorylating components of the PR pathway. The system returns to its initial state, characterized by the mutual antagonism of SmDOC1 and SmDOC2, as well as non-assembled PR signaling complex.

Since our experiments in *S. macrospora* were performed in homokaryotic cultures, lacking any allelic variety of *Smdoc1/2*, our results and therefore this model reflect a compatible allorecognition interaction. When extending our model to incompatible interactions, we envision that SmDOC1/2 might be regulated by a two-step activation mechanism through multi-site phosphorylation. The sensing of an allelic incompatibility through a receptor could lead to additional phosphorylation and activation of SmDOC1/2, further increasing the inhibitory activity on the PR complex. The regulatory potential of SmDOC1 and SmDOC2 is demonstrated by the high number of phosphosites that were identified in the proximity labeling data (14 and 9 distinct phosphosites, respectively). Multi-phosphorylated SmDOC1/2 might bind PR components more tightly without releasing them. This would result in a situation of competitive inhibition, which prevents the assembly of the MAPK signaling cascade and thereby suppresses oscillation. However, this theory of SmDOC1/2 as phosphorylation-gated inhibitors requires extensive experimental validation of the biological relevance of the individual phosphosites.

## Methods

### Generation of plasmids

All plasmids and primers used in this study are listed in S1 and S2 Tables. Cloning and propagation of plasmids was performed with *Escherichia coli* MACH1 (Thermo Fisher Scientific). Plasmids were assembled using homologous recombination in *Saccharomyces cerevisiae* (Colot et al., 2006) or using the NEBuilder^®^ HiFi DNA Assembly kit (New England Biolabs GmbH, E2621S).

### Generation of knockout strains ΔSmdoc1, ΔSmdoc2 and ΔSmdoc1ΔSmdoc2

Primers used for the construction of plasmids are listed in S2 Table and strains generated during this study are listed in S3 Table. The *Smdoc* knockouts were constructed by replacing the respective open reading frames (ORFs) with the hygromycin resistance cassette (*hyg^R^*) using homologous recombination in the Δku70 background. For construction of the knockout plasmids pdoc1-KO, pdoc2-KO and pdoc1+2-KO the 1 kb 5’ and 3’ flanking regions were amplified from *S. macrospora* wild type (DSM997) gDNA and contain 29 bp overhangs to the pRS426 vector (Christianson et al., 1992) or the hygromycin resistance cassette (*hyg^R^*). The 1 kb 5’ flanks were amplified using primer pairs LH01/LH02 (pdoc1-KO and pdoc1+2-KO) and LH05/LH06 (pdoc2-KO). The 1 kb 3’ flanks were amplified with primer pairs LH03/LH04 (pdoc1-KO), LH07/LH08 (pdoc2-KO) and LH12/LH13 (pdoc1+2-KO). The *hyg^R^* cassette (1418 bp) expresses the *hygromycin B phosphotransferase* gene (*hph*) of *Escherichia coli* under control of the constitutive *trpC* promoter from *Aspergillus nidulans* and was amplified from plasmid pSmnbr1-KO (Werner et al., 2019) with the primer pair hph-f/hph-r. The fragments were integrated into *Xho*I linearized pRS426 by homologous recombination in *S. cerevisiae* strain PJ64-4A (Colot et al., 2006; James et al., 1996). The deletion cassettes consisting of the 1 kb flanking regions and the *hyg^R^* cassette were amplified from the sequenced plasmids using the primers LH01/LH04 (ΔSmdoc1), LH05/LH08 (ΔSmdoc2); LH01/LH13 (ΔSmdoc1ΔSmdoc2). The PCR amplicons were desalted and transformed into the nourseothricin-resistant *S. macrospora* Δku70 deletion strain for homologous recombination (Pöggeler & Kück, 2006). In the Δku70 deletion strain, the *ku70* gene was replaced with the nourseothricin resistance cassette (*nat^R^*) expressing the *nourseothricin acetyltransferase* gene of *Streptomyces noursei* under control of the constitutive *trpC* promoter from *A. nidulans*. The wild-type *ku70* gene was restored by crossing the primary transformants of the *Smdoc* knockouts with spore color mutant fus1-1 and isolating hygromycin resistant, but nourseothricin sensitive spores. These single spore isolates were genotyped using PCR with one of the primers binding outside of the 1 kb flanks within the genomic region to verify the 5’ and 3’ junctions of the *hyg^R^* integration at the respective *Smdoc* locus. Additionally, the absence of the respective wild-type *Smdoc* ORF was verified by PCR. Southern hybridization was performed to rule out off target integrations of the *hyg^R^* cassette.

### Light and fluorescence microscopy

For microscopic analyses the *S. macrospora* strains were grown on sterile microscopy slides that were covered in BMM medium. The slides were inoculated with agar plugs containing mycelium. The slides were grown for 1-2 days at 27 °C under constant light conditions. The slides were analyzed using an Axio Imager M1 microscope (Zeiss, Germany). For detection of EGFP signal (termed green channel) the Chroma filter set 49002 was used and TagRFP-T signals (referred to as red channel) were acquired using the Chroma filter set 49005. Pictures were captured with the Axiocam 503 camera (Zeiss, Germany). Magnifications of fungal colonies for phenotypic analyses were taken with a VHX-500F Digital Microscope (Keyence, Japan).

### Protein extraction and Western blot hybridization

In brief, the *S. macrospora* strains were grown in large Petri dishes (⌀ = 150 mm) filled with 50 ml of liquid medium. The medium was inoculated with 5-7 solid agar plugs from precultures. The strains were grown for three to four days at 27 °C under constant light conditions. Afterwards, the agar plugs were removed, the mycelium was pressed dry and ground in liquid nitrogen. Proteins were extracted from the mycelium powder using the lysis buffer 10 mM Tris pH 7.5, 150 mM NaCl, 0.5 mM EDTA pH 8, 1 mM PMSF, 2 mM DTT, 0.5 % Nonidet-P40, 2x cOmplete™ EDTA-free Proteinase Inhibitor Cocktail (Roche, Switzerland), 1x PhosSTOP™ (Roche, Switzerland) followed by centrifugation for 20 min at 10 000 x g. Roughly 50 µg of protein were loaded onto a SDS polyacrylamide gel electrophoresis (PAGE). Blotting was performed with the Trans-Blot Turbo Transfer System (BioRad) using the Trans-Blot Turbo RTA Mini 0.2 µm Nitrocellulose Transfer Kit according to the manufacturer’s instructions. After blotting, reversible Ponceau S staining (0.1 % Ponceau S in 5% glacial acetic acid) was used to stain total protein on the membrane to assess the loading. Biotinylated proteins were detected with a Streptavidin-HRP conjugate (Thermo Scientific, 21130, 1:30000). Membranes were incubated with 500 µl WesternBright ECL HRP (Advansta Corporation, USA) substrate and documented with the digital Lourmat FUSION SL (Vilber, France) imaging system.

### BioID sample preparation

BioID experiments were performed as described in Hollstein et al. (2025). For BioID experiments, the proteins were extracted from cell lysates at denaturing conditions (4 % SDS) including an incubation for 5 min at 65 °C. Biotinylated proteins were enriched using 100 µl slurry of Strep-Tactin^®^ Sepharose^®^ (IBA Lifesciences GmbH, 2-1201-002) per 1 ml of protein crude extract according to the manufacturer’s instructions for batch purification. Biotinylated proteins were released from the Strep-Tactin^®^ Sepharose^®^ resin by competitive elution using the biotin-containing buffer BXT (0.1 M Tris-Cl, 0.15 M NaCl, 1 mM EDTA, 50 mM biotin, pH 8). The BXT-eluate was purified by chloroform methanol precipitation (Wessel & Flügge, 1984). The protein pellet was resuspended in 1x SDS loading dye and separated on a polyacrylamide gel. Proteins were digested in-gel by trypsin and the eluted peptides were desalted with C18 StageTips prior to LC-MS analysis (Rappsilber et al., 2003; Shevchenko et al., 1996).

### LC-MS analysis

Processed BioID samples were analyzed with LC-MS by the Service Unit LC-MS Protein Analytics of the Göttingen Center for Molecular Biosciences (GZMB) of the University of Göttingen. Dried peptide samples were reconstituted in 20 µl LC-MS sample buffer (2 % acetonitrile, 0.1 % formic acid). 2 to 8 µl of each sample were subjected to reverse phase liquid chromatography for peptide separation using an RSLCnano Ultimate 3000 system (Thermo Fisher Scientific): Peptides were loaded on an Acclaim PepMap 100 pre-column (100 μm x 2 cm, C18, 5 μm, 100Å; Thermo Fisher Scientific) with 0.07 % trifluoroacetic acid at a flow rate of 20 µl/min for 3 min. Analytical separation of peptides was done on an Acclaim PepMap RSLC column (75 μm x 50 cm, C18, 2 μm, 100Å; Thermo Fisher Scientific) at a flow rate of 300 nL/min. The solvent composition was gradually changed within 94 min from 96 % solvent A (0.1 % formic acid) and 4 % solvent B (80 % acetonitrile, 0.1 % formic acid) to 10 % solvent B within 2 min, to 30 % solvent B within the next 58 min, to 45 % solvent B within the following 22 min, and to 90 % solvent B within the last 12 min of the gradient. All solvents and acids had Optima grade for LC-MS (Fisher Chemical). Eluting peptides were on-line ionized by nano-electrospray (nESI) using the Nanospray Flex Ion Source (Thermo Fisher Scientific) at 1.5 kV (liquid junction) and transferred into a Q Exactive HF mass spectrometer (Thermo Fisher Scientific). Full scans in a mass range of 300 to 1650 m/z were recorded at a resolution of 30,000 followed by data-dependent top 10 HCD fragmentation at a resolution of 15,000 (dynamic exclusion enabled). LC-MS method programming and data acquisition was performed with the XCalibur 4.0 software (Thermo Fisher Scientific). The LC-MS raw files were analyzed using MaxQuant version 1.6.10.43 and the detailed configuration is documented in the “mqpar.xml” file provided in the uploaded MaxQuant search results folder. Due to their vastly different nature in protein content and distribution, proteome controls were configured in a different parameter group than BioID eluate samples and the option “Separate LFQ in parameter groups” was activated in MaxQuant. This avoids the cross-quantification of BioID eluate samples with their non-enriched input controls, which reflect the whole proteome. Therefore, LFQ intensities for BioID eluates are exclusively quantified with other BioID eluate samples. The *S. macrospora* protein sequences from version 4 were used as sequence input into MaxQuant (Breuer et al., 2024). The data processing steps of LC-MS raw data with MaxQuant and the statistical evaluation with Perseus version 1.6.15.0 are listed in Table 1. The naming scheme of the LC-MS raw files is listed S5 Table.

**Table 1.**
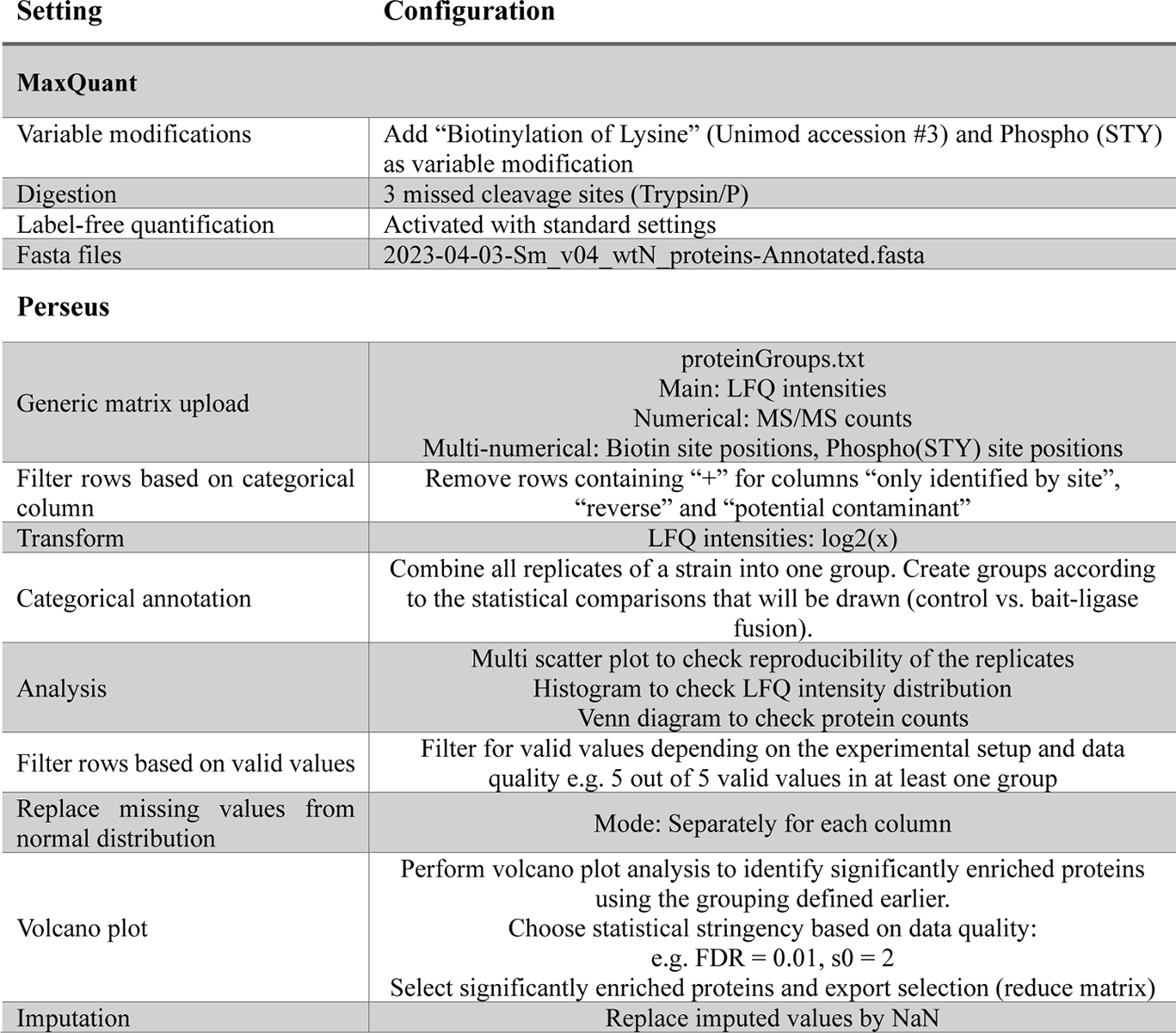
Data processing steps of LC-MS raw files. LC-MS raw files were processed with MaxQuant version 1.6.10.43 and statistical evaluation was performed with Perseus version 1.6.15.0.

## Acknowledgements

We thank our technicians Gertrud Stahlhut and Ulrike Brandt for their technical support. We also thank Dr. Ines Teichert for providing PR pathway Y2H plasmids. We thank our students Frauke Liesegang and Katharina Bornemann for their contribution to this project. This work has been funded by the Deutsche Forschungsgemeinschaft, PO 523/10-1 project number 538832008, INST 186/1230-1 FUGG and INST 186/1465-1.

## Data availability

The mass spectrometry proteomics data have been deposited to the ProteomeXchange Consortium via the PRIDE partner repository (Perez-Riverol et al., 2022) with the dataset identifiers PXD069198 and PXD069539.

## Declaration of interests

The authors declare that they have no known competing financial interests or personal relationships that could have appeared to influence the work reported in this paper.

## Author contributions (CRediT statement)

**Lucas S. Hollstein:** Project administration, Conceptualization, Methodology, Formal analysis, Investigation, Data Curation, Visualization, Methodology, Writing - Original Draft, Writing - Review & Editing

**Kerstin Schmitt:** Investigation, Methodology, Resources, Writing - Review & Editing

**Lucas Well:** Investigation, Writing - Review & Editing

**André Fleißner:** Investigation, Conceptualization, Writing - Review & Editing

**Oliver Valerius:** Investigation, Methodology, Resources, Writing - Review & Editing, Funding acquisition

**Stefanie Pöggeler:** Supervision, Conceptualization, Resources, Writing - Review & Editing, Funding acquisition

## Notes

### Competing Interest Statement

The authors have declared no competing interest.

