## Supplementary Material for "Capture of the SmSTRIPAK proxiome identifies the greenbeard proteins SmDOC1/2 as regulators of the MAK2 pathway to control sexual development in *Sordaria macrospora*"

Generation of plasmids

The plasmids used in this study are listed in S2 Table. The plasmid pHRSmdoc1-TurboID was cloned by amplifying the *Smdoc1* 1 kb 5’ flank with primer pair LH88/LH89 from *S. macrospora* wild-type gDNA, the *Smdoc1*-ORF with primer pair LH90/91 from *S. macrospora* wild-type gDNA, the *TurboID+TtrpC* sequence with primer pair LH98/LH93 from p5'-sci1-L-TurboID, the $nat^R$ with primer pair LH94/95 from pRS_nat, and the *Smdoc1* 1 kb 3’ flank with primer pair LH96/LH97 from *S. macrospora* wild-type gDNA. The fragments were integrated into *Not*I and *EcoR*I linearized pRS426 using the NEBuilder® HiFi DNA Assembly kit (New England Biolabs GmbH, E2621S) according to the manufacturer’s instructions.

The plasmid pHRSmdoc1-TagRFP-T was cloned by amplifying the *Smdoc1* 1 kb 5’ flank with primer pair LH88/LH89 from *S. macrospora* wild-type gDNA, the *Smdoc1*-ORF with primer pair LH90/91 from *S. macrospora* wild-type gDNA, the *TagRFP-T+TtrpC* sequence with primer pair LH92/LH93 from p5’vac14-TagRFP-T, the $nat^R$ with primer pair LH94/95 from pRS_nat, and the *Smdoc1* 1 kb 3’ flank with primer pair LH96/LH97 from *S. macrospora* wild-type gDNA. The fragments were integrated into *Not*I and *EcoR*I linearized pRS426 using the NEBuilder® HiFi DNA Assembly kit (New England Biolabs GmbH, E2621S) according to the manufacturer’s instructions. The plasmid pHRccg1Smdoc1-TagRFP-T was generated to put the *Smdoc1-TagRFP-T* fusion under control of a constitutive promoter. Therefore, pHRSmdoc1-TagRFP-T was linearized by *EcoR*I and a fragment containing the constitutive *ccg1* promoter was amplified from pc-sci1-TurboID using the primer pair LH109/LH110 and inserted in between the *Smdoc1* 1 kb 5’ flank and the start codon of *Smdoc1*.

The plasmid p5’Smdoc2-L-TurboID for ectopic integration was cloned by amplifying the *Smdoc2* promoter and *Smdoc2* ORF from *S. macrospora* wild-type gDNA with primers LH05/LH09 and the sequences for TurboID with the 3x HA Tag and T*trpC* terminator were amplified from p5'-sci1-L-TurboID using primers TtrpC_pRS_r/ SmtBioID-L-f-2. Both fragments were cloned into *Xho*I linearized pRS_nat using the NEBuilder® HiFi DNA Assembly kit (New England Biolabs GmbH, E2621S) according to the manufacturer’s instructions.

For construction of the plasmid pHRSmdoc2-TurboID the *S. macrospora Smdoc2* promoter, *Smdoc2*-ORF and TurboID with the 3x HA Tag and T*trpC* terminator were amplified from plasmid p5’Smdoc2-L-TurboID with primers LH05/LH37. The $nat^R$ was amplified from plasmid pRS_nat using primers LH38/nat-1r. The 3’ flank of *Smdoc2* was amplified from *S. macrospora* wild-type gDNA with primers LH36/LH08. The resulting fragments were integrated into *Xho*I linearized pRS426 using the NEBuilder® HiFi DNA Assembly kit (New England Biolabs GmbH, E2621S) according to the manufacturer’s instructions.

The plasmid pHRSmdoc2-egfp was cloned by amplifying the *Smdoc2* 1 kb 5’ flank with primer pair LH61/62 from *S. macrospora* wild-type gDNA, the *Smdoc2*-ORF with primer pair LH63/64 from *S. macrospora* wild-type gDNA, the *egfp-TtrpC* sequence with primer pair LH65/66 from template p1783-1 and the $nat^R$ fused to the Smdoc2 1 kb 3’ flank with primer pair LH67/68 from template pHRSmdoc2-TurboID. The fragments were integrated into *Not*I linearized pRS426 using the NEBuilder® HiFi DNA Assembly kit (New England Biolabs GmbH, E2621S) according to the manufacturer’s instructions. The plasmid pHRccg1Smdoc2-egfp was generated to put the *Smdoc2-egfp* fusion under control of a constitutive promoter. Therefore, pHRSmdoc2-egfp was linearized by *EcoR*I and *Bgl*II and a fragment containing the constitutive *ccg1* promoter was amplified from pc-sci1-TurboID using the primer pair LH86/LH87 and inserted in between the *Smdoc2* 1 kb 5’ flank and the start codon of *Smdoc2* using the NEBuilder® HiFi DNA Assembly kit (New England Biolabs GmbH, E2621S) according to the manufacturer’s instructions.

The plasmid pHRSmdoc2-TagRFP-T was cloned by amplifying the *Smdoc2* 1 kb 5' flank and the *Smdoc2* ORF with primer pair LH111/112 from pHRSmdoc2-TurboID, the *TagRFP-T+TtrpC+nat*$^R$ sequence with primer pair LH113/114 from pHRSmdoc1-TagRFP-T and the *Smdoc2* 1 kb 3' flank with primer pair LH115/116 from pHRSmdoc2-TurboID. The fragments were integrated into *Not*I linearized pRS426 using the NEBuilder® HiFi DNA Assembly kit (New England Biolabs GmbH, E2621S) according to the manufacturer's instructions. For expression of the *Smdoc2-TagRFP-T* fusion under control of the *ccg1* promoter in pHRccg1-Smdoc2-TagRFP-T, the *Smdoc2* 1 kb 5' flank, the *ccg1* promoter and the *Smdoc2* ORF were amplified with primer pair LH111/112 from template pHRccg1Smdoc2-egfp. The fragments were integrated into *Not*I linearized pRS426 using the NEBuilder® HiFi DNA Assembly kit (New England Biolabs GmbH, E2621S) according to the manufacturer's instructions.

Yeast Two-Hybrid

Yeast Two-Hybrid (Y2H) experiments were performed based on the Matchmaker® Gold Yeast Two-Hybrid system (Takara Bio USA). The plasmids and strains used for Y2H experiments are listed in S2 and S3 Table. The plasmids were constructed using the supplied vectors pGADT7 containing the Gal4 activation domain (AD), and pGBKT7 containing the Gal4 DNA-binding domain (BD). The coding sequences of *Smdoc1*(2523 bp, AD LH125/126 or BD LH127/LH128), *Smdoc2* (2646 bp, AD LH129/LH130 or BD LH131/LH132), *mek2-t1* (1160 bp and 123 bp fragments, AD LH119/LH120 and LH121/LH122, BD LH123/LH120 and LH121/LH124), *mek2-t2* (1160 bp and 438 bp fragments, AD LH119/LH137 and LH138/LH139, BD LH123/LH137 and LH138/LH140) and *ham5* (5139 bp, AD LH133/LH134, BD LH135/LH136) were amplified from *S. macrospora* wild-type cDNA and integrated into *EcoR*I linearized pGADT7 or pGBKT7 via the NEBuilder® HiFi DNA Assembly kit (New England Biolabs GmbH, E2621S). The plasmids pAD-mak2 and pBD-mak2 were kindly donated by Schmidt et al. (2020). The bait plasmids pGBKT7, pBD-Smdoc1, pBD-Smdoc2, pBD-mek2-t1, pBD-mek2-t2, pBD-ham5 and pBD-mak2 were transformed into *S. cerevisiae* strain Y2HGold (MATa, Takara Bio) and transformants were selected on media lacking tryptophan. The prey plasmids pGADT7, pAD-Smdoc1, pAD-Smdoc2, pAD-mek2-t1, pAD-mek2-t2, pAD-ham5, pAD-mak2 and pAD-ranBPM were transformed into strain Y187 (MATα, Harper et al., 1993) and selected for leucine prototrophy. Recombinant bait and prey strains were mated and plated on SC medium lacking tryptophan and leucine to screen for the cells containing both pAD and pBD plasmids. Interactions of bait and prey fusion proteins were analyzed by drop dilution assays. Therefore, the cells were grown in selective SD medium lacking tryptophan and leucine. The cells were diluted to 0.1 optical density (OD) and 25 µL were spotted in 1:10 serial dilutions onto selective SD plates. The plates were incubated at 30 °C for 5 days.

Generation of *S. macrospora* strains

All *S. macrospora* strains used in this study are listed in S3 Table. The selection of recombinant strains was performed by supplementation of the antibiotics hygromycin B (110 U/ml) or nourseothricin (50 µg/ml). Transformation of S. macrospora was carried out as previously described (Walz & Kück, 1995). Primary transformants were crossed to the color spore mutant fus1-1 (Nowrousian et al., 2012) and single spores were isolated from recombinant perithecia. *S. macrospora* was grown in either liquid or solid biomalt maize medium (BMM) or *Sordaria* Westergaard's (SWG) fructification medium at 27 °C under constant light (Esser, 1982; Westergaard & Mitchell, 1947).

**S1 Table. Plasmids used in this study.**

| Plasmid | Characteristics | Reference |
|---|---|---|
| p1783-1 | *amp*$^R$, *ura3, hyg*$^R$ *gpd::egfp:TtrpC* | Pöggeler et al. (2003) |
| p5'vac14-TagRFP-T | *amp*$^R$, *ura3, nat*$^R$ *5'vac14::vac14::TagRFP-T::TtrpC* | Groth et al. (2021) |
| p5'-sci1-L-TurboID | *5'sci1*::sci1-L-TurboID::3xHA::TtrpC | Hollstein et al. (2022) |
| p5'Smdoc2-L-TurboID | *amp*$^R$, *ura3, nat*$^R$ *5'Smdoc2::Smdoc2::L-TurboID::3xHA::TtrpC* | This study |
| pc-sci1-TurboID | *amp*$^R$, *ura3, hyg*$^R$ *ccg1::sci1-TurboID::3xHA::TtrpC* | Hollstein et al. (2022) |
| pHRccg1Smdoc1-TagRFP-T | *amp*$^R$, *ura3, nat*$^R$ *5'Smdoc1::ccg1::Smdoc1::L-TagRFP-T::TtrpC::nat*$^R$*::3'Smdoc1* | This study |
| pHRccg1Smdoc2-egfp | *amp*$^R$, *ura3, nat*$^R$ *5'Smdoc2::ccg1::Smdoc2::L-egfp::TtrpC::nat*$^R$*::3'Smdoc2* (GGGGS linker) | This study |
| pHRccg1Smdoc2-TagRFP-T | *amp*$^R$, *ura3, nat*$^R$ *5'Smdoc2::ccg1::Smdoc2::L-TagRFP-T::TtrpC::nat*$^R$*::3'Smdoc2* | This study |
| pHRSmdoc1-TagRFP-T | *amp*$^R$, *ura3, nat*$^R$ *5'Smdoc1::Smdoc1::L-TagRFP-T::TtrpC::nat*$^R$*::3'Smdoc1* | This study |
| pHRSmdoc1-TurboID | *amp*$^R$, *ura3, nat*$^R$ *5'Smdoc1::Smdoc1::L-TurboID::3xHA::TtrpC::nat*$^R$*::3'Smdoc1* | This study |
| pHRSmdoc2-egfp | *amp*$^R$, *ura3, nat*$^R$ *5'Smdoc2::Smdoc2::L-egfp::TtrpC::nat*$^R$*::3'Smdoc2* (GGGS linker) | This study |
| pHRSmdoc2-TagRFP-T | *amp*$^R$, *ura3, nat*$^R$ *5'Smdoc2::Smdoc2::L-TagRFP-T::TtrpC::nat*$^R$*::3'Smdoc2* | This study |
| pHRSmdoc2-TurboID | *amp*$^R$, *ura3, nat*$^R$ *5'Smdoc2::Smdoc2::L-TurboID::3xHA::TtrpC::nat*$^R$*::3'Smdoc2* | This study |
| pRS_hyg | *amp*$^R$, *ura3, hyg*$^R$ | Bloemendal et al. (2012) |
| pRS_nat | *amp*$^R$, *ura3, nat*$^R$ | Klix et al. (2010) |
| pRS426 | *amp*$^R$, *ura3* | Christianson et al. (1992) |
| pSmdoc1+Smdoc2-KO | *amp*$^R$, *ura3, hyg*$^R$ *5'Smdoc1::hyg*$^R$*::5'Smdoc2* | This study |
| pSmdoc1-KO | *amp*$^R$, *ura3, hyg*$^R$ *5'Smdoc1::hyg*$^R$*::3'Smdoc1* | This study |
| pSmdoc2-KO | *amp*$^R$, *ura3, hyg*$^R$ *5'Smdoc2::hyg*$^R$*::3'Smdoc2* | This study |
| **Yeast Two-Hybrid plasmids** | | |
| pAD-HAM5 | *HAM5* (*SMAC_02471*) cDNA in pGADT7 | Schmidt et al. (2020) |
| pAD-MAK2 | *MAK2* (*SMAC_03492*) cDNA in pGADT7 | Schmidt et al. (2020) |
| pAD-MEK2-t1 | *MEK2-t1* (*SMAC_06526-t1*) cDNA in pGADT7 | Schmidt et al. (2020) |
| pAD-MEK2-t2 | *MEK2-t2* (*SMAC_06526-t2*) cDNA in pGADT7 | This study |
| pAD-ranBPM | *ranBPM* cDNA in pGADT7 | Tucker et al. (2009) |
| pAD-Smdoc1 | *Smdoc1* (*SMAC_06903*) cDNA in pGADT7 | This study |
| pAD-Smdoc2 | *Smdoc2* (*SMAC_06902*) cDNA in pGADT7 | This study |
| pBD-HAM5 | *HAM5* (*SMAC_02471*) cDNA in pGBKT7 | Schmidt et al. (2020) |

| | | |
|---|---|---|
| pBD-MAK2 | *MAK2* (*SMAC_03492*) cDNA in pGBKT7 | Schmidt et al. (2020) |
| pBD-MEK2-t1 | *MEK2-t1* (*SMAC_06526-t1*) cDNA in pGBKT7 | Schmidt et al. (2020) |
| pBD-MEK2-t2 | *MEK2-t2* (*SMAC_06526-t2*) cDNA in pGBKT7 | This study |
| pBD-Smdoc1 | *Smdoc1* (*SMAC_06903*) cDNA in pGBKT7 | This study |
| pBD-Smdoc2 | *Smdoc2* (*SMAC_06902*) cDNA in pGBKT7 | This study |
| pGADT7 | *LEU2, GAL4-AD, amp$^R$, PADH1::GAL4AD::TADH1* | Takara Bio USA |
| pGBKT7 | *TRP1, GAL4-BD, kan$^R$ PADH1::GAL4BD::TADH1* | Takara Bio USA |

*amp$^R$,* ampicillin resistance; ura3, *orotidine-5'-phosphate decarboxylase* gene of *Saccharomyces cerevisiae*; *hyg$^R$*, hygromycin resistance cassette expressing the *hygromycin B phosphotransferase* gene from *Escherichia coli* under control of the constitutive *trpC* promoter from *Aspergillus nidulans; nat$^R$,* nourseothricin resistance cassette expressing the *nourseothricin acetyltransferase* gene from *Streptomyces noursei* under control of the constitutive *trpC* promoter from *A. nidulans*; *ccg1*, constitutive promoter of the *clock controlled gene 1* of *Neurospora crassa; gpd,* constitutive promoter of the *glyceraldehyde-3-phosphate dehydrogenase* gene of *A. nidulans; TtrpC,* terminator of the *anthranilate synthase* gene of *A. nidulans*; *egfp*, gene for the enhanced green fluorescent protein of *Aequorea victoria*; *TagRFP-T*, gene for the red fluorescent protein TagRFP-T of *Entacmaea quadricolor*; P, promoter; T, terminator; L, linker with the sequence GGGGSGGGGS if not indicated otherwise.

**S2 Table. Primers used in this study**

| Name | Sequence |
|---|---|
| h3 | gtactcgccgatagtggaaac |
| hph-f | gttaactgatattgaaggagcatttttgg |
| hph-f (RL76) | ccgcgacgtctgtcgagaag |
| hph-r | gttaactggttcccggtcggcatctactc |
| hph-r (RL77) | cattgtccgtcaggacattg |
| LH01-doc1-ko-5f | gtaacgccagggttttcccagtcacgacgacatcactccgttaaccacc |
| LH02-doc1-ko-5r | ccaaaaatgctccttcaatatcagttaactgtttcccgttccacaaagg |
| LH03-doc1-ko-3f | agagtagatgccgaccgggaaccagttaacaaactcatttcgagcgatt |
| LH04-doc1-ko-3r | gcggataacaatttcacacaggaaacagctcagacgcgctgcaattgca |
| LH05-doc2-ko-5f | gtaacgccagggttttcccagtcacgacgttcatacctcagatcggcac |
| LH06-doc2-ko-5r | ccaaaaatgctccttcaatatcagttaacgatggtggtaataaaacaat |
| LH07-doc2-ko-3f | agagtagatgccgaccgggaaccagttaatgacgatggaatgggaaagg |
| LH08-doc2-ko-3r | gcggataacaatttcacacaggaaacagcccgggcgacgggtgtcaatt |
| LH09-doc2-L-TurboID-r | ccgccgccgccgctgccgccgccgcccattgaaccaagctcaagagtca |
| LH101-d1HR-f | acatcactccgttaaccaccc |
| LH102-d1HR-r | tgcaattgcacaatgcctgatc |
| LH103-d2HR-f | tcatacctcagatcggcacatg |
| LH104-d2HR-r | acgggtgtcaattccttccag |
| LH109-d1-ccg1-f | aacacacaacctttgtggaacgggaaacagaattctagaaggagcagtccatctg |
| LH110-d1-ccg1-r | cgggagagcccttgccgctgctacccatgaattctttggttgatgtgaggggttgtg |
| LH119-AD-MEK2-t1f | tatggccatggaggccagtgaattcatggccgacccattcgcc |
| LH120-MEK2-t1-intr | tttggtacgctgaagtgggtgtaggttcatcgc |
| LH121-MEK2-t1-intf | acccacttcagcgtaccaaaccgacccc |
| LH122-AD-MEK2-t1r | tatcgatgcccacccgggtgctactcgcgctcccagtc |
| LH123-BD-MEK2-t1f | gcatatggccatggaggccgaattcatggccgacccattcgcc |
| LH124-BD-MEK2-t1r | gcaggtcgacggatccccggctactcgcgctcccagtc |
| LH125-AD-SmDOC1-f | tatggccatggaggccagtgaattcatgggtagcagcggcaag |
| LH126-AD-SmDOC1-r | tatcgatgcccacccgggtgtcaagcaataggcaaatccatacc |
| LH127-BD-SmDOC1-f | gcatatggccatggaggccgaattcatgggtagcagcggcaag |

| | |
|---|---|
| LH128-BD-SmDOC1-r | gcaggtcgacggatccccggtcaagcaataggcaaatccatacc |
| LH129-AD-SmDOC2-f | tatggccatggaggccagtgaattcatgatggcctccgcgacc |
| LH12-doc2-ko-5f2 | gagtagatgccgaccgggaaccagttaacgatggtggtaataaaacaat |
| LH130-AD-SmDOC2-r | tatcgatgcccacccgggtgtcatgaaccaagctcaagagtcaac |
| LH131-BD-SmDOC2-f | gcatatggccatggaggccgaattcatgatggcctccgcgacc |
| LH132-BD-SmDOC2-r | gcaggtcgacggatccccggtcatgaaccaagctcaagagtcaac |
| LH133-AD-HAM5-f | tatggccatggaggccagtgaattcatgtcggtccccggacac |
| LH134-AD-HAM5-r | tatcgatgcccacccgggtgtcagatcatctcactatgatgcaatcc |
| LH135-BD-HAM5-f | gcatatggccatggaggccgaattcatgtcggtccccggacac |
| LH136-BD-HAM5-r | gcaggtcgacggatccccggtcagatcatctcactatgatgcaatcc |
| LH137-MEK2-t2-int-r | gaatctcacctgaagtgggtgtaggttcatcgc |
| LH138-MEK2-t2-int-f | acccacttcaggtgagattcccattctg |
| LH139-AD-MEK2-t2-r | tatcgatgcccacccgggtgttaaaatcggccggttgg |
| LH13-doc2-ko-5r2 | cggataacaatttcacacaggaaacagctcatacctcagatcggcacat |
| LH140-BD-MEK2-t2-r | gcaggtcgacggatccccggttaaaatcggccggttgg |
| LH15-doc1-v5f | gcgatgtactccaagggtgg |
| LH16-doc1-v3r | cgatggatcttgccgttcct |
| LH18-doc1-vORF-3f | agcaggtcgtgtcaccattt |
| LH19-doc2-v5f | gcgacggcttcttcagtgat |
| LH20-doc2-v3r | gtgtacttgccacggacctt |
| LH21-doc2-vORF-3f | tgttgatggggagagtgtgc |
| LH24-so-doc1-3f | aagaaggcggggaaaacagg |
| LH25-so-doc2-5r2 | ctgcaccaagctgatggatg |
| LH26-so-doc2-5f2 | aggcgaaaagagtccacgg |
| LH27-doc1-vORF-f | tcagggtcatcgtcctccaa |
| LH28-doc1-vORF-r | tggtaggagcggaattgtgg |
| LH30-doc2-vORF-r | accaagctcaagagtcaacg |
| LH31-so-doc1-ORF-f | cggtgatacggacaggaacc |
| LH32-so-doc1-ORF-r | ataggtgccactctcgttgg |
| LH33-doc2-vORF-f2 | acgaagccaaggtcaaggag |
| LH36-HR-doc2TbID3f | cgctctacatgagcatgccctgcccctgatgacgatggaatgggaaagg |
| LH37-HR-T-pTrpC-r | agcccaaaaaatgctccttcaatatcagttcgagtggagatgtggagtg |
| LH38-HR-doc2-nat-f | actgatattgaaggagcattttttgggcttg |
| LH41-doc2-HR-TurboID | acagtccaaacaactccgct |
| LH85-doc1-5r-BglII-2 | tgcttaagatcttgtttcccgttccacaaaggt |

| | |
|---|---|
| LH86-ccg1-f-EcoRI-2 | taagcagaattctagaaggagcagtccatctgc |
| LH88-HR5'doc1-f | cgacggtatcgataagcttgatatcgcggccgcacatcactccgttaaccaccc |
| LH89-HR5'doc1-r | acccatgaattctgtttcccgttccacaaaggttg |
| LH90-HRdoc1-ORF-f | ggaacgggaaacagaattcatgggtagcagcggcaagggc |
| LH91-HRdoc1-ORF-r | gcttcctcctcctccggatcctccgccgccagcaataggcaaatccatacccccc |
| LH92-HRd1-tRFP-1 | ggcggcggaggatccggaggaggaggaagcgtgtctaagggcgaagagctg |
| LH93-HRTtrpC-nat | ttcaatatcagttcgagtggagatgtggagtgg |
| LH94-HRnat-TtrpC | catctccactcgaactgatattgaaggagcatttttgggcttg |
| LH95-HRd1-nat-3' | cgaaatgagtttgtcaggggcagggcatgctcatg |
| LH96-HRd13'-f | ccctgcccctgacaaactcatttcgagcgattcatgaac |
| LH97-HRd13'-r | caaaagctggagctccaccgcggtgtcagacgcgctgcaattgcac |
| LH98-HRd1-Tb | ggcggcggaggatccggaggaggaggaagcaaggacaacaccgtccccctc |
| MAK2KO1 | ctcctgtttattcctccatcagct |
| MAK2KO2 | cctcctcgagagcgaacacatcat |
| nat-1r | tcaggggcagggcatgctca |
| pro11-21 | aagcgcgcttgccagtcgctgc |
| Pro115r_wohph | agttgtgcggttgtcgttggtcgaagc |
| pro11-KOr | acgatcagcctcggaaagaccgc |
| pRS_seq_F | ggcctcttcgctattacgccag |
| pRS_seq_R | cactcattaggcaccccagg |
| sci_f | tcgcaagtcatgtacaccat |
| sci1_vf | aggtcagctcccaacagcaagt |
| sci1_vr | ttgggatcggttcgtggaga |
| Seq_tBioID_rev | gtctggatgtgcttgttgatg |
| SmtBioID-L-f-2 | atgggcggcggcggcagcggcggc |
| sosci1_f | ctggtaggcacctagcccac |
| sosci1_r | gccgtggatgaatggagatc |
| tC1 | caccgcctggacgactaaacc |
| tRFP-f | acgtcgagcagcacgaggtg |
| TtrpC_pRS_r | gcggataacaatttcacacaggaaacagctcgagtggagatgtggagtgg |

**S3 Table. Strains used in this study.**

| Strain | Genotype | Reference |
|---|---|---|
| ***Sordaria macrospora*** | | |
| Δpro11 | *Δpro11:hyg^R^*, ssi, sterile | Bloemendal et al. (2012) |
| Δpro11Δsci1::sci1-TurboID$^{ect}$ | cross of Δsci1::sci1-TurboID$^{ect}$ with Δpro11, ssi, sterile, *hyg*$^R$, *nat*$^R$ | This study |
| Δsci1 | *Δsci1::hyg*$^R$, ssi, sterile | Reschka et al. (2018) |
| Δsci1::sci1-TurboID$^{ect}$ | ectopic integration of *sci1-TurboID* (native *sci1*-promoter) into Δsci1, ssi, fertile, *hyg*$^R$, *nat*$^R$ | Hollstein et al. (2022) |
| fus1-1 | mutation in the *Trihydroxynaphthalene reductase* gene (*SMAC_05650*), light brown ascospores, ssi, fertile | Nowrousian et al. (2012) |
| Smdoc1::Smdoc1-TagRFP-T | integration of *Smdoc1-TagRFP-T* at the native *Smdoc1* locus, ssi, fertile, *Smdoc1-TagRFP-T::nat*$^R$ | This study |
| Smdoc1::Smdoc1-TurboID | integration of *Smdoc1-TurboID* at the native *Smdoc1* locus, ssi, fertile, *Smdoc1-TurboID::nat*$^R$ | This study |
| Smdoc2::Smdoc2-egfp | integration of *Smdoc2-egfp* at the native *Smdoc2* locus, ssi, fertile, *Smdoc2-egfp::nat*$^R$ | This study |
| Smdoc2::Smdoc2-TurboID | integration of *Smdoc2-TurboID* at the native *Smdoc2* locus, ssi, fertile, *Smdoc2-TurboID::nat*$^R$ | This study |
| wild type (wt) | wild type strain, black ascospores, fertile | DSM997, DSMZ |
| wt::egfp-TurboID$^{ect}$ | ectopic integration of *egfp-TurboID* into the *S. macrospora* wild type, ssi, fertile, *nat*$^R$ | This study |
| wt:free-TurboID$^{ect}$ | ectopic integration of free unfused *TurboID* (*ccg1* promoter) into the *S. macrospora* wild type, *hyg*$^R$ | Hollstein et al. (2022) |
| Δku70 | *Δku70::nat*$^R$, ssi, fertile | Pöggeler and Kück (2006) |
| Δku80 | *Δku80::hyg*$^R$, ssi, fertile | Groth et al. (2021) |
| Δku80::pccg1-Smdoc1-TagRFP-T$^{ect}$ | ectopic integration of *ccg1::Smdoc1-TagRFP-T::nat*$^R$ into Δku80, ssi, fertile, *hyg*$^R$, *nat*$^R$ | This study |
| Δku80::pccg1-Smdoc2-egfp$^{ect}$ | ectopic integration of *ccg1::Smdoc2-egf*p::natR into Δku80, ssi, fertile, *hyg*$^R$, *nat*$^R$ | This study |
| Δku80::pccg1-Smdoc2-TagRFP-T$^{ect}$ | ectopic integration of *ccg1:Smdoc2-TagRFP-T::natR* into Δku80, ssi, fertile, *hyg*$^R$, *nat*$^R$ | This study |

| | | |
|---|---|---|
| Δku80::Smdoc2-TagRFP-T$^{ect}$ | ectopic integration of *Smdoc2-TagRFP-T::nat$^{R}$* (native 1 kb 5' region) into Δku80, ssi, fertile, *hyg$^{R}$, nat$^{R}$* | This study |
| Δmak2 | marker-less deletion of *mak2*, Δmak2, fus 1-1, brown ascospores, ssi, sterile | Schmidt et al. (2020) |
| Δmak2;Smdoc2::Smdoc2-TurboID | cross of Δmak2 with Smdoc2::Smdoc2-TurboID, expression of *Smdoc2-TurboID* at the native *Smdoc2* locus in the Δmak2 deletion strain background, ssi, sterile, *Smdoc2-TurboID::nat$^{R}$* | This study |
| ΔSmdoc1 | deletion of *SMAC_06903*, *ΔSmdoc1::hyg$^{R}$*, ssi, impaired sexual development | This study |
| ΔSmdoc1::Smdoc1$^{ect}$ | ectopic integration of the *Smdoc1* ORF with 1 kb 5' and 3' flanks, fertile, *hyg$^{R}$, nat$^{R}$* | This study |
| ΔSmdoc1ΔSmdoc2 | deletion of the genomic region expressing *SMAC_0690*3 and *SMAC_06902*, *ΔSmdoc1ΔSmdoc2::hygR*, ssi, fertile | This study |
| ΔSmdoc2 | deletion of *SMAC_06902*, *ΔSmdoc2::hyg$^{R}$*, ssi, impaired sexual development | This study |
| ΔSmdoc2::Smdoc2$^{ect}$ | ectopic integration of the *Smdoc2* ORF with 1 kb 5' and 3' flanks, fertile, *hyg$^{R}$, nat$^{R}$* | This study |
| ***Neurospora crassa*** | | |
| GN18-16 | *mat a, Δdoc-1::hyg$^{R}$* | Heller et al. (2016) and this study |
| GN18-17 | *mat A, Δdoc-1::hyg$^{R}$* | Heller et al. (2016) and this study |
| GN18-19 | *mat A, Δdoc-1Δdoc-2::hyg$^{R}$* | Heller et al. (2016) and this study |
| GN18-26 | *mat a, Δdoc-1Δdoc-2::hyg$^{R}$* | Heller et al. (2016) and this study |
| FGSC 14643 | *mat a, Δdoc-2::hyg$^{R}$* | FGSC |
| FGSC 14644 | *mat A, Δdoc-2::hyg$^{R}$* | FGSC |
| ***Saccharomyces cerevisiae*** | | |
| PJ69-4A | *MATa, trp1-901 leu2-3_112 ura3-52 his3_200 gal4Δ gal8O Δ LYS2::GAL1-HIS3 GAL2-ADE2 met2::GAL7-lacZ* | James et al. (1996) |
| Y187 | *MATα, ura3-52, his3-200,ade2-101, trp1-901, leu2-3, 112, gal4Δ, gal80Δ, met–, URA3::GAL1$_{UAS}$–Gal1$_{TATA}$–LacZ, MEL1* | Harper et al. (1993) |

| | | |
|---|---|---|
| Y2H Gold | *MATa, trp1-901, leu2-3, 112, ura3-52, his3-200, gal4Δ, gal80Δ, LYS2::GAL1$_{UAS}$–Gal1$_{TATA}$–His3, GAL2$_{UAS}$–Gal2$_{TATA}$–Ade2, URA3::MEL1$_{UAS}$–Mel1$_{TATA}$, AUR1-C MEL1* | Nguyen, unpublished obtained from Takara Bio USA (Matchmaker® Gold Yeast Two-Hybrid System) |
| ***Escherichia coli*** | | |
| MACH1™ | F- φ80(lacZ)ΔM15 ΔlacX74 hsdR($r_K^- m_K^+$) ΔrecA1398 endA1 tonA | Invitrogen |

ect, ectopic integration; ssi, single spore isolate; *hyg$^R$*, hygromycin resistance cassette expressing the *hygromycin B phosphotransferase* gene from *E. coli* under control of the constitutive *trpC* promoter from *A. nidulans; nat$^R$,* nourseothricin resistance cassette expressing the *nourseothricin acetyltransferase* gene from *S. noursei* under control of the constitutive *trpC* promoter from *A. nidulans; ccg1,* promoter of the *clock controlled gene 1* of *N. crassa*; *egfp*, gene for green fluorescent protein enhanced green fluorescent protein of *Aequorea victoria*; *TagRFP-T,* gene for red fluorescent protein TagRFP-T of *Entacmaea quadricolor;* ORF, open reading frame; DSMZ, Deutsche Sammlung von Mikroorganismen und Zellkulturen (Leibniz Institut, Braunschweig).

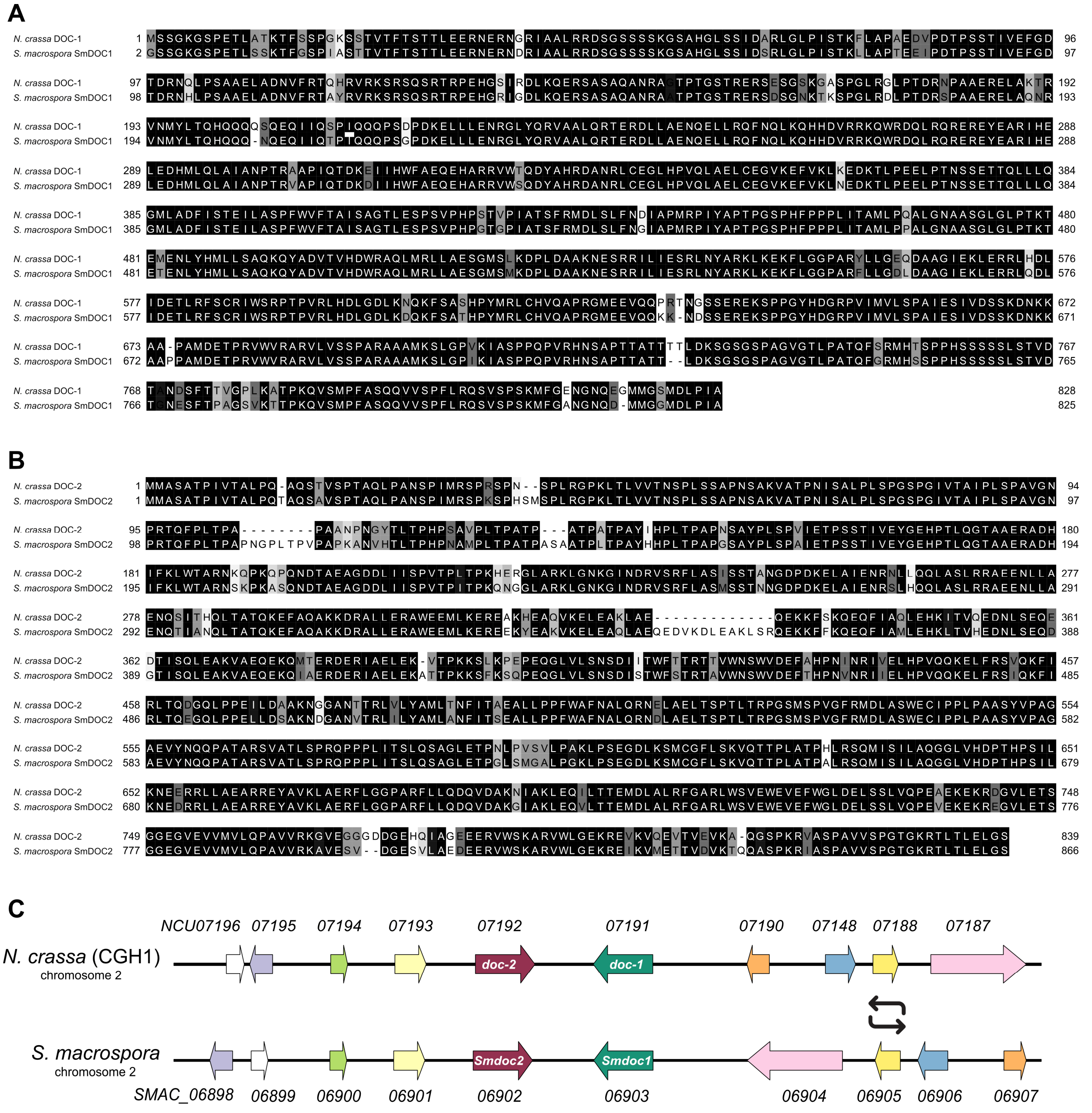

**S1 Fig. Sequence alignments of *S. macrospora* SmDOC1/2 with *N. crassa* DOC-1/2.**

**AB)** Pairwise amino acid sequence alignments of **A)** SmDOC1 (SMAC_06903) with *N. crassa* DOC-1 (NCU07191) and **B)** SmDOC2 (SMAC_06902) with *N. crassa* DOC-2 (NCU07192). Amino acids are colored according to their conservation: regions with low conservation are shaded in white, whereas a high degree of conservation is highlighted in black. Amino acid positions are given on the left and right side of the alignment. The DOC proteins of *S. macrospora* share high sequence identities to *N. crassa*: SmDOC1 = 91.8%; SmDOC2 = 86.3%. This alignment was created with Jalview (Waterhouse et al., 2009). **C)** Genomic map showing the highly syntenic loci encoding the *doc* genes on chromosome two of *S. macrospora* and *N. crassa* strain FGSC 2489, a member of communication group haplotype 1 (CGH1). Genes that are filled in any other color than white are homologs. The region spanning from *NCU07191* up to *NCU07187* is inverted in *S. macrospora*. The *N. crassa doc* locus of CGH1 is based on Heller et al. (2016).

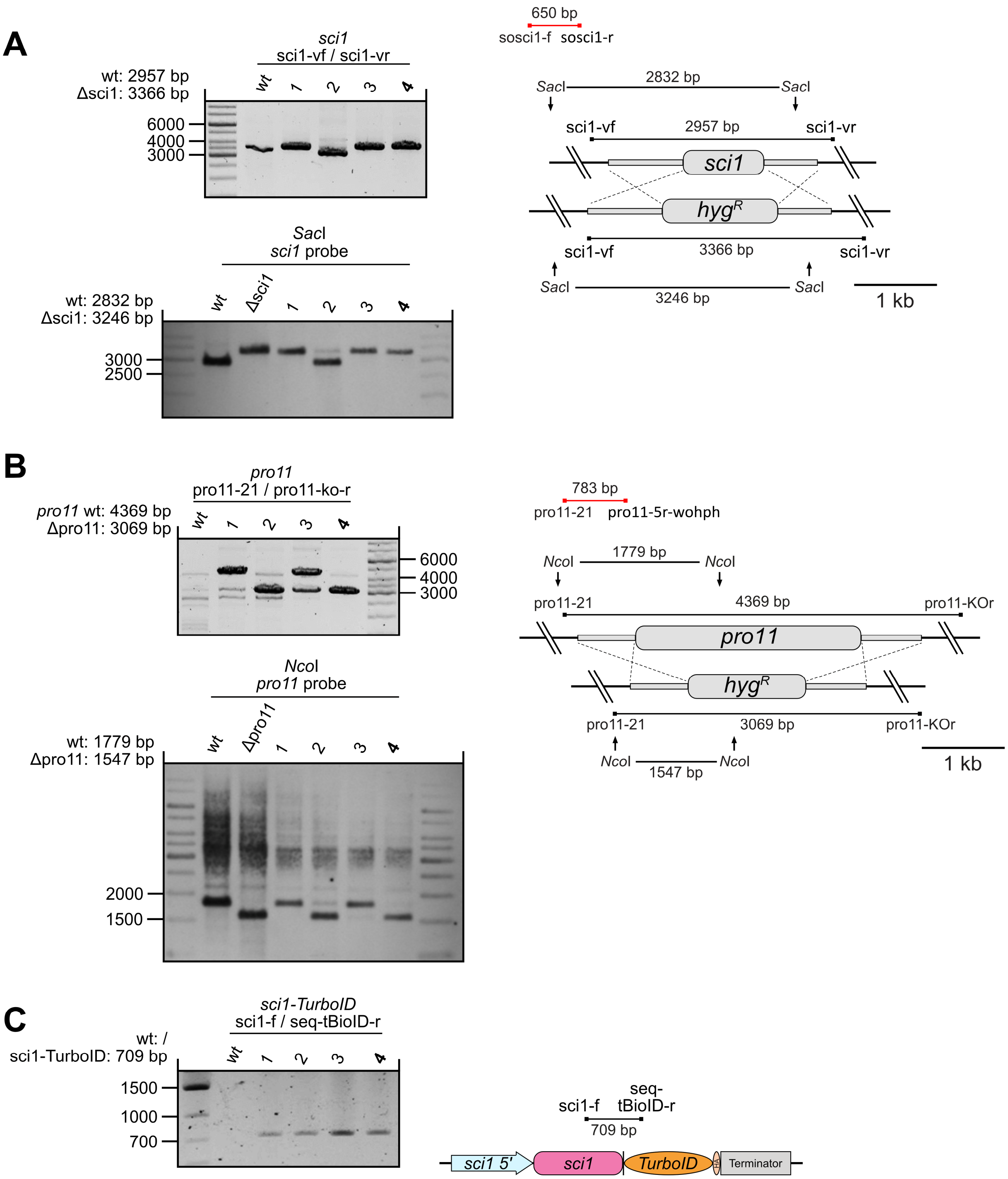

**S2 Fig. Verification of the BioID control strain Δpro11Δsci1::sci1-TurboID$^{ect}$.**
Screening of the progeny from the cross Δpro11 x Δsci1::sci1-TurboID$^{ect}$ for the construction of a BioID control strain. Single spore isolates (1-4) of the putative Δpro11Δsci1::sci1-TurboID$^{ect}$ strain were analyzed by PCR and Southern hybridization experiments to verify **A)** the Δsci1 background, **B)** Δpro11 background and **C)** the ectopic integration of the *sci1-TurboID* fusion gene under control of the native *sci1* promoter (*sci1 5'*). Single spore isolate 4 (printed in bold) was used for further experiments.

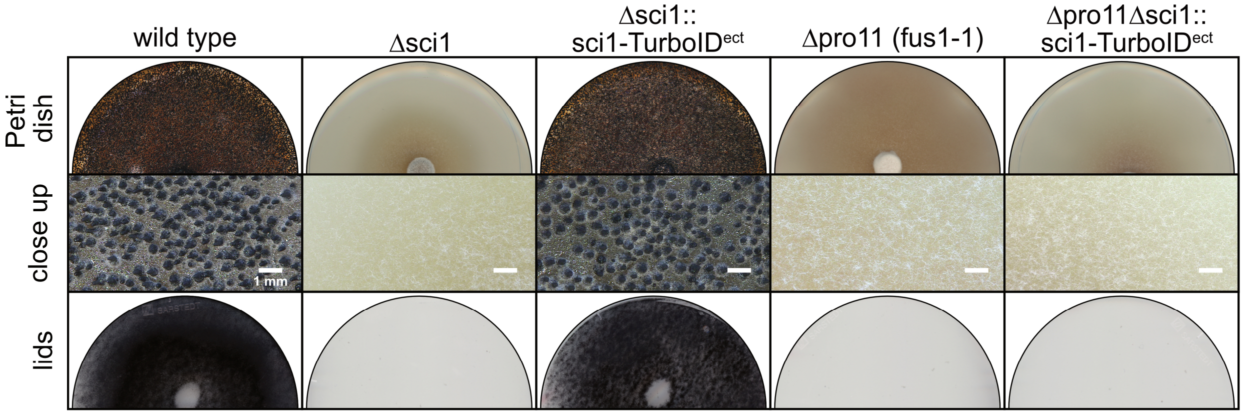

**S3 Fig. Phenotypic characterization of the BioID control Δpro11Δsci1::sci1-TurboID$^{ect}$.**

To analyze the SmSTRIPAK dependent protein environment of SCI1-TurboID, we crossed the Δsci1::sci1-TurboID$^{ect}$ with the Δpro11 deletion strain. The sterile Δpro11 mutant does not produce any fruiting bodies in homokaryotic cultures. The Δpro11 knockout carries the spore color mutant background fus1-1, which results in brown ascospores in crosses. Spores were selected from recombinant perithecia (with brown and black spores) and were verified for the double deletion of *sci1* and *pro11* (S2). For the growth test, the single spore isolates were grown at 27 °C on solid *Sordaria* Westergaard's (SWG) fructification medium. Pictures of the Petri dishes, the close ups and the lids were taken after 14 days of incubation. Once the black ascospores are fully matured inside the fruiting bodies, they are forcefully ejected towards the light source and will stick to the lids of the Petri dishes, thereby staining them black and opaque. The BioID control strain Δpro11Δsci1::sci1-TurboID$^{ect}$ is sterile and does not produce any fruiting bodies.

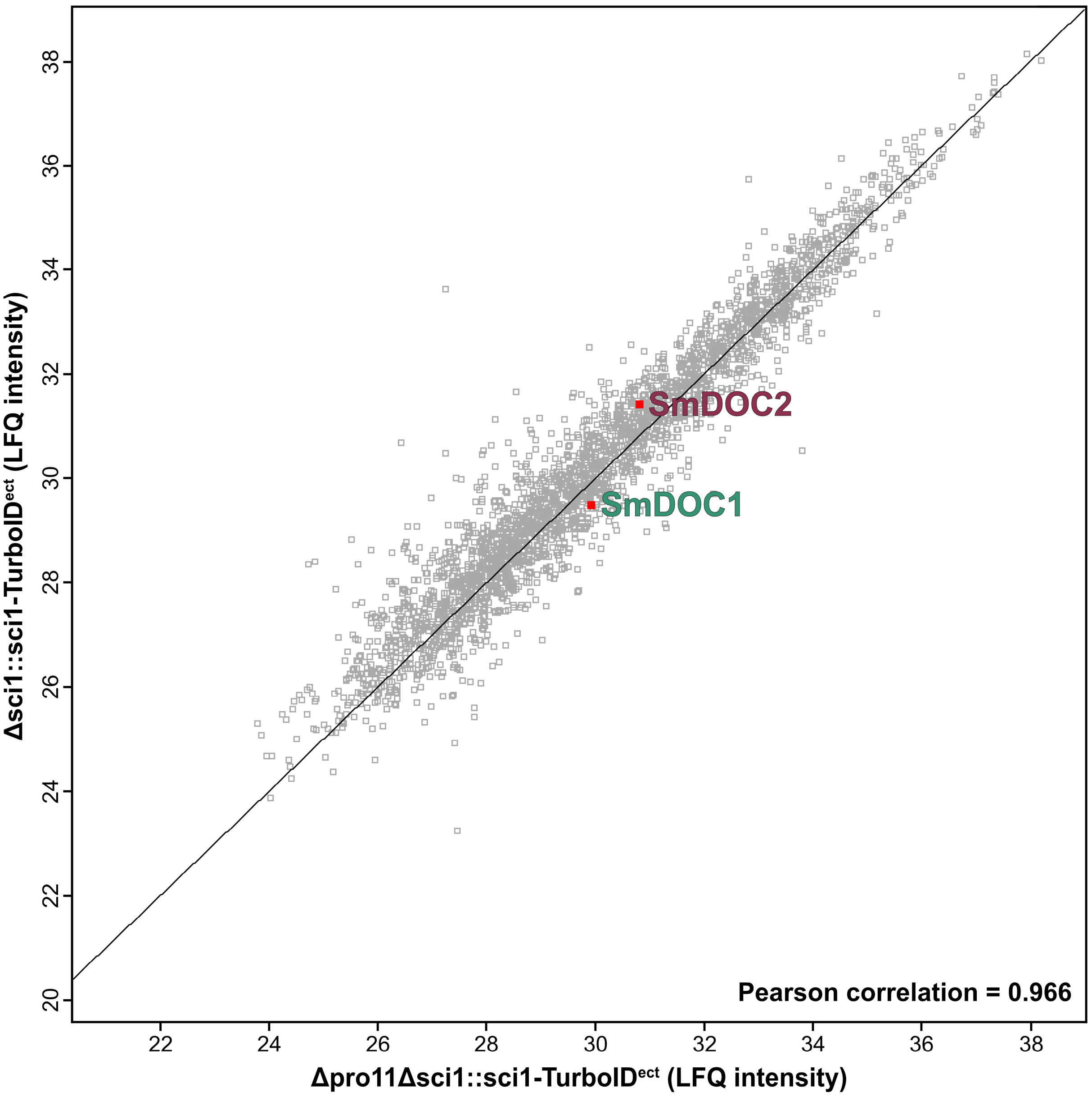

**S4 Fig. Scatterplot analysis of the SCI1-BioID input samples.**

Mass spectrometry analysis of the BioID input samples before biotin affinity purification of the Δsci1::sci1-TurboID strain (fertile) and the Δpro11Δsci1::sci1-TurboID control strain (sterile). A total of 4268 proteins were identified, with 2856 of them being quantified in both strains. The Pearson correlation of this scatterplot is 0.966. This scatterplot plots the LFQ intensity of proteins which were quantified in both strains against each other. The line shows the best fit (1.006079*x). SmDOC2 and SmDOC1 are marked with red rectangles. The protein abundances of SmDOC1 and SmDOC2 do not appear to be regulated in the proteome of the given strains. The LFQ intensities and spectral counts are listed in S4. The full proteomic dataset is listed in S8 Table. LFQ, label free quantification; ect, ectopic integration.

**S4 Table. Selected protein intensities of the proteome analysis of Δpro11Δsci1::sci1-TurboID and Δsci1::sci1-TurboID.**

In this analysis of the input control before biotin affinity purification, SmDOC1 and SmDOC2 were identified with similar LFQ protein intensities in the SCI1-BioID strains Δpro11Δsci1::SCI-TurboID$^{ect}$ (sterile) and Δsci1::sci1-TurboID$^{ect}$ (fertile). This experiment shows that the native protein levels of SmDOC1 and SmDOC2 are not affected by the deletion of *pro11* in the Δpro11Δsci1::SCI-TurboID control strain. The full proteomic dataset is listed in S8 Table. LFQ, label free quantification; ect, ectopic integration; NaN, not a number

| | | LFQ intensities | | MS/MS counts | | |
|---|---|---|---|---|---|---|
| Locus tag | Description | Δpro11Δsci1::sci1-TurboID$^{ect}$ | Δsci1::sci1-TurboID$^{ect}$ | total | Δpro11Δsci1::sci1-TurboID$^{ect}$ | Δsci1::sci1-TurboID$^{ect}$ |
| **SmDOC proteins** | | | | | | |
| SMAC_06903 | SmDOC1 | 29.9 | 29.5 | 38 | 20 | 18 |
| SMAC_06902 | SmDOC2 | 30.8 | 31.4 | 95 | 40 | 55 |
| **Melanin biosynthesis pathway** | | | | | | |
| SMAC_03130 | Polyketide synthase | NaN | 32.4 | 137 | 1 | 136 |
| SMAC_05880 | Tetrahydroxynaphthalene reductase | NaN | 34.7 | 51 | 1 | 50 |
| SMAC_02101 | Scytalone dehydratase | 27.3 | 33.6 | 37 | 4 | 33 |
| SMAC_05650 | Trihydroxynaphthalene reductase | NaN | 33.1 | 28 | 1 | 27 |
| **Developmental proteins** | | | | | | |
| SMAC_08696 | Stage V sporulation protein K | NaN | 31.2 | 99 | 0 | 99 |
| SMAC_08698 | Stage V sporulation protein K | NaN | 29.2 | 42 | 2 | 40 |
| SMAC_12778 | methyltransferase LaeA | NaN | 30.2 | 23 | 2 | 21 |

The plausibility of this proteomic dataset is demonstrated by the absence or downregulation of developmental proteins in the sterile Δpro11 background. RNA seq experiments identified a downregulation of the components of the 1,8 dihydroxynaphthalene (DHN) melanin biosynthesis pathway in *S. macrospora* mutants with impaired fruiting body development. The pathway consists of the four genes for the polyketide synthase (*SMAC_03130*), the tetrahydroxynaphthalene reductase (*SMAC_05880*), the scytalone dehydratase (*SMAC_02101*) and the trihydroxynaphthalene reductase (SMAC_05650). (Dirschnabel et al., 2014; Engh et al., 2010). Consistent with the transcriptome data, all four melanin biosynthesis enzymes show reduced protein intensities in the Δpro11 background in this proteomic dataset. Another major difference between the strains lies in the absence of the proteins SMAC_08696 and SMAC_08698 in the Δpro11 background. Both proteins appear to be homologs of the *N. crassa* Stage V sporulation protein K (NCU09357), which is reported to be highly expressed in late stages of conidiation (Sun et al., 2019). Additionally, the protein with the locus tag SMAC_12778, the homolog of the *N. crassa* methyltransferase LAE-1 (NCU00646), is downregulated in the Δpro11 background. LaeA functions as a global regulator of sexual development and secondary metabolism in *A. nidulans* and its deletion in *N. crassa* results in impaired fruiting body development (Bayram et al., 2019; Cea-Sánchez et al., 2024).

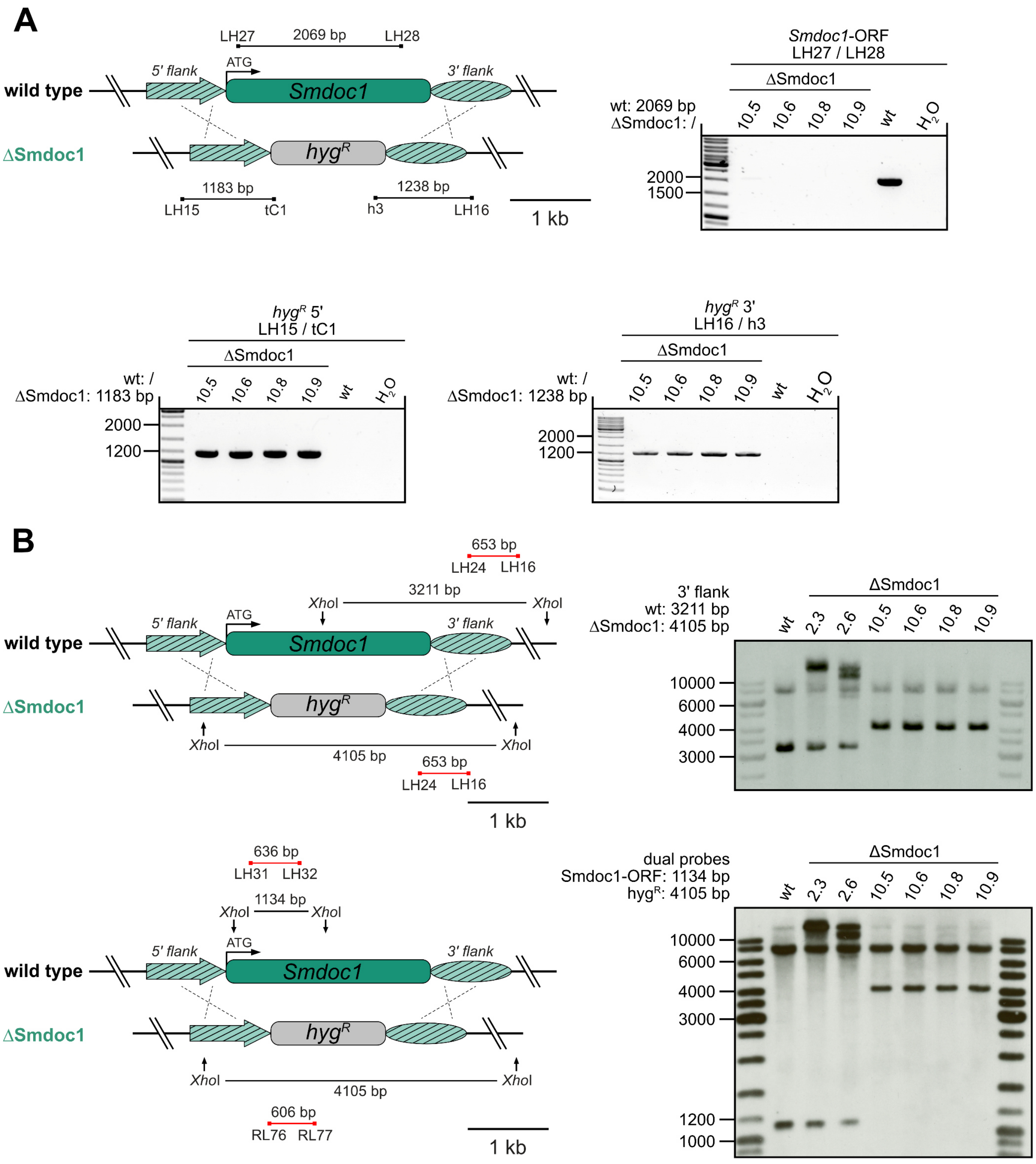

**S5 Fig. Generation and verification of the ΔSmdoc1 deletion strain.**
**A)** Genomic situation of the *Smdoc1* locus (*SMAC_06903)* in the wild type (wt) and the ΔSmdoc1 deletion strain. The amplicons of ΔSmdoc1 PCR verification reactions are indicated by black lines. The absence of the *Smoc1* open reading frame (ORF) was verified with primer pair LH27/LH28 (2069 bp). In single spore isolates 10.5, 10.6, 10.8 and 10.9. The integration of the *hyg*$^R$ cassette at the desired genomic locus was verified using primer pairs LH15/tC1 (1183 bp, *hyg*$^R$ 5' junction) and LH16/h3 (1238 bp, *hyg*$^R$ 3' junction). **B)** Southern hybridization verification of ΔSmdoc1. The probes are indicated by red lines and the restriction enzyme sites are marked by arrows. *hyg*$^R$, hygromycin resistance cassette expressing the *hygromycin B phosphotransferase* gene from *E. coli* under control of the constitutive *trpC* promoter from *A. nidulans*.

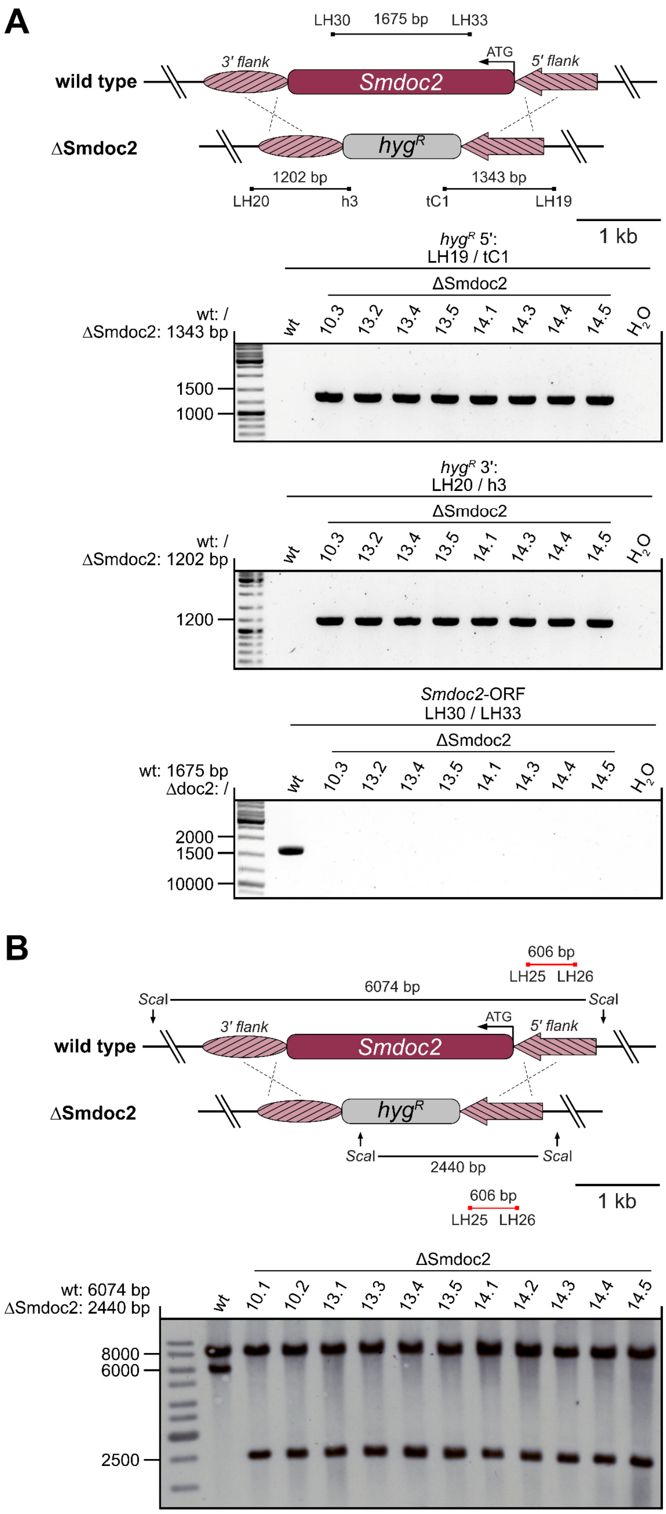

**S6 Fig. Generation and verification of the ΔSmdoc2 deletion strain.**

**A)** Genomic situation of the *Smdoc2* locus (*SMAC_06902*) in the wild type (wt) and the ΔSmdoc2 deletion strain. The amplicons of ΔSmdoc2 PCR verification reactions are indicated by black lines. The integration of the $hyg^R$ cassette at the desired genomic locus of single spore isolates (10.3, 13.2, 13.4, 13.5, 14.1, 14.3, 14.4 and 14.5) was verified using primer pairs LH19/tC1 (1343 bp $hyg^R$ 5' junction) and LH20/h3 (1202 bp, $hyg^R$ 3' junction). The absence of the *Smdoc2* open reading frame (ORF) was verified with primer pair LH30/LH33 (1675 bp). **B)** Southern hybridization verification of ΔSmdoc2. The probe is indicated by a red line and the *Sca*I restriction enzyme sites are marked by arrows. $hyg^R$, hygromycin resistance cassette expressing the *hygromycin B phosphotransferase* gene from *E. coli* under control of the constitutive *trpC* promoter from *A. nidulans*.

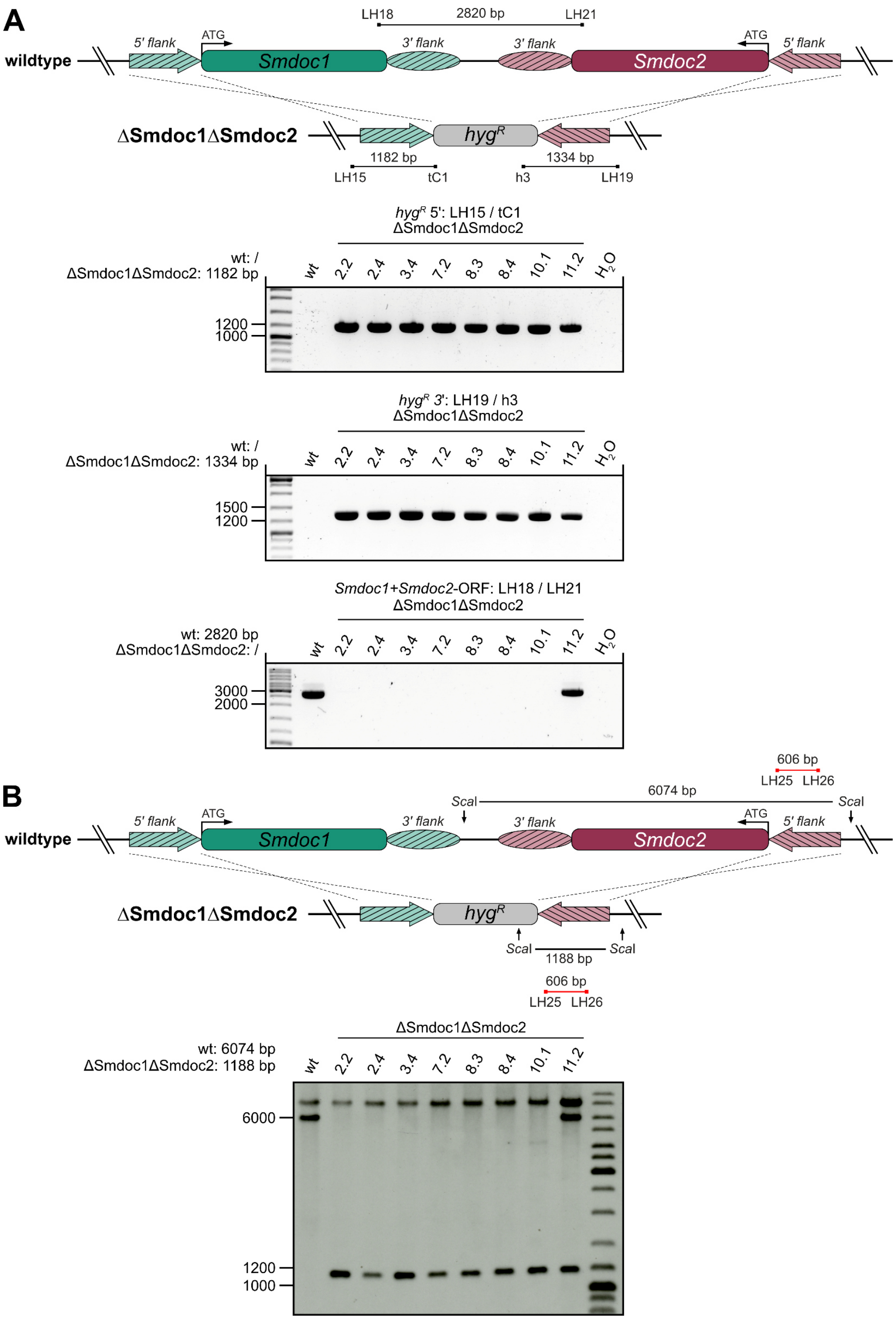

**S7 Fig. Generation and verification of the ΔSmdoc1ΔSmdoc2 double knockout.**
**A)** Genomic situation of the *Smdoc1* (*SMAC_06903*) and *Smdoc2* (*SMAC_06902*) locus in the wild type (wt) and the ΔSmdoc1ΔSmdoc2 double deletion strain. The amplicons of the PCR verification reactions are indicated by black lines. The integration of the $hyg^R$ cassette at the desired genomic locus of the single spore isolates (2.2, 2.4, 3.4, 7.2, 8.3, 8.4, 10.1 and 11.2) was verified using primer pairs LH15/tC1 (1182 bp, $hyg^R$ 5' junction) and LH19/h3 (1334 bp, $hyg^R$ 3' junction). The absence of the *Smdoc1* and *Smdoc2* open reading frames (ORFs) was verified with primer pair LH18/LH21 (2820 bp). **B)** Southern hybridization verification of ΔSmdoc1ΔSmdoc2. The probe is indicated by a red line and the *Sca*I restriction enzyme sites are marked by arrows. The single spore isolate 11.2 appears to be a heterokaryon which carries both, the deletion of *Smdoc1* and *Smdoc2* and the wild-type locus. $hyg^R$, hygromycin resistance cassette expressing the *hygromycin B phosphotransferase* gene from *E. coli* under control of the constitutive *trpC* promoter from *A. nidulans*.

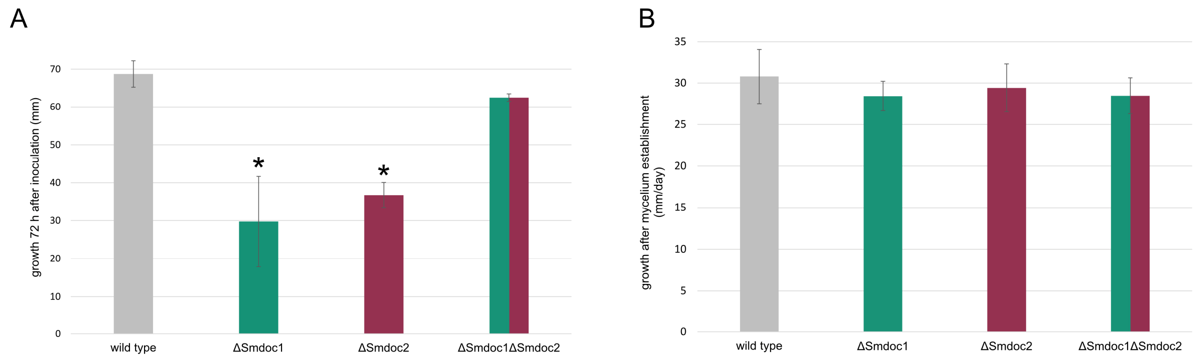

**S8 Fig. Vegetative growth rates of *Smdoc* knockouts.**
The vegetative growth rate was determined in race tubes experiments using the *Sordaria* Westergaard's (SWG) fructification medium over a time frame of 10 days. **A)** Total mycelium length 3 days after incubation of the race tubes. Colony establishment of ΔSmdoc1 and ΔSmdoc2, but not the double knockout, ΔSmdoc1ΔSmdoc2, is impaired. **B)** Average growth rate per day calculated based on growth from day 7 to 10 after inoculation. The error bars show the standard deviation from four replicates. Asterisks indicate significant differences to the wild type according to Student's t-test ($p<0.005$).

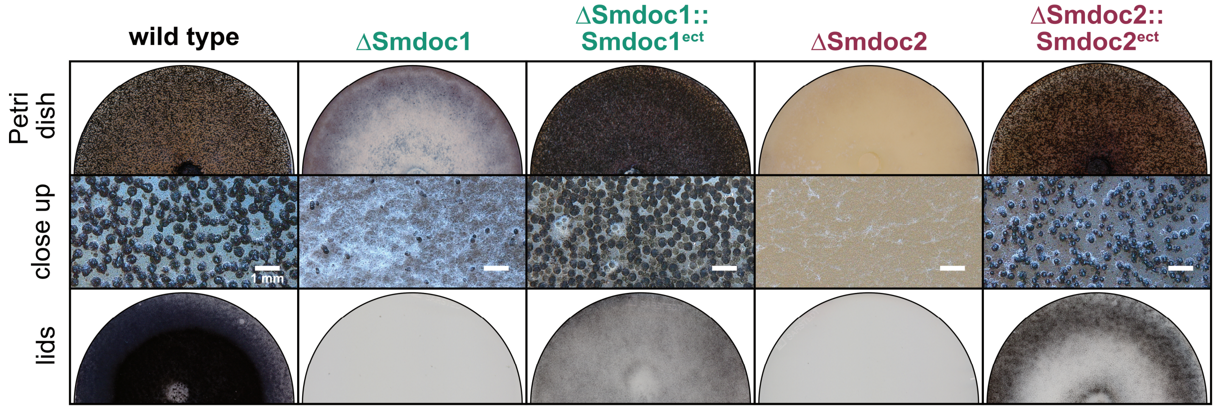

**S9 Fig. Phenotypic characterization of *Smdoc1/2* complementation strains.**
The single spore isolates were grown at 27 °C on solid *Sordaria* Westergaard's (SWG) fructification medium. While the single knockout strains ΔSmdoc1 and ΔSmdoc2 show severe impairment in sexual development, the transformation with the complementation constructs restored fruiting body formation. The complementation constructs were ectopically integrated into the knockout strains and express the open reading frame of *Smdoc1* or *Smdoc2* flanked by 1 kb upstream and downstream regions. The ΔSmdoc1::Smdoc1$^{ect}$ complementation shows increased fruiting body formation when compared to the wild type. Pictures of the Petri dishes, the close ups and the lids were taken after 14 days of incubation. Once the black ascospores are fully matured within the fruiting bodies, they are forcefully ejected towards the light source and will stick to the lids of the Petri dishes.

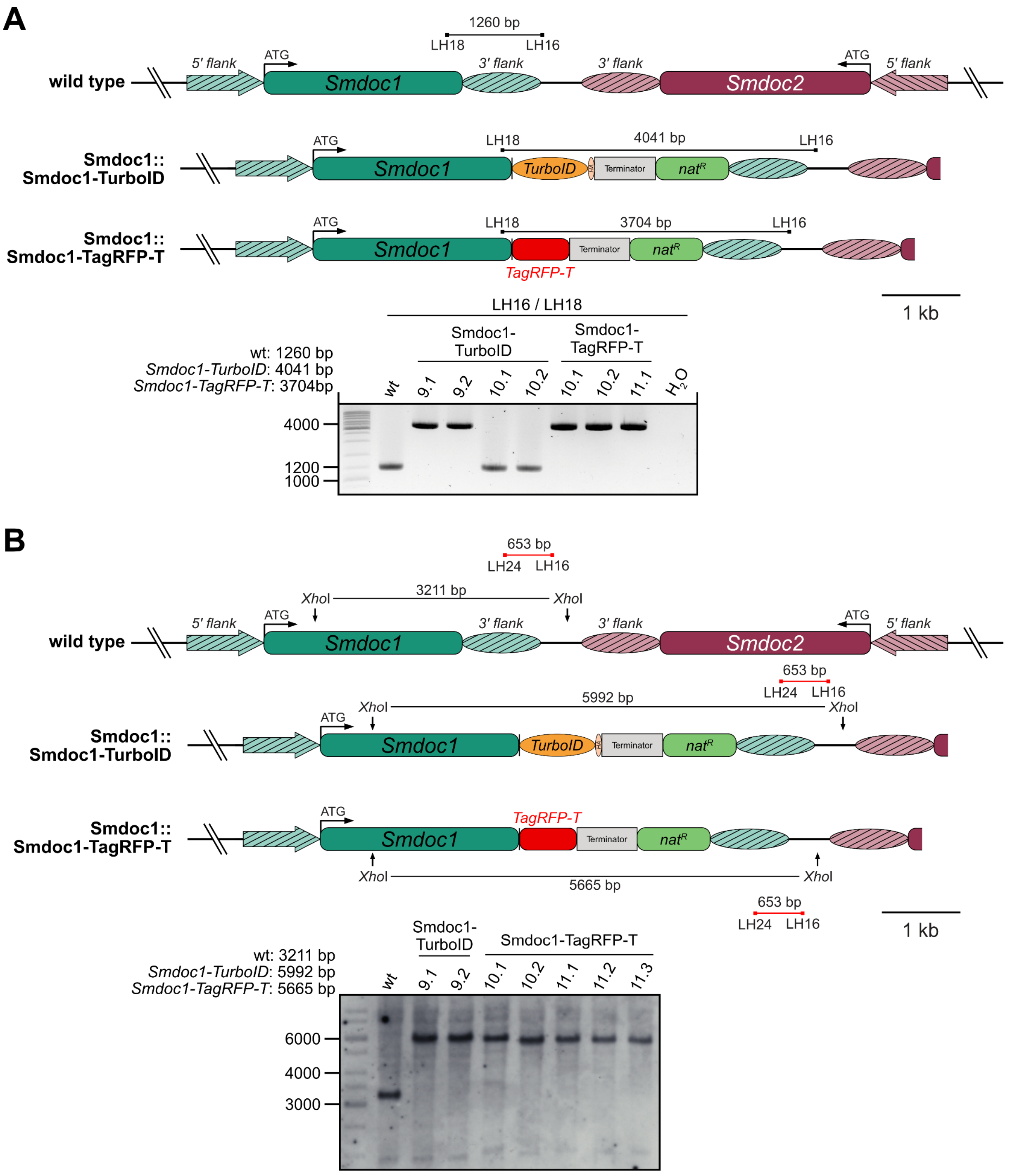

**S10 Fig. *In locus* tagging of *Smdoc1* with *TurboID* and *TagRFP-T*.**

**A)** Genomic situation of the *Smdoc1* (*SMAC_06903*) locus in the wild type and after the integration of *Smdoc1-TurboID* or *Smdoc1-TagRFP-T* fusions. Expression is regulated by the native *Smdoc1* 5' region and the *TtrpC* terminator of the *anthranilate synthase* gene of *A. nidulans* was used for transcription termination. TurboID and TagRFP-T are C-terminally fused to SmDOC1 via a GGGGSGGGGS linker. TurboID is C-terminally tagged with a triple HA-tag. The amplicons of the PCR verification reaction are indicated by black lines. The integration of the fusion at the *Smdoc1* locus was verified using the PCR primer pair LH16/LH18 (*Smdoc1-TurboID* = 4041 bp, *Smdoc1-TagRFP-T* = 3704 bp). **B)** Southern hybridization verification of Smdoc1::Smdoc1-TurboID and Smdoc1::Smdoc1-TagRFP-T. The probe is indicated by a red line and the *Xho*I restriction enzyme sites are marked by arrows. *nat*$^R$, nourseothricin resistance cassette expressing the *nourseothricin acetyltransferase* gene from *S. noursei* under control of the constitutive *trpC* promoter from *A. nidulans*.

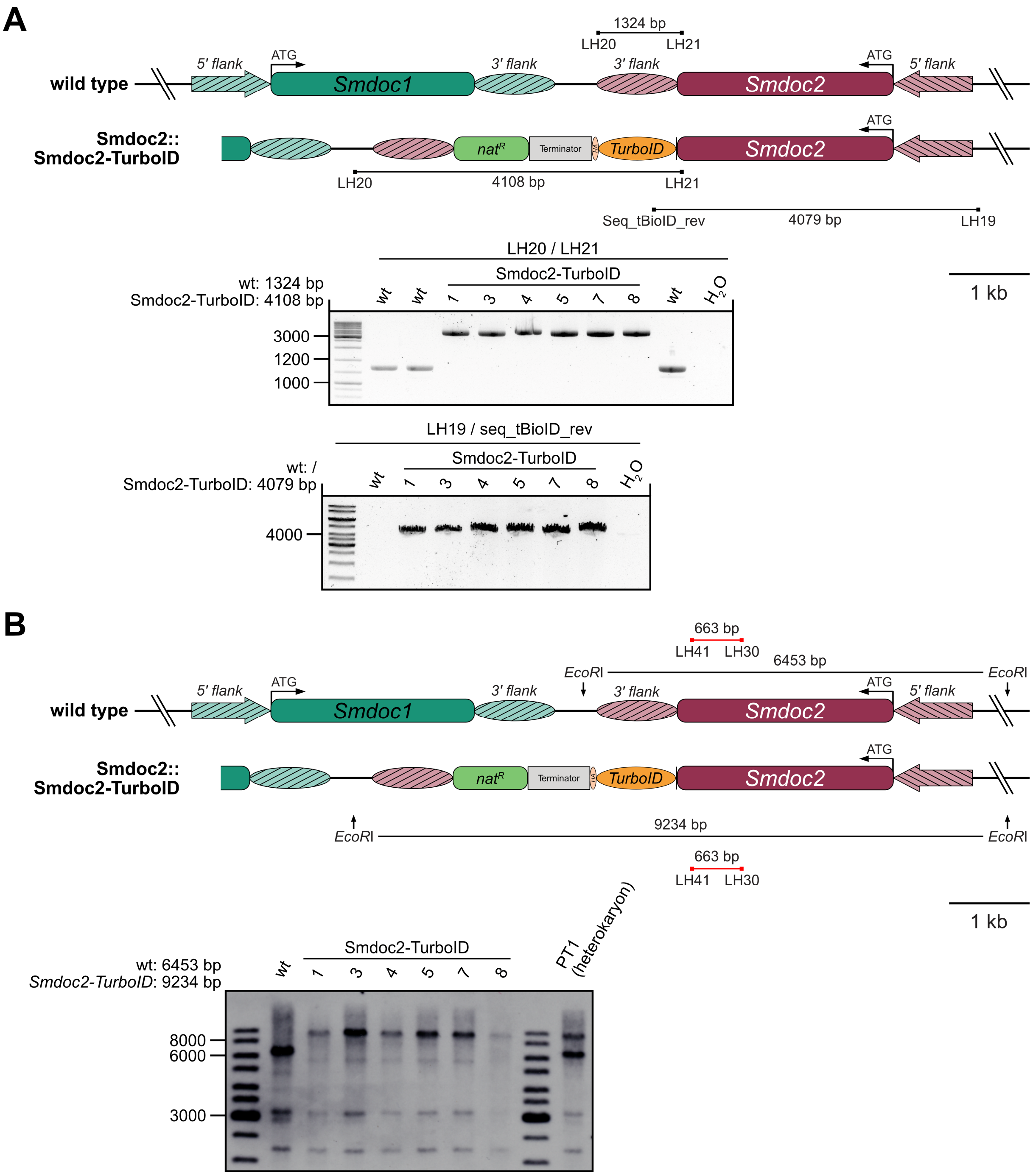

**S11 Fig. *In locus* tagging of *Smdoc2* with *TurboID*.**

**A)** Genomic situation of the *Smdoc2* (*SMAC_06902*) locus in the wild type and after the integration of *Smdoc2-TurboID*. Expression is regulated by the native *Smdoc2* 5' region and the *TtrpC* terminator of the *anthranilate synthase* gene of *A. nidulans* was used for transcription termination. *TurboID* is fused to the C-terminus of *SmDOC2* via a GGGSGGGS linker. Moreover, *TurboID* is C-terminally tagged with a triple HA-tag. The amplicons of the PCR verification reaction are indicated by black lines. The integration of the fusion at the *Smdoc2* locus was verified using the PCR primer pairs LH20/LH21 (4108 bp) and LH19/Seq_tBioID_rev (4079 bp). **B)** Southern hybridization verification of Smdoc2::Smdoc2-TurboID. The probe is indicated by a red line and the *Eco*RI restriction enzyme sites are marked by arrows. The primary transformant (PT) 1 is a heterokaryon and exhibits signals for the wild-type locus and the transformed locus. *nat*$^R$, nourseothricin resistance cassette expressing the *nourseothricin acetyltransferase* gene from *S. noursei* under control of the constitutive *trpC* promoter from *A. nidulans*. PT, primary transformant

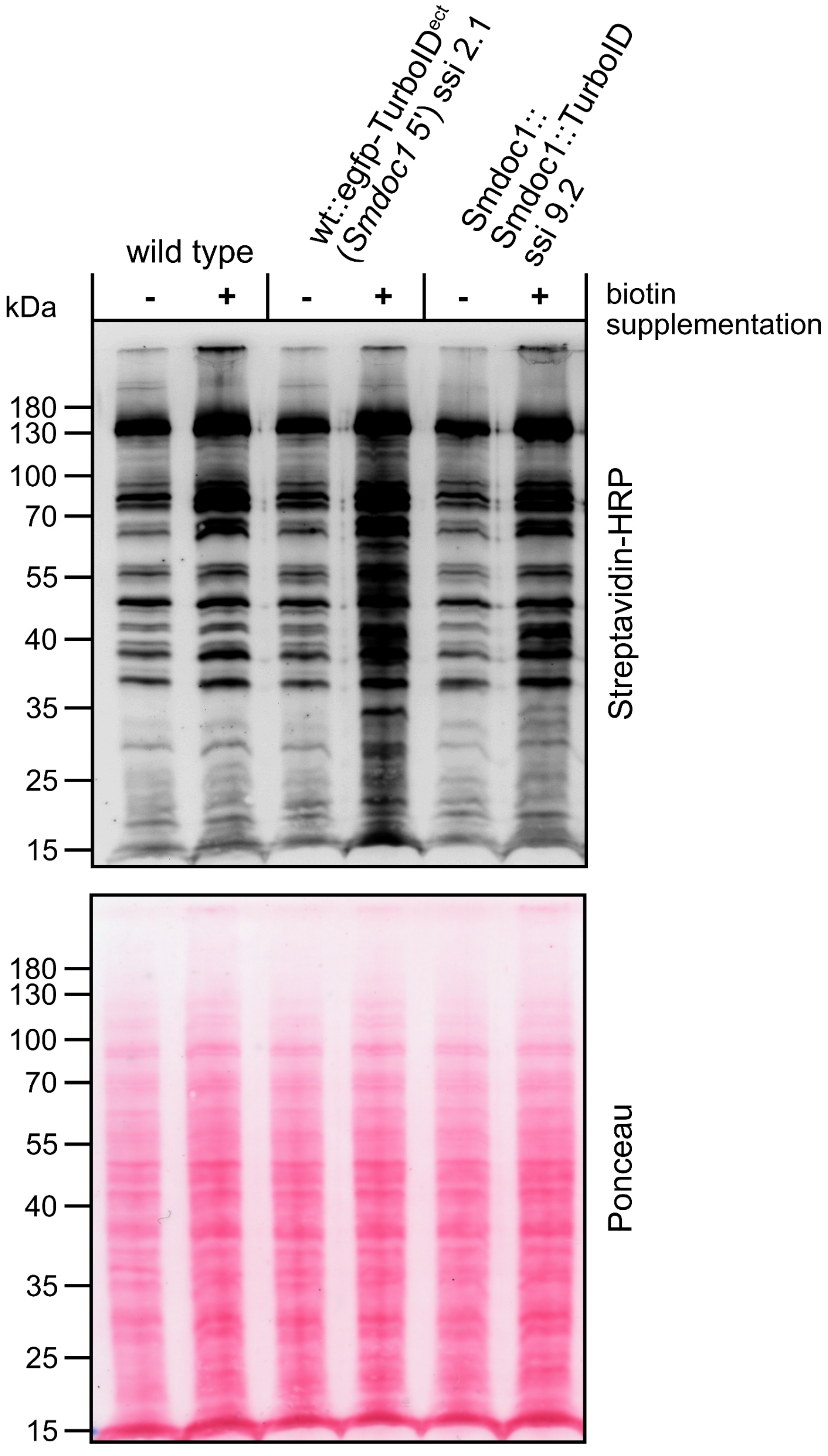

**S12 Fig. Biotinylation activity of the EGFP-TurboID control and the SmDOC1-TurboID fusion protein.** As control in SmDOC BioID experiments, the *egfp-TurboID* fusion gene under control of the *Smdoc1* 1 kb 5' region was ectopically integrated into the *S. macrospora* wild type (wt), resulting in strain wt::egfp-TurboID$^{ect}$. For SmDOC1 BioID experiments, the TurboID ligase was fused to the C-terminus of the SmDOC1 protein via a GGGGSGGGGS linker. The *Smdoc1-TurboID* fusion construct was integrated at the native *Smdoc1* locus (Smdoc1::Smdoc1-TurboID, ssi 9.2). To show the catalytic activity of the ligase, the strains were supplemented with exogenous biotin before harvest of the mycelium for protein extraction. Roughly 50 µg of protein were loaded onto the polyacrylamide gel. Signals were detected with a Streptavidin-HRP conjugate to visualize biotinylated protein. Ponceau S protein staining was used for loading control. ssi, single spore isolate

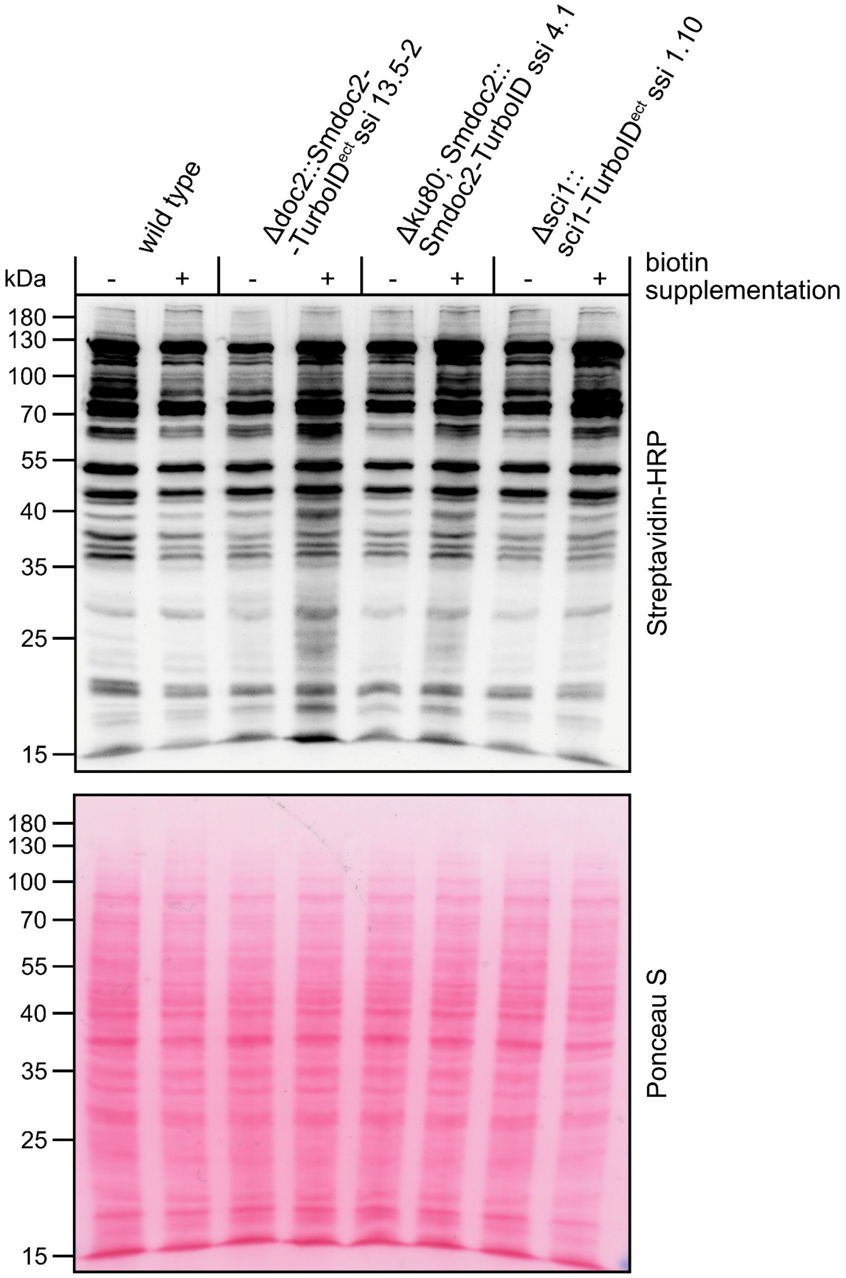

**S13 Fig. Biotinylation activity of the SmDOC2-TurboID fusion protein.**
For SmDOC2 BioID experiments, the TurboID ligase was fused to the C-terminus of the SmDOC2 protein via a GGGGSGGGGS linker. This *Smdoc2-TurboID* fusion construct was either ectopically integrated into the ΔSmdoc2 mutant (ΔSmdoc2::Smdoc2-TurboID$^{ect}$ ssi 13.5-2) or it was integrated at the native *Smdoc2* locus using the non-homologous end joining deficient strain Δku80 (ku80::Smdoc2-TurboID ssi 4.1).To show the catalytic activity of the ligase, the strains were supplemented with exogenous biotin before harvest of the mycelium for protein extraction. Roughly 50 µg of protein were loaded onto the polyacrylamide gel. Signals were detected with a streptavidin-HRP conjugate to visualize biotinylated protein. Ponceau S protein staining was used for loading control. The first lane (wild type without biotin supplementation) shows a slightly increased signal in the loading control. The ectopic integration of *Smdoc2-TurboID* in ΔSmdoc2, the *in locus* integration of *Smdoc2-TurboID* at the native *Smdoc2* locus and the ectopic integration of *sci1-TurboID* into Δsci1 exhibit a substantial increase in overall biotinylation upon biotin supplementation, reaching stronger signals than the wild type. ect, ectopic integration

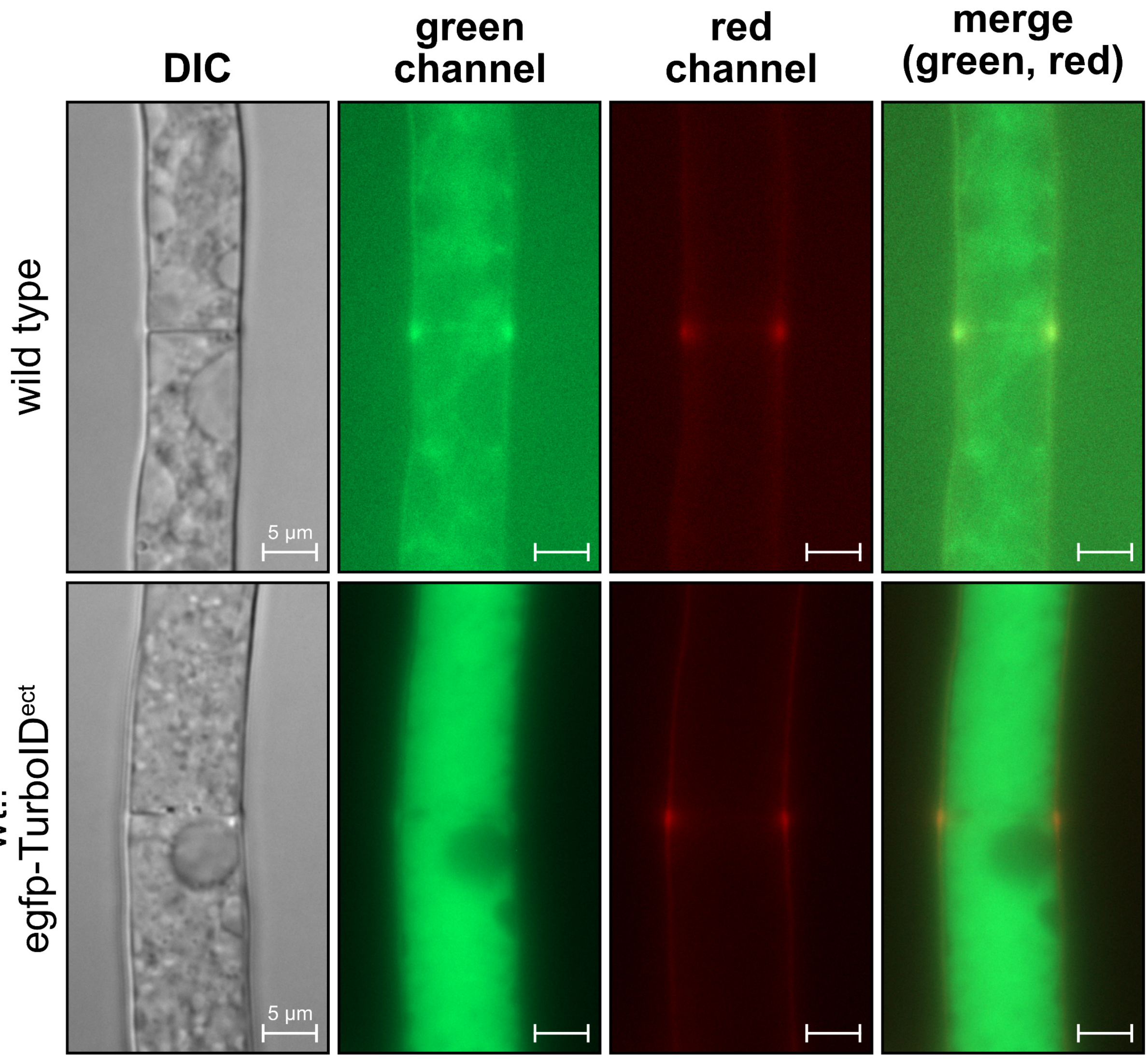

**S14 Fig. Fluorescence microscopy of the BioID control wt::egfp-TurboID$^{ect}$.**
Fluorescence microscopy of *S. macrospora* wild type (wt) and wt with ectopically integrated *egfp-TurboID* under control of the *Smdoc1* 1 kb 5' region (wt::egfp-TurboID$^{ect}$). The exposure settings were identical for the green and the red channel for both strains. The weak autofluorescence in the wild type, especially in the green channel, results in high noise levels in the background. The pronounced cytoplasmic signal of the EGFP-TurboID fusion protein in the green channel results in a less noisy and darker background. The autofluorescence in the red channel is mainly restricted to the septum and the area of the cell wall. A scale bar (5 µm) is indicated.

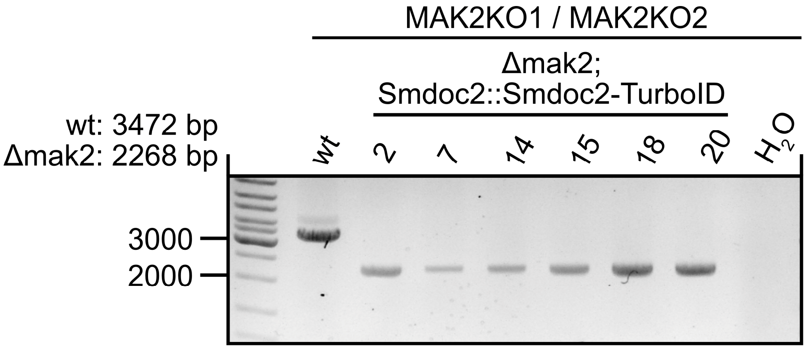

**S15 Fig. PCR verification of the *mak2* deletion in the BioID control strain Δmak2; Smdoc2::Smdoc2-TurboID.**
In order to map the MAK2-dependent proteinaceous environment of SmDOC2, we crossed the Smdoc2::Smdoc2-TurboID (*nat*$^R$) strain with the sterile Δmak2 deletion strain (Schmidt et al., 2020). The Δmak2 deletion strain was generated by replacement of the *mak2* ORF with a resistance cassette (*hyg*$^R$), which was removed using the FLP/*FRT* recombination to create a marker-less Δmak2 deletion strain (Kopke et al., 2010; Schmidt et al., 2020). Spores were isolated from recombinant perithecia of the cross and the sterile single spore isolates (2, 7, 14, 15, 18 and 20) were tested for the deletion of the *mak2* gene using the primer pair MAK2KO1 and MAK2KO2 from Schmidt et al. (2020). *nat*$^R$*,* nourseothricin resistance cassette expressing the *nourseothricin acetyltransferase* gene from *S. noursei* under control of the constitutive *trpC* promoter from *A. nidulans*. wt, wild type

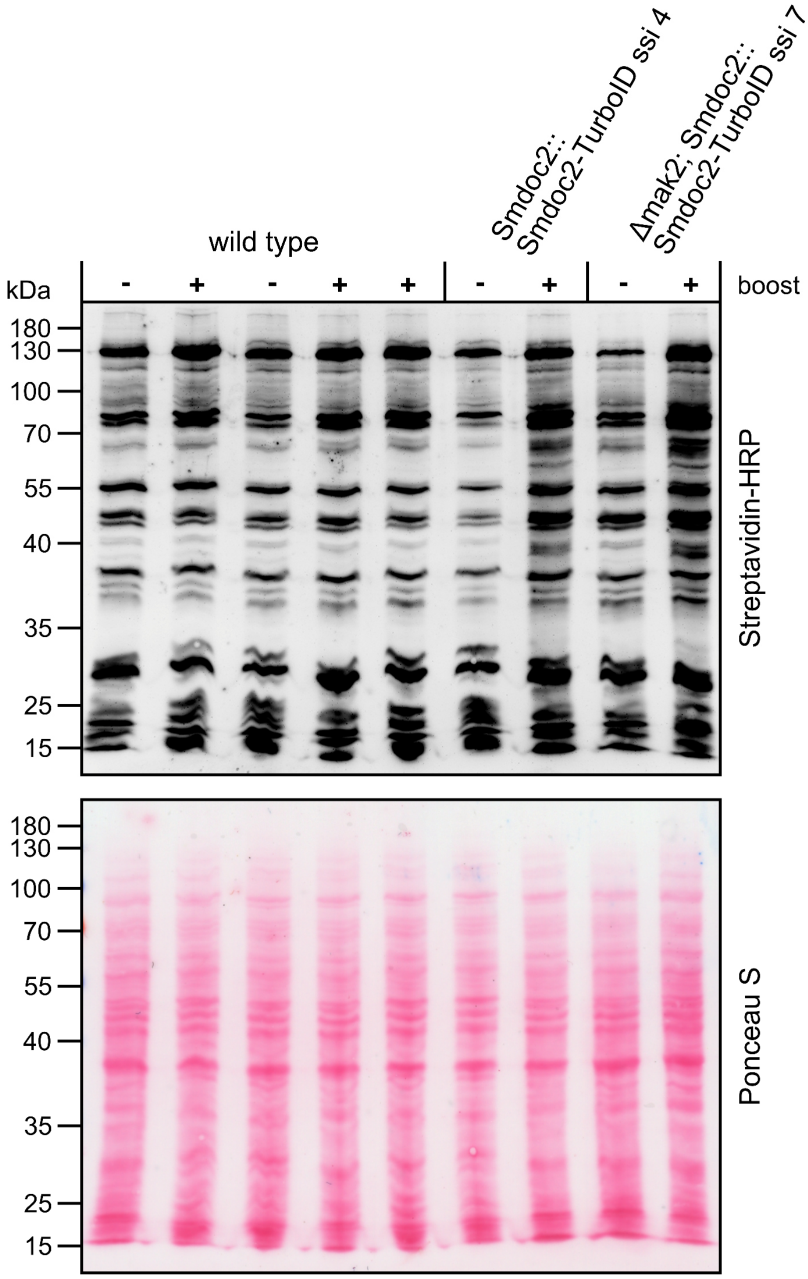

**S16 Fig. SmDOC2-TurboID biotinylation activity.**

For SmDOC2 BioID experiments, the TurboID ligase was fused to the C-terminus of the SmDOC2 protein via a GGGGSGGGGS linker. The *Smdoc2-TurboID* fusion construct was integrated at the native *Smdoc2* locus (Smdoc2::Smdoc2-TurboID ssi 4 and ssi 7). To demonstrate the catalytic activity of the ligase, the strains were supplemented with exogenous biotin before harvest of the mycelium for protein extraction. Roughly 50 µg of protein were loaded onto the polyacrylamide gel. Signals were detected with a Streptavidin-HRP conjugate to visualize biotinylated protein. Ponceau S protein staining was used for loading control.

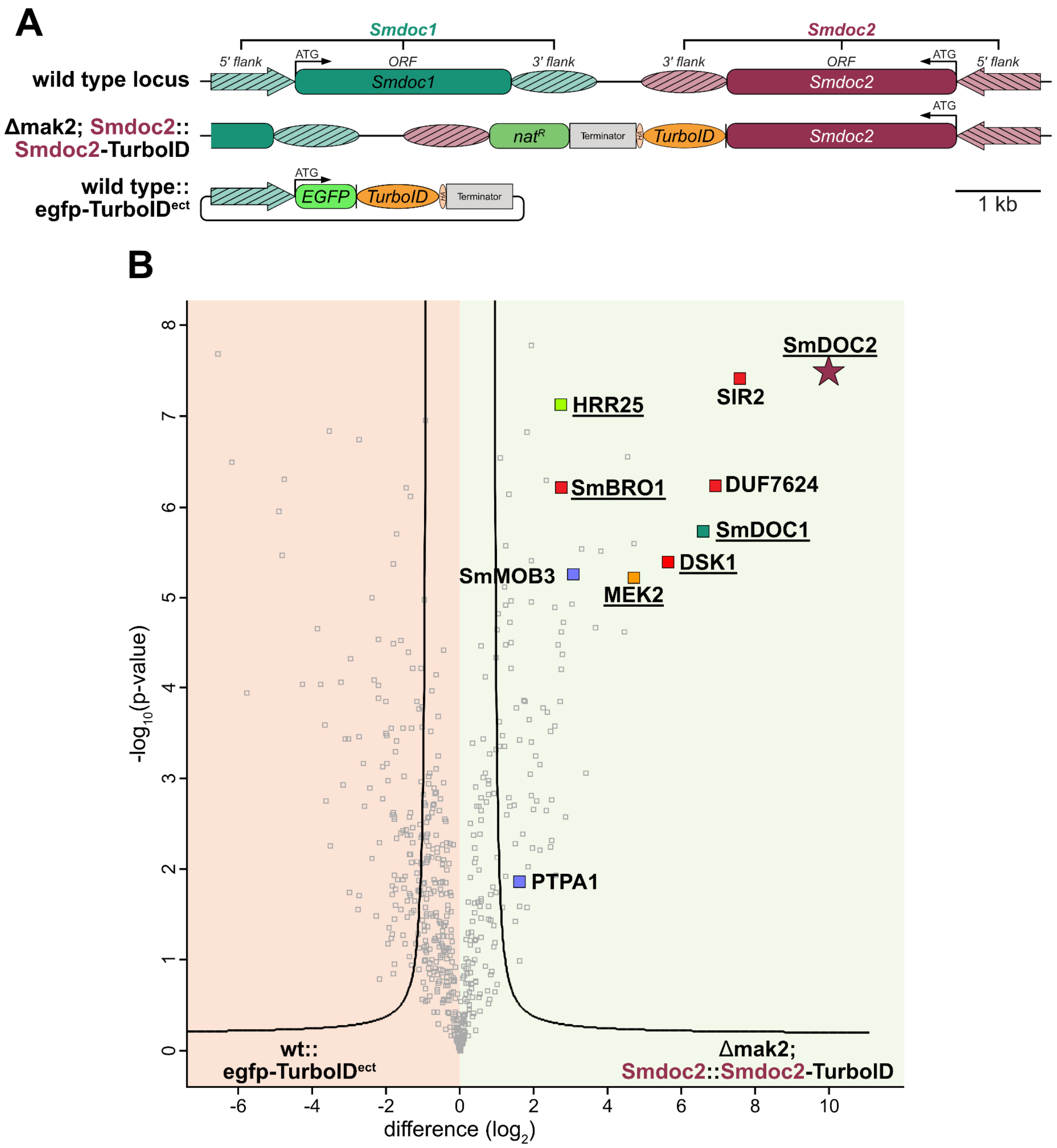

**S17 Fig. Volcano plot analysis of the MAK2-dependent environment of SmDOC2-TurboID.**
Genomic organization of the *Smdoc2* locus in *S. macrospora.* The Δmak2; Smdoc2::Smdoc2-TurboID strain was constructed by crossing the Smdoc2::Smdoc2-TurboID strain with the Δmak2 deletion strain. For relative quantification in BioID experiments, we ectopically integrated an *egfp-TurboID* fusion gene under control of the 1 kb 5' *Smdoc1* region into the *S. macrospora* wild type (wt::egfp-TurboID$^{ect}$). Expression of the *TurboID* ligase is terminated by the terminator of the *anthranilate synthase trpC* gene of *A. nidulans*. TurboID is C-terminally tagged with a triple HA-tag. **B)** The graph plots the difference in label free quantification (LFQ) intensities ($log_2$ transformed) of the control strain on the left and the SmDOC2-TurboID fusion in the Δmak2 deletion strain on the right. The -$log_{10}$(p-value) is plotted on the y axis. Significantly enriched proteins are separated from non-significant proteins by the plotted curve. Proteins that were identified with biotin site information are underlined. Among the significantly enriched proteins, components of the MAK2 pathway are marked in orange, the casein kinase HRR25 is marked in green and the SmSTRIPAK associated protein SmMOB3 and PTPA1 are marked in blue. 4 biological replicates of wt::egfp-TurboID$^{ect}$ and Δmak2; Smdoc2::Smdoc2-TurboID were grown in liquid BMM medium for 4 days at 27 °C under constant light. Statistical parameters of both volcano plots: FDR = 0.01; s0 = 0.1. For details such as MS/MS counts, sequence coverage and other metrics see S11 Table. *nat$^R$*, nourseothricin resistance cassette expressing the *nourseothricin acetyltransferase* gene from *S. noursei* under control of the constitutive *trpC* promoter from *A. nidulans*.

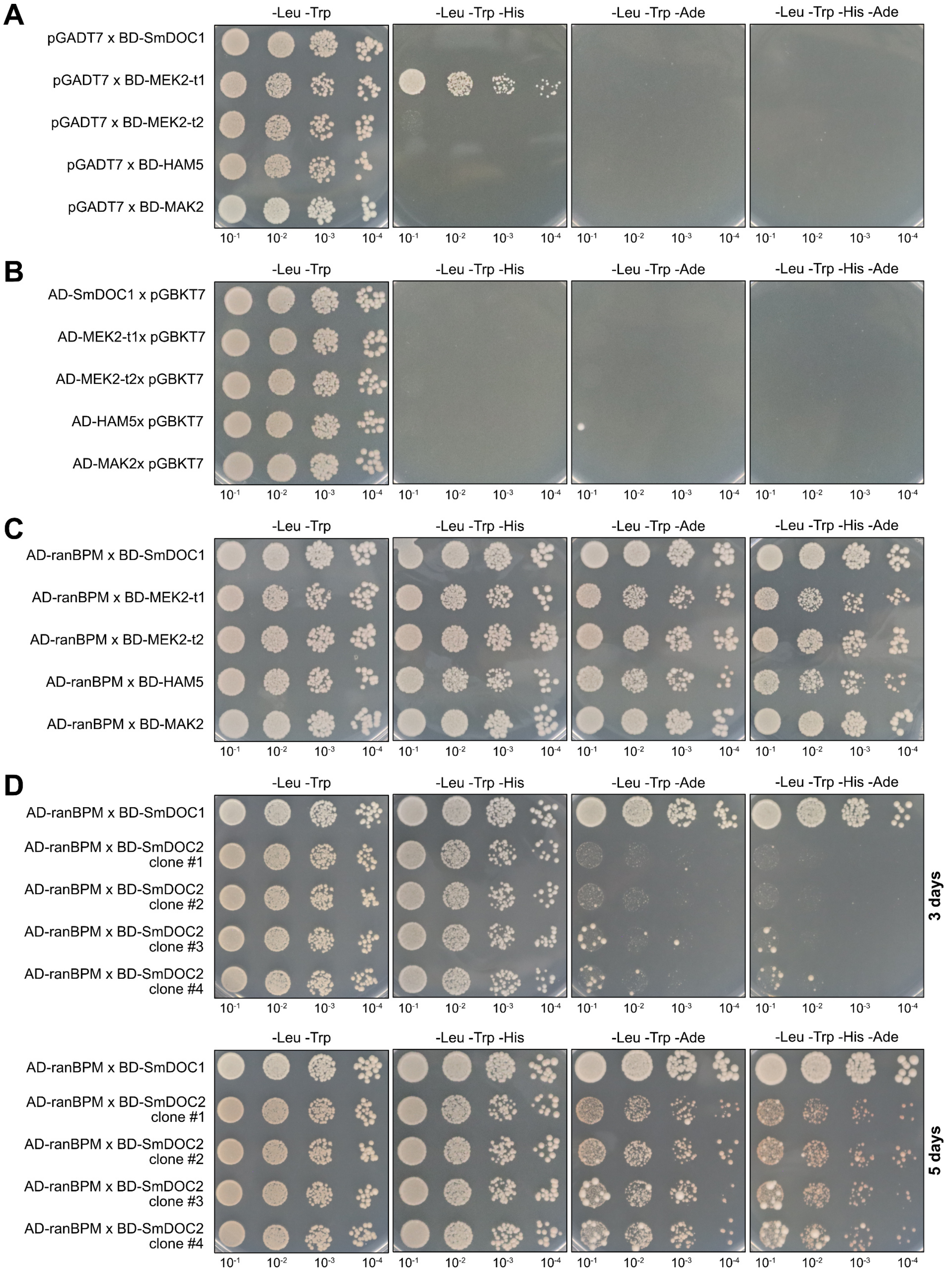

**S18 Fig. Controls for the SmDOC1/2 Y2H experiment.**

**AB**) Negative controls for the interaction studies of SmDOC1 with MEK2-t1, MEK2-t2, HAM5 and MAK2. **A**) Y2H analysis with empty pGADT7 vectors and **B**) Y2H analysis with empty pGBKT7 vectors. **C**) Positive controls using the AD-ranBPM control (Tucker et al., 2009), which directly binds to the Gal4 binding domain of the BD-bait fusion, thereby confirming expression and competency of the BD-SmDOC1/BD-MEK2-t1/BD-MEK2-t2/BD-HAM5/BD-MAK2 fusions. **D**) Positive controls of the AD-ranBPM x BD-SmDOC2 mating, testing different clones All four clones with BD-SmDOC2 showing drastically reduced viability when compared to the AD-ranBPM x BD-SmDOC1 interaction.

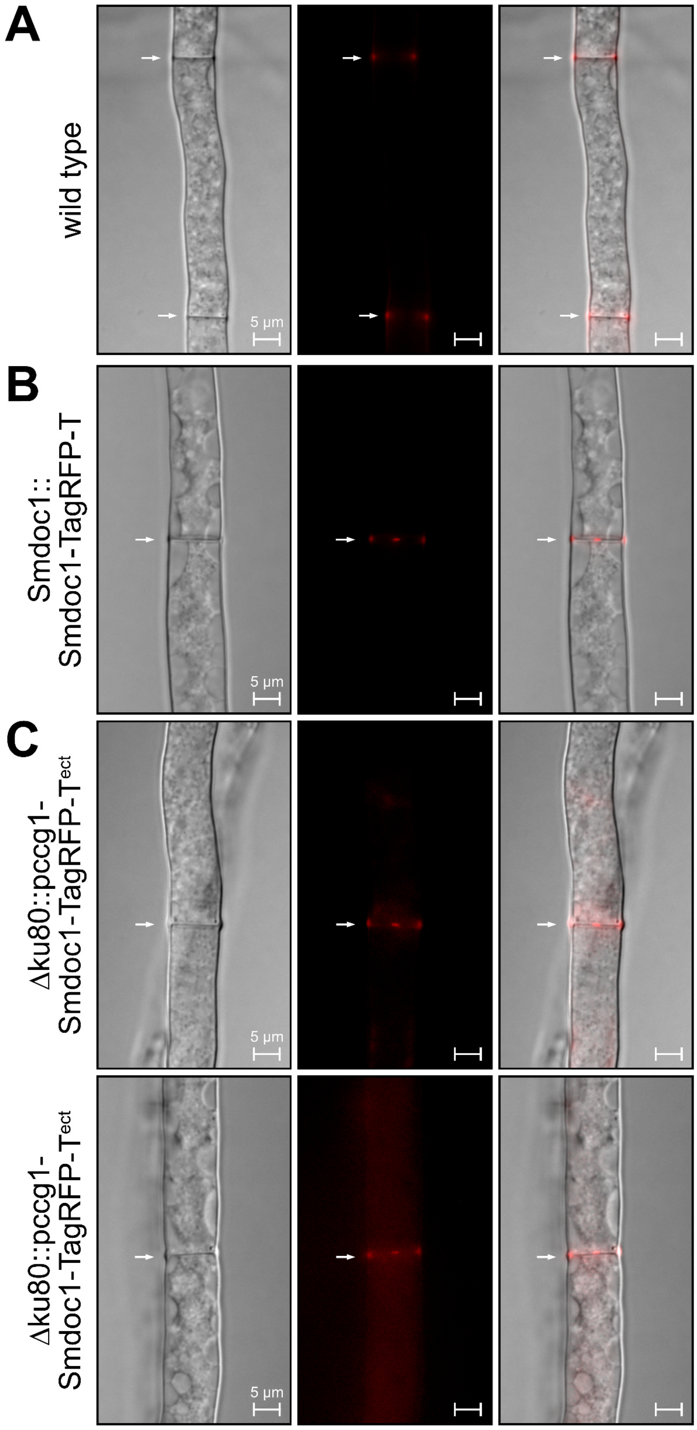

**S19 Fig. Localization studies of SmDOC1-TagRFP-T.**
Fluorescence microscopy of SmDOC1 tagged with TagRFP-T. **A)** Autofluorescence of the *S. macrospora* wild type. **B)** *In locus* integration of *Smdoc1-TagRFP-T* at the *Smdoc1* locus using the native 5' region for expression regulation. **C)** Expression of *Smdoc1-TagRFP-T* under control of the constitutive promoter of the *clock controlled gene 1* (*ccg1*) from *N. crassa* integrated into the *S. macrospora* Δku80 strain, which is used for homologous recombination due to its impaired non-homologous end joining pathway. The fluorescent signal in the red channel did not increase after swapping the native *Smdoc1* 5' region with the constitutive *ccg1* promoter. White arrows indicate septa. A scale bar (5 µm) is indicated. ect, ectopic integration

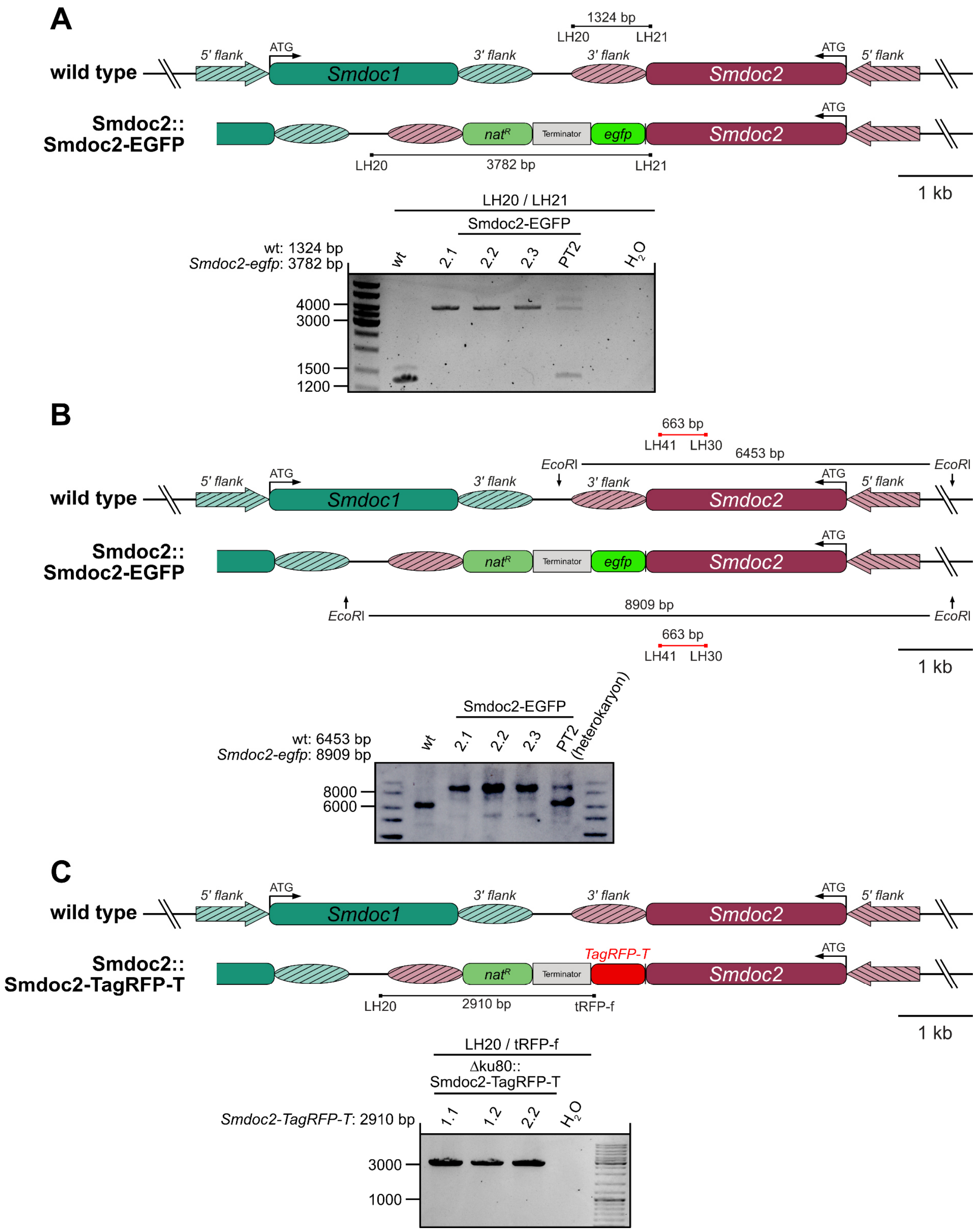

**S20 Fig. *In locus* tagging of *Smdoc2* with *egfp* and *TagRFP-T*.**
**A)** Genomic situation of the *Smdoc2* (*SMAC_06902*) locus in the wild type and after the integration of *Smdoc2-TurboID*. Expression is regulated by the native *Smdoc2* 5' region and the *TtrpC* terminator of the *anthranilate synthase* gene of *A. nidulans* was used for transcription termination. EGFP is fused to the C-terminus of SmDOC2 via a GGGGS linker. The amplicons of the PCR verification reaction are indicated by black lines. The integration of the fusion at the *Smdoc2* locus was verified using the PCR primer pair LH20/LH21 (3782 bp) in the single spore isolates 2.1, 2.2, 2.3. **B)** Southern hybridization verification of Smdoc2::Smdoc2-EGFP. The probe is indicated by a red line and the *Eco*RI restriction enzyme sites are marked by arrows. The primary transformant (PT) 2 is a heterokaryon and exhibits signals for the wild-type locus and the transformed locus. **C)** *TagRFP-T* tagging of the *Smdoc2* gene was performed analogously to the strain construction of Smdoc2::Smdoc2-EGFP. Primer pair LH20/tRFP-f (2910 bp) was used to verify the *in locus* integration in the single spore isolate 1.1, 1.2 and 2.1. *nat*$^R$, nourseothricin resistance cassette expressing the *nourseothricin acetyltransferase* gene from *S. noursei* under control of the constitutive *trpC* promoter from *A. nidulans*. PT, primary transformant

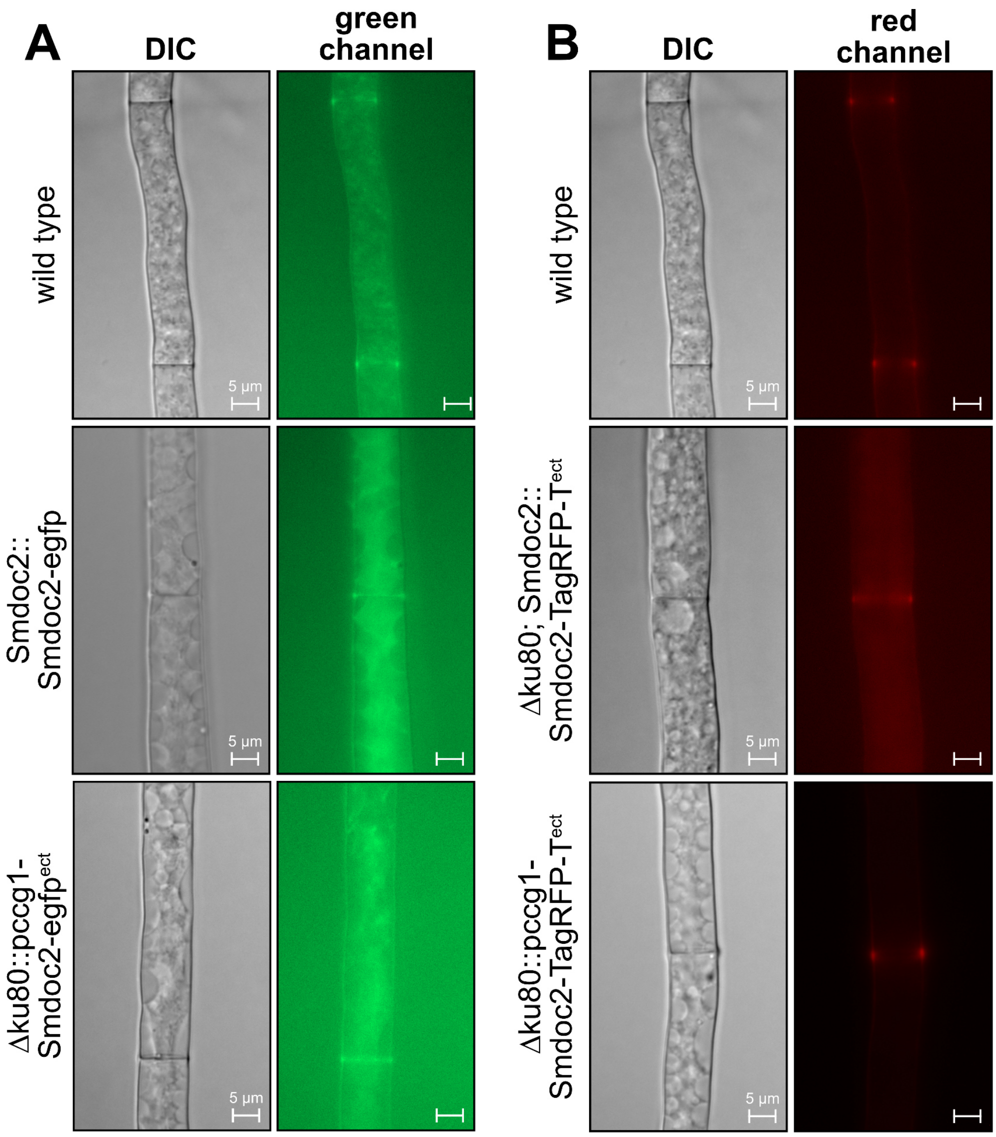

**S21 Fig. Fluorescence microscopy of SmDOC2-EGFP and SmDOC2-TagRFP-T.**
Fluorescence microscopy of *Smdoc2* tagged with **A)** *egfp* and **B)** *TagRFP-T*. **A)** *In locus* integration of *Smdoc2-egfp* at the *Smdoc2* locus using the native 5' region for expression regulation. The construct was transformed into the *S. macrospora* Δku80 strain, which is used for homologous recombination due to its impaired non-homologous end joining pathway. The strain was fully verified for the *in locus* integration of *Smdoc2-egfp* and the absence of the wild-type *Smdoc2* gene by PCR and Southern hybridization. The wild-type *ku80* gene was restored by crossing with the color spore mutant fus 1-1 expressing the wild type *ku80* gene. The fluorescent signal of SmDOC2-EGFP did not surpass the intensity of the autofluorescence observed in the *S. macrospora* wild type. The exchange of the 5' region with the constitutive promoter of the *clock controlled gene 1* (*ccg1*) from *N. crassa* did not result in any changes in fluorescent output. **B)** Since the autofluorescence of the *S. macrospora* wild type is less pronounced in the red channel in fluorescence microscopy, we fused *Smdoc2* to the red fluorescent protein *TagRFP-T*. Expression of this *Smdoc2-TagRFP-T* fusion is either controlled by the 1 kb 5' *Smdoc2* region or the constitutive *ccg1* promoter. However, no fluorescent signal was observed for SmDOC2-TagRFP-T using either promoter for expression regulation. A scale bar (5 µm) is indicated. ect, ectopic integration
